# Vaginal *Lactobacillus* fatty acid response mechanisms reveal a novel strategy for bacterial vaginosis treatment

**DOI:** 10.1101/2023.12.30.573720

**Authors:** Meilin Zhu, Matthew W. Frank, Christopher D. Radka, Sarah Jeanfavre, Megan W. Tse, Julian Avila Pacheco, Kerry Pierce, Amy Deik, Jiawu Xu, Salina Hussain, Fatima Aysha Hussain, Nondumiso Xulu, Nasreen Khan, Vanessa Pillay, Krista L. Dong, Thumbi Ndung’u, Clary B. Clish, Charles O. Rock, Paul C. Blainey, Seth M. Bloom, Douglas S. Kwon

## Abstract

Bacterial vaginosis (BV), a common syndrome characterized by *Lactobacillus*-deficient vaginal microbiota, is associated with adverse health outcomes. BV often recurs after standard antibiotic therapy in part because antibiotics promote microbiota dominance by *Lactobacillus iners* instead of *Lactobacillus crispatus*, which has more beneficial health associations. Strategies to promote *L. crispatus* and inhibit *L. iners* are thus needed. We show that oleic acid (OA) and similar long-chain fatty acids simultaneously inhibit *L. iners* and enhance *L. crispatus* growth. These phenotypes require OA-inducible genes conserved in *L. crispatus* and related species, including an oleate hydratase (*ohyA*) and putative fatty acid efflux pump (*farE*). FarE mediates OA resistance, while OhyA is robustly active in the human vaginal microbiota and sequesters OA in a derivative form that only *ohyA*-harboring organisms can exploit. Finally, OA promotes *L. crispatus* dominance more effectively than antibiotics in an *in vitro* model of BV, suggesting a novel approach for treatment.

## Introduction

Female genital tract (FGT) microbiota composition is linked to numerous adverse health outcomes, such as preterm birth^1^, infertility^2–5^, cervical dysplasia^6–8^, and sexually transmitted infections^9^ including human immunodeficiency virus (HIV) risk^10,11^. Bacterial vaginosis (BV) – a syndrome of the FGT microbiota associated with these adverse outcomes – affects up to 58% of women worldwide^12^ and has clinical features including watery discharge, odor, and mucosal inflammation^13–15^. Microbiologically, BV is characterized by a paucity of lactobacilli and high abundance of diverse obligate anaerobic species^16,17^. By contrast, health-associated FGT microbial communities are typically dominated by *Lactobacillus* species, most notably *Lactobacillus crispatus*, but also including *Lactobacillus gasseri*, *Lactobacillus jensenii*, and *Lactobacillus mulieris*. However, FGT microbial communities dominated by a different common FGT *Lactobacillus* species – *Lactobacillus iners* – have several sub-optimal health associations^6,10,18^, including higher risk of transitioning to BV^19–21^.

Standard first-line BV therapy with the antibiotic metronidazole (MTZ) has partial efficacy, but ≥50% of treated patients experience recurrence within one year^22–24^. One explanation for high recurrence rates is the high probability of MTZ treatment to shift FGT microbiota composition towards dominance by *L. iners*^21,25–28^ instead of more health-associated lactobacilli like *L. crispatus*. Resistance to MTZ among BV-associated bacteria may also impair treatment efficacy and promote recurrence, although the clinical significance of MTZ resistance in BV remains unclear^29–31^. Experimental interventions to improve MTZ efficacy using adjunctive non-antibiotic strategies such as vaginal microbiota transplants and *L. crispatus*-containing intravaginal live biotherapeutic products have shown some promise compared to MTZ alone, but reported recurrence rates from these studies remain high^24,32,33^. To our knowledge, no therapies currently exist to promote *L. crispatus* dominance in the FGT microbiota by selectively inhibiting *L. iners* or enhancing *L. crispatus* growth.

Mammalian mucosal surfaces are rich with LCFAs^34–37^, which serve as critical nutrients and building blocks for bacterial membrane components and other biological processes, but can also display antimicrobial properties^38^. Certain unsaturated long-chain fatty acids (uLCFAs) exert antimicrobial activity against host-adapted Gram positive organisms such as *Staphylococcus aureus*^39,40^. However, uLCFA effects on growth or inhibition of FGT *Lactobacillus* species have not been systematically assessed. *L. iners* has a reduced genome size and metabolic capacity relative to other FGT lactobacilli^41,42^, suggesting feasibility of exploiting metabolic differences between *Lactobacillus* species to selectively target *L. iners*^43^. We hypothesized that long-chain fatty acid (LCFA) metabolism might constitute a metabolic target to differentially modulate FGT lactobacilli and other bacteria.

In this study, we investigated effects of *cis-*9-uLCFAs on FGT *Lactobacillus* species and assessed their potential to selectively modulate FGT microbiota composition. We found that *cis-* 9-uLCFAs selectively inhibited *L. iners* while simultaneously robustly promoting growth of non-*iners* FGT lactobacilli. Guided by transcriptomic and genomic analyses, we identified a putative fatty acid efflux pump (*farE*) and an oleate hydratase enzyme (*ohyA*) that were upregulated by *cis-*9-uLCFAs in non-*iners* FGT *Lactobacillus* species. These genes were genomically conserved in non*-iners* lactobacilli but completely absent in *L. iners*, mirroring the observed growth phenotypes. Using a genetically tractable strain of *L. gasseri*, we showed that *farE* is required for both the *cis-*9-uLCFA resistance and growth enhancement phenotypes. Biochemical characterization of the *Lactobacillus* OhyA orthologs revealed them to be robustly active *in vivo* in the human FGT microbiota and demonstrated that non-*iners* FGT lactobacilli can utilize OhyA to bioconvert and sequester exogenous OA for use in phospholipid synthesis. Taken together, our data support that *ohyA* and *farE* confer non-*iners* lactobacilli a growth advantage in *cis*-9-uLCFA-rich environments. We leverage this discovery to selectively modulate *in vitro* BV-like bacterial communities towards *L. crispatus* dominance using *cis*-9-uLCFA treatment alone or in combination with MTZ. Collectively, this work identifies important species-level differences in FGT *Lactobacillus* metabolism, elucidates the mechanisms underlying those differences, and provides evidence for a novel metabolism-targeted intervention to improve female genital and reproductive health.

## Results

### *cis*-9-uLCFAs selectively inhibit *L. iners* and promote growth of *L. crispatus* and other FGT lactobacilli

To investigate the effects of *cis-*9-uLCFAs on FGT *Lactobacillus* species, we cultured representative strains with varying concentrations of several *cis-*9-uLCFAs in modified *Lactobacillus* MRS broth (MRS+CQ broth)^43^. Strikingly, oleic acid (OA; 18:1 *cis-*9), linoleic acid (LOA; 18:2 *cis-*9,12), and palmitoleic acid (POA; 16:1 *cis-*9) each selectively and potently inhibited *L. iners* while exerting little or no inhibitory effect on *L. crispatus*, *L. gasseri*, *L. jensenii*, and *L. mulieris* (Figure 1A; corresponding *cis-*9-uLCFA chemical structures are shown in Figure S1A). In particular, OA had no inhibitory effect towards any of the non-*iners* species at concentrations up to 3.2 mM, despite robustly inhibiting *L. iners* at a half-maximal inhibitory concentration (IC50) of less than 400 µM. To assess whether these phenotypes showed species-level conservation, we tested a diverse collection of strains from each species, including isolates from geographically diverse donors with varying BV status^43^ (Table S1). All *L. iners* strains (n=14) were sensitive to OA with a median IC50 of 100 µM in MRS+CQ broth, while none of the non-*iners* FGT *Lactobacillus* strains (n=30) were inhibited (Figure 1B). Similar patterns were observed in NYCIII broth, a rich, non-selective media formulation commonly used to culture FGT bacteria^44^ (Figure S1B). These results demonstrate that OA is toxic to *L. iners* but exerts no inhibitory effect on non-*iners* FGT *Lactobacillus* species.

**Figure 1.**
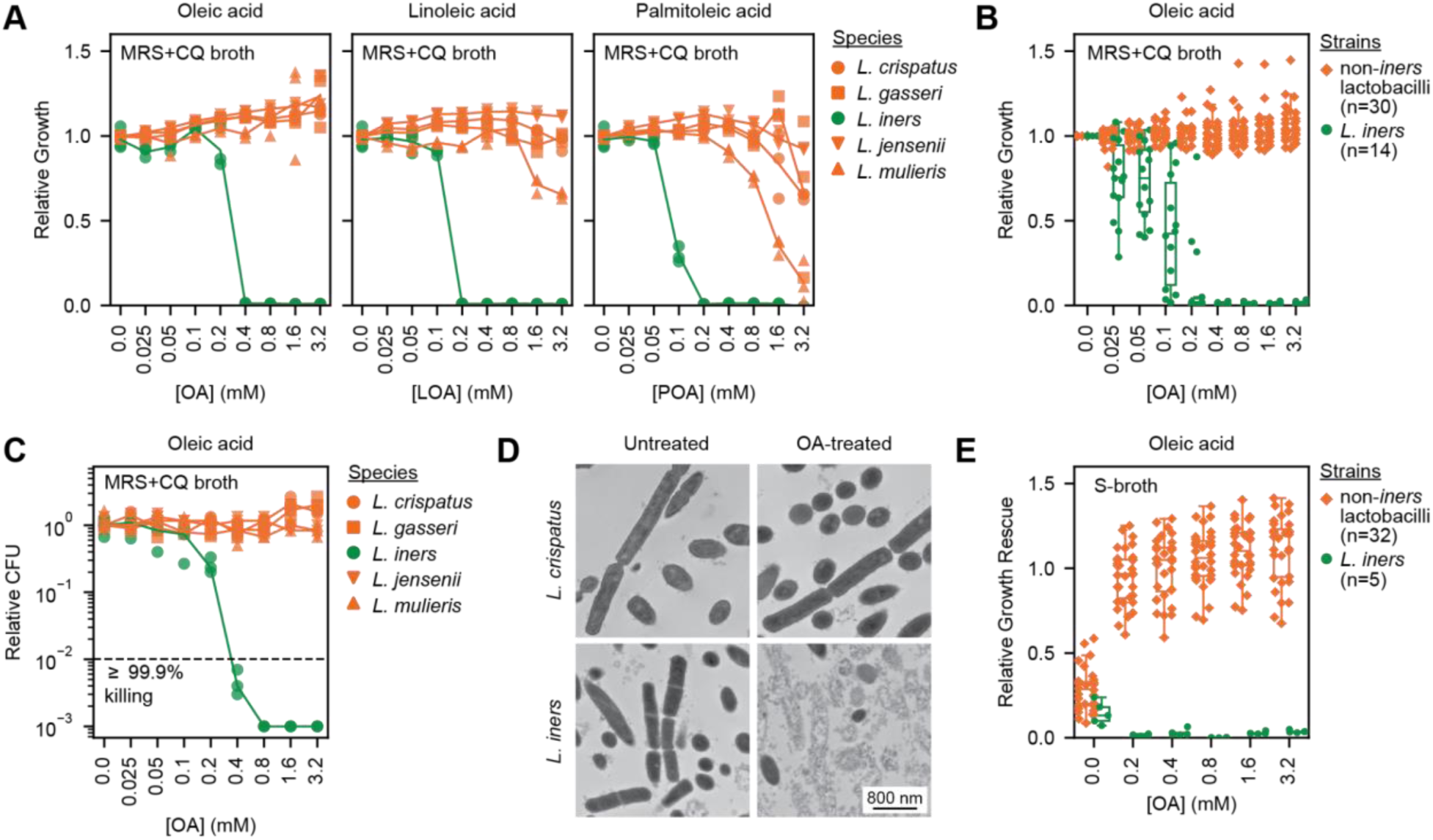
*cis*-9-uLCFAs selectively inhibit *L. iners* and promote growth of *L. crispatus* and other FGT lactobacilli. (A) Relative growth of representative *L. crispatus*, *L. gasseri*, *L. iners*, *L. jensenii*, and *L. mulieris* strains in modified *Lactobacillus* MRS broth (MRS+CQ broth) with varying concentrations of oleic acid (OA, left), linoleic acid (LOA, middle), or palmitoleic acid (POA, right). (B) Relative growth of diverse non-*iners* FGT *Lactobacillus* (n=30) and *L. iners* (n=14) strains in MRS+CQ broth supplemented with varying concentrations of OA. (C) Minimum bactericidal concentration (MBC) assay results for representative *L. crispatus*, *L. gasseri*, *L. iners*, *L. jensenii*, and *L. mulieris* strains in MRS+CQ broth. Colony forming units (CFU) were measured after 24 hours by standard serial dilution and colony counting and expressed relative to the number of CFU recovered from the no fatty acid supplementation control. (D) Transmission electron microscopy (TEM) images of *L. crispatus* (top) and *L. iners* (bottom) treated with 3.2 mM OA (right) or untreated (left) for 1 hour. (E) Relative growth rescue of diverse non-*iners* FGT *Lactobacillus* (n=32) and *L. iners* (n=5) strains in S-broth supplemented with varying concentrations of OA. Relative growth rescue was calculated as growth relative to the median OD600 measurement in the S-broth supplemented with 0.1% Tween-80. (A, B, and E) Growth was measured by optical density at 600 nm (OD600) after 72 hours of culture. (A and B) Relative growth was calculated relative to the median OD600 measurement in the no fatty acid supplementation control. (A and C) Plotted points represent 3 technical replicates per condition and are representative of ≥2 independent experiments per condition. (B and E) Plotted points represent the median relative growth or growth rescue for 3 technical replicates per condition and are representative of ≥2 independent experiments per condition.

To evaluate whether OA inhibited *L. iners* via a bactericidal mechanism, we performed a minimum bactericidal concentration (MBC) assay^45^ using representative *Lactobacillus* strains in MRS+CQ broth. The minimum OA concentration required to kill ≥99.9% of the *L. iners* inoculum within 24 hours (the MBC) was 400 µM (Figure 1C), which was equal to the corresponding MIC (minimum concentration achieving ≥99.9% growth inhibition, Figure 1A), indicating a bactericidal effect. In contrast, OA had no bactericidal activity against non-*iners Lactobacillus* species. Transmission electron microscopy (TEM) revealed that exposure to 3.2 mM of OA induced catastrophic cell wall and membrane disruption in *L. iners* within 1 hour of treatment, whereas *L. crispatus* cell integrity and morphology remained unaffected by OA (Figure 1D; full images, Supp. Data 1). These results indicate that *cis*-9-uLCFAs selectively kills *L. iners* via disruption of their cell wall and membrane.

We next investigated OA growth effects in conditions less optimized for *Lactobacillus* growth. S-broth is a rich media formulation in which FGT *Lactobacillus* species grow poorly if not supplemented with Tween-80^43^. Consistent with observations from other media types, supplementing S-broth with OA inhibited *L. iners* (n=5 strains; Figure 1E). However, OA supplementation in S-broth robustly promoted growth of non-*iners* FGT *Lactobacillus* species (n=32 strains). Thus, OA not only exerts potent, selective toxicity against *L. iners*, but simultaneously promotes growth of non-*iners* FGT lactobacilli.

### Non-*iners* FGT lactobacilli possess a conserved set of *cis*-9-uLCFA-response genes that *L. iners* lacks

We hypothesized that the similar growth responses of non-*iners* FGT *Lactobacillus* species to uLCFAs reflected a shared set of uLCFA-response genes. We therefore assessed transcriptomic responses of representative *L. crispatus*, *L. gasseri*, and *L. jensenii* strains to OA. Bacteria were grown to exponential phase, then treated with OA for 1 hour. We used bulk RNA-sequencing and DESeq2^46^ analysis to identify genes that were differentially expressed (DE) by each species compared to matched, untreated control cultures (Figure 2A). This analysis revealed just three unique gene functions were consistently DE (all upregulated) in all species in response to OA (Figures 2A-B). These included a predicted oleate hydratase enzyme (*ohyA*), a putative fatty acid efflux pump (*farE*), and its putative regulator (*tetR*). To further validate these findings, we investigated transcriptomic responses of the same strains to the *cis-*9-uLCFAs POA and LOA (Figure S2A). All three OA-induced gene functions (*ohyA*, *farE*, and *tetR*) were also upregulated by POA and by LOA treatment in all three species (Figures S2A-C; full dataset, Supp. Data 2), confirming the hypothesis that non-*iners* FGT lactobacilli shared a core set of *cis*-9-uLCFA-inducible response genes.

**Figure 2.**
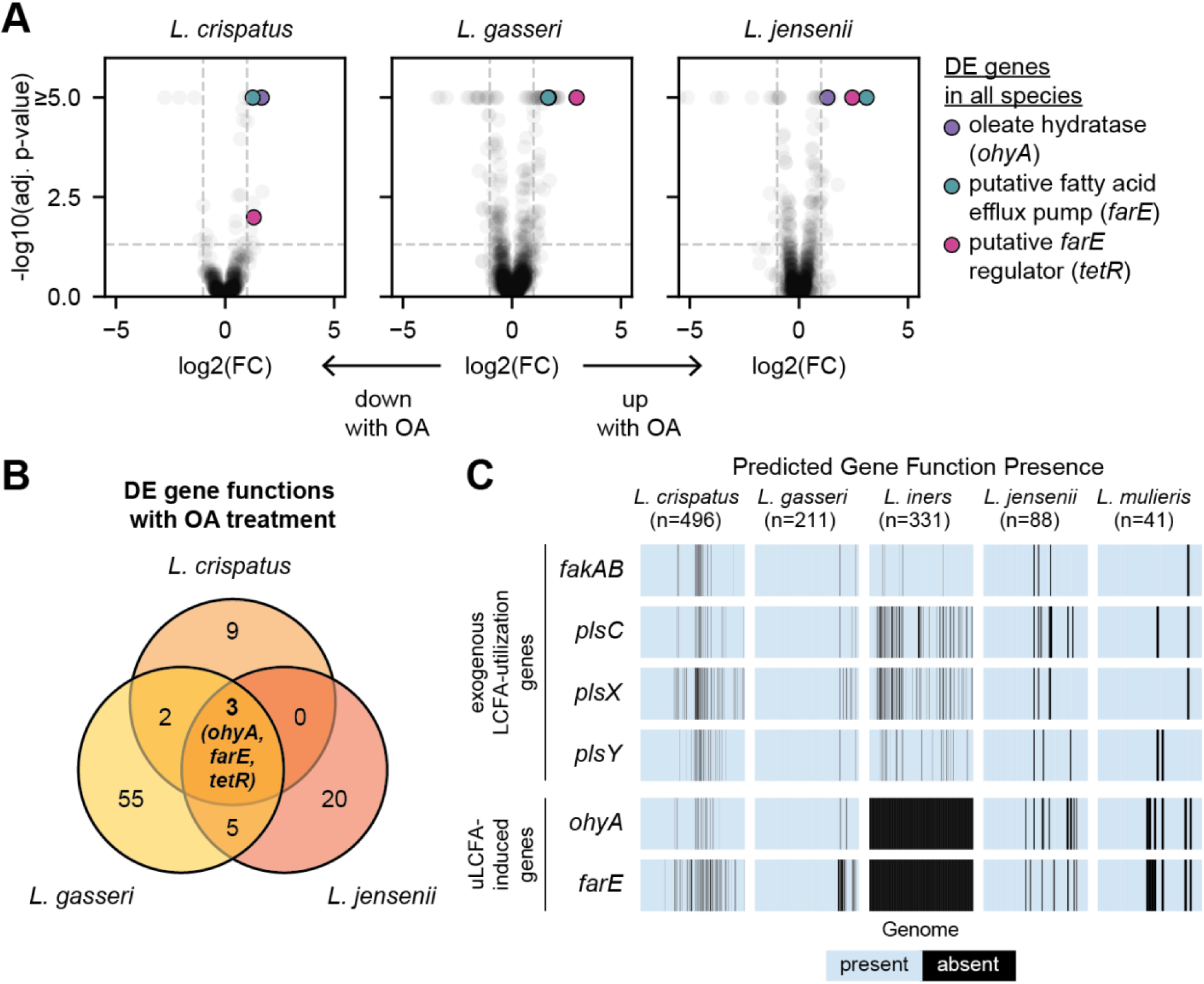
Non-*iners* FGT lactobacilli possess a conserved set of OA response genes that *L. iners* lacks. (A) Transcriptomic responses of cultured *L. crispatus* (left), *L. gasseri* (middle), and *L. jensenii* (right) grown to exponential phase in MRS+CQ broth, then exposed to OA (3.2 mM) for 1 hour. Bulk RNA-sequencing was performed and DESeq2^46^ analysis was used to identify differentially expressed (DE) genes relative to the matched, no OA supplementation control. The Benjamini-Hochberg procedure was used to control the false discovery rate (FDR) with ɑ=0.05. Plots depict the log2(fold change, FC) of mRNA expression with OA relative to control (x-axis) and - log10(adjusted p-value) for each gene (y-axis). Dotted lines represent cutoffs used to define significant differential expression (−1≥FC≥1 and adjusted p-value≤0.05). DE genes observed in all species included a predicted oleate hydratase (*ohyA*, purple; COG4716), putative fatty acid efflux pump (*farE*, teal; COG2409), and its putative regulator (*tetR*, pink; COG1309). (B) Venn diagram showing the number of DE genes per species with OA treatment and overlap of DE gene functions across species. Three gene functions (*ohyA*, *farE*, and *tetR*; all upregulated in response to OA) were shared among all three species. (C) Presence of gene functions predicted to encode functional oleate hydratase (*ohyA*) and putative fatty acid efflux pump (*farE*) activity in isolate genomes and metagenome-assembled genomes (MAGs) of the indicated FGT *Lactobacillus* species (n=1,167 total genomes and MAGs)^43^. Presence of gene functions involved in exogenous fatty acid acquisition and utilization (*fakAB*, *plsC*, *plsX*, and *plsY*) are shown for comparison. (B and C) Gene function was predicted using eggNOG 5.0^85^ employing eggNOG-mapper v2.1.9^86^.

Given its strikingly different phenotypic response to OA exposure, we hypothesized that *L. iners* would lack the OA-induced gene functions found in non-*iners* lactobacilli. We therefore investigated gene function presence in a previously reported FGT *Lactobacillus* genome catalog comprising 1,167 isolate genomes and metagenome-assembled genomes (MAGs) from geographically and clinically diverse sources, supplemented with six additional completed *L. iners* isolate genomes (Table S2)^43,47^. We excluded *tetR*, the putative *farE* regulator, from this analysis due to its nonspecific functional annotation. As hypothesized, *ohyA* and *farE* gene functions were completely absent from *L. iners* genomes but highly conserved among non-*iners* FGT *Lactobacillus* genomes (Figure 2C). In contrast, gene functions involved in acquiring (*fakA* and *fakB*) and utilizing (*plsC*, *plsX*, and *plsY*) exogenous fatty acids for phospholipid synthesis were conserved among all FGT *Lactobacillus* species (Figure 2C)^48^, showing that *L. iners* retains other key fatty acid-related pathways. Collectively, these transcriptomic and genomic analyses suggest roles for *ohyA* and *farE* in mediating species-specific uLCFA growth phenotypes in FGT lactobacilli.

### Comparative genomics reveals conserved phylogeny of FGT *Lactobacillus ohyA* and *farE*

To our knowledge, *ohyA* and *farE* have not previously been studied in FGT *Lactobacillus* species. We therefore investigated their intraspecies diversity and their homology to better-characterized orthologs from other human-adapted bacteria. Most *L. crispatus* and *L. gasseri* genomes contained two distinct predicted *ohyA* orthologs (only one of which was OA-induced), whereas *L. jensenii* and *L. mulieris* genomes each contained a single ortholog (Figure S3A). In contrast, each species contained a single *farE* ortholog (Figure S3B). To compare FGT *Lactobacillus ohyA* orthologs to other species, we performed a phylogenetic reconstruction comprising 21 distinct OhyA protein sequences from 16 different species, including all 7 distinct FGT *Lactobacillus* OhyA orthologs (Figure 3A; percent identity matrix shown in Figure S3C). Interestingly, patterns of *Lactobacillus* OhyA phylogeny differed from underlying species phylogeny. The OA-induced *L. crispatus* and *L. gasseri* orthologs (LCRIS_00661 and LGAS_1351, respectively) clustered with previously characterized orthologs from *S. aureus*^49,50^, *Bifidobacterium breve*^51^, *S. pyogenes*^52^, and *Lactobacillus acidophilus*^53,54^, while the non-OA-induced *L. crispatus* and *L. gasseri* orthologs (LCRIS_00558 and LGAS_0484, respectively) clustered separately. The *L. jensenii* and *L. mulieris* OhyA orthologs constituted a more distantly related phylogenetic clade, clustering with orthologs from *Streptococcus salivarius* and *Enterococcus faecium*. In contrast, phylogenetic reconstruction of FarE orthologs from FGT and non-FGT *Lactobacillus* species closely matched underlying species phylogeny (Figures S4A and S4B). FGT *Lactobacillus* FarE orthologs shared more distant homology (18-20% protein sequence identity) to the previously characterized *S. aureus* FarE, which also neighbors a TetR family regulator (Figures S4C and S4D). Together, these phylogenetic analyses reveal that non-*iners* FGT *Lactobacillus* species harbor phylogenetically diverse *ohyA* orthologs, suggesting prior horizontal gene transfer and/or species-specific gene loss, whereas each species harbors a single *farE* ortholog that was likely acquired vertically.

**Figure 3.**
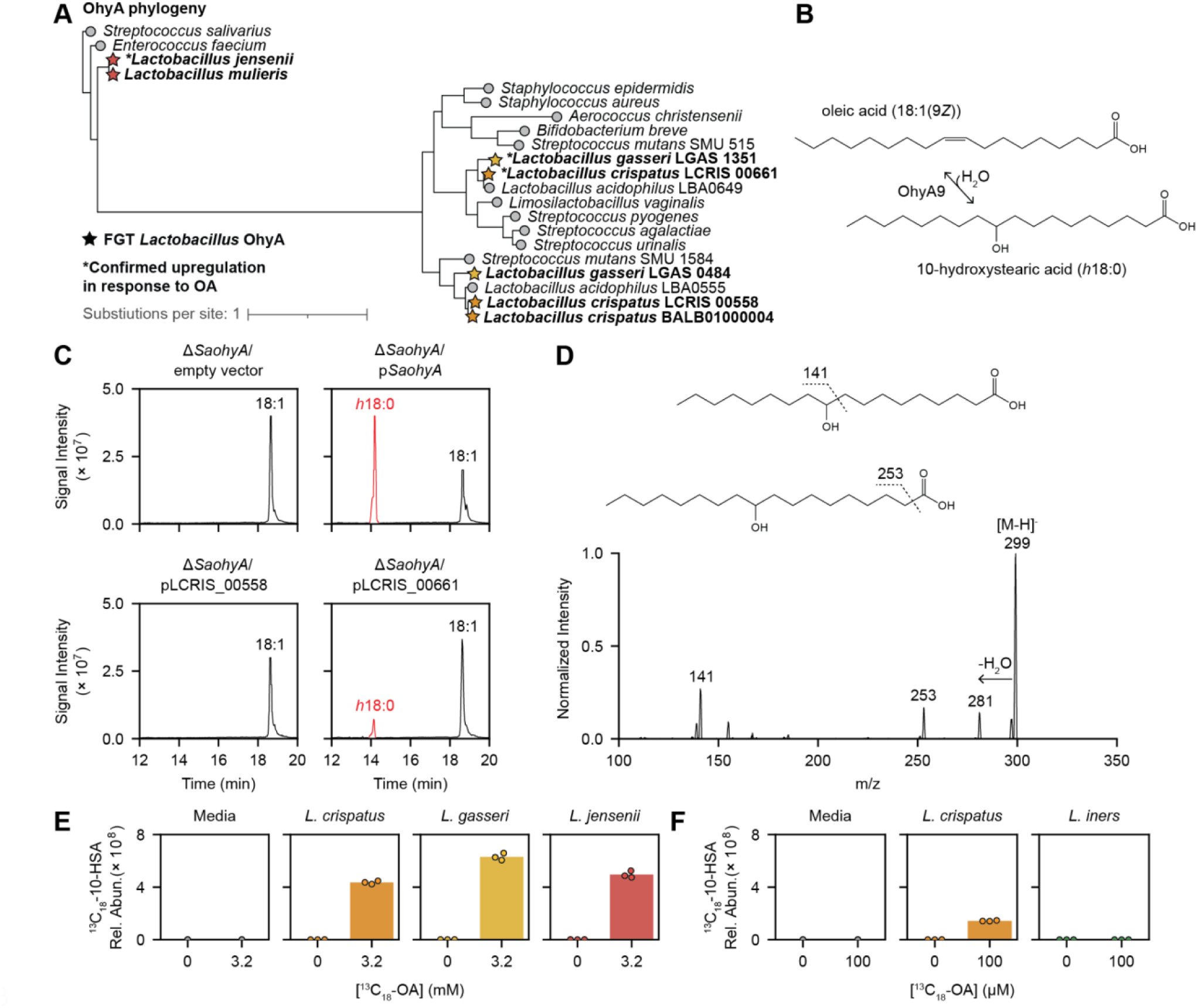
FGT *Lactobacillus* OhyA enzymes are functional and physiologically active. (A) OhyA protein phylogenetic tree for representative orthologs from the indicated species. Tree was constructed from MUSCLE v5.1^87^-aligned protein sequences using RAxML-NG^88^ (see methods). Starred leaf tips correspond to FGT *Lactobacillus* orthologs shown in bold. OhyA orthologs upregulated in response to OA (Figure 2A) are marked with asterisks. (B) OhyA9 enzymatic activity reaction diagram with OA substrate. (C) Extracted ion chromatograms from supernatants of *ohyA*-gene deleted strain of *S. aureus*^49^ complemented with an empty vector (*ΔSaohyA*/empty vector), *SaohyA*-expressing plasmid (*ΔSaohyA*/p*SaohyA*), LCRIS_00558-expressing plasmid (*ΔSaohyA*/pLCRIS_00558), and LCRIS_00661-expressing plasmid (*ΔSaohyA*/pLCRIS_00661). Strains were cultured with OA for 1 hour. Annotated peaks include OA (18:1) and 10-HSA (*h*18:0). (D) MS2 spectra with major fragmentation labels for the *h*18:0 peak shown in the lower right panel of Figure 3C for the supernatant of *ΔSaohyA*/pLCRIS_00558 cultured with OA. (E) Detection of ^13^C-labeled 10-hydroxystearic acid (^13^C_18_-10-HSA; *h*18:0) in supernatants of *L. crispatus*, *L. gasseri*, and *L. jensenii* cultured for 72 hours in NYCIII broth with and without universally ^13^C-labeled OA (^13^C_18_-OA; 3.2 mM, which is a lethal concentration for *L. iners*). (F) Detection of ^13^C_18_-10-HSA in supernatants of *L. crispatus* and *L. iners* cultured for 72 hours in NYCIII broth with and without ^13^C_18_-OA (100 µM, which is a sublethal concentration for *L. iners*). (E and F) The no-OA control data shown for media and *L. crispatus* supernatant in E and F are derived from the same samples. Plotted points represent 3 technical replicates per condition.

We next investigated the presence and co-occurrence of *farE* and *ohyA* within the Lactobacillaceae family, including species with diverse lifestyles (e.g., vertebrate-associated, insect-associated, free living, or nomadic)^55^. Many species encoded both *farE* and *ohyA*, although the genes did not co-occur in all cases (Figure S5). Notably, *L. iners* was unique among species in the genus *Lactobacillus* to lack *farE* and was the only vertebrate-associated *Lactobacillus* species to lack *ohyA*. The widespread distribution of *farE* and *ohyA* within the Lactobacillaceae family suggests an important conserved role in *cis*-9-uLCFA responses.

### FGT *Lactobacillus* OhyA enzymes are functional and physiologically active

Enzymatic activities of the predicted OhyA orthologs from non-*iners* FGT lactobacilli have not previously been determined, so we assessed their function by heterologous expression in a well-characterized *ohyA*-knockout strain of *S. aureus* (Δ*SaohyA*)^49^. We complemented the Δ*SaohyA* strain with each of the two most common *L. crispatus ohyA* orthologs (LCRIS_00661 and LCRIS_00558), along with the *S. aureus ohyA*^49,50,56^ (*SaohyA*) as a positive control, and empty vector as a negative control. Supernatants from the complemented Δ*SaohyA* strains cultured with OA or LOA were harvested for identification of OhyA-produced metabolites by targeted lipidomics. The strain expressing the OA-inducible OhyA ortholog from *L. crispatus* (LCRIS_00661) produced the same hydroxy fatty acid (*h*FA) metabolites as the *SaohyA*-expressing strain when cultured with OA or with LOA, confirming they shared the biochemical activity of hydrating the *cis*-9 double bond to produce 10-hydroxystearic acid (10-HSA or *h*18:0) from OA and 10-hydroxy-12-octadecenoic acid (*h*18:1) from LOA (OA reaction, Figures 3B-D; 10-HSA standard MS2 spectra, Figure S6A; LOA reaction, Figures S6B-D). Thus, we named this *L. crispatus* ortholog OhyA9. In contrast, when complemented with the *L. crispatus* OhyA ortholog that was not induced by OA (LCRIS_00558), the *ΔSaohyA* strain did not produce *h*FAs when cultured in presence of OA, showing it lacked ability to hydrate OA’s *cis*-9 double bond (Figure 3C). However, it did hydrate the *cis*-12 double bond of LOA to produce 13-hydroxy-9-octadecenoic acid, so we called this ortholog OhyA12 (Figures S6B and S6E-F). These results show that OhyA orthologs in *L. crispatus* have distinct enzymatic activities, with only the OA-induced ortholog (OhyA9) exhibiting the ability to hydrate *cis*-9 double bonds of OA and related *cis*-9-uLCFAs.

We next sought to assess physiologic OhyA9 enzymatic activity in *L. crispatus* and other non-*iners* FGT *Lactobacillus* species and to confirm the genomic prediction that *L. iners* lacks OhyA activity. We cultured representative strains of *L. crispatus*, *L. iners*, *L. gasseri*, and *L. jensenii* in NYCIII broth supplemented with universally ^13^C-labeled OA (^13^C_18_-OA). Concentrations of the labeled OhyA9 product 10-HSA (^13^C_18_-10-HSA) were measured in culture supernatants and cell pellets. We observed robust production of ^13^C_18_-10-HSA by all three non-*iners* species in both media and cell pellets, confirming that each species has a physiologically active OhyA9 (supernatants, Figure 3E; pellets, Figure S6G). By contrast, *L. iners* did not produce ^13^C_18_-10-HSA when cultured in a sub-lethal concentration of ^13^C_18_-OA (supernatants, Figure 3F; pellets, Figure S6H). Collectively these results confirmed transcriptomic and genomic predictions that non-*iners* FGT lactobacilli encode and express physiologically active OhyA9 enzymes, while *L. iners* does not.

### Women with non-*iners* lactobacilli have uniquely elevated vaginal concentrations of OhyA products

We hypothesized that the OhyA activity observed *in vitro* would also be active in the vaginal microbiota of women with bacterial communities dominated by non-*iners* FGT lactobacilli. We tested this hypothesis by measuring concentrations of OhyA enzymatic products in cervicovaginal lavage (CVL) fluid samples from participants in the FRESH (Females Rising through Education, Support and Health) study, which enrolls South African women aged 18–23 years who provide cervicovaginal samples at three-month intervals^57^. We examined 180 distinct samples from 106 unique FRESH study participants (Table S3). We profiled microbiota composition via bacterial 16S rRNA gene sequencing of matched vaginal swabs and classified bacterial communities into distinct cervicotypes (CTs) using previously validated criteria^10,19,43^ (full microbiome dataset, Supp. Data 4). Briefly, samples were classified as cervicotype 1 (CT1, n=34 samples), defined based on dominance by non-*iners Lactobacillus* species consisting primarily of *L. crispatus*; CT2 (n=57), defined by dominance of *L. iners*; CT3 (n=60), defined by *Gardnerella* predominance; or CT4 (n=29), which consists of non-*Lactobacillus*-dominant diverse anaerobes, typically with high *Prevotella* abundance^10^ (Figure 4A). We measured concentrations of hydroxy fatty acids (*h*FAs) including *h*18:0 and hydroxy 18:1 (*h*18:1) in paired cervicovaginal lavage (CVL) supernatant samples using a validated targeted lipidomics approach involving picolylamine-based derivatization^56,58^ (representative human sample chromatograms shown in Figures S7A-B). *h*18:0 and *h*18:1 concentrations were significantly and uniquely elevated in CT1 compared to all other CTs, with *h*18:0 exhibiting 4.8-fold higher median concentration in CT1 samples compared to other samples and *h*18:1 exhibiting a 3.6-fold higher median concentration (Figure 4B). Remarkably, *h*18:0 and *h*18:1 concentrations segregated CT1 samples from other samples with near-perfect sensitivity and specificity, as shown by respective area under the curve (AUC) values of 0.99 and 1.00 for *h*18:0 and *h*18:1 in receiver operating characteristic analysis. These results confirm robust *in vivo* physiologic activity of non-*iners Lactobacillus* OhyA enzymes in the human FGT microbiota.

**Figure 4.**
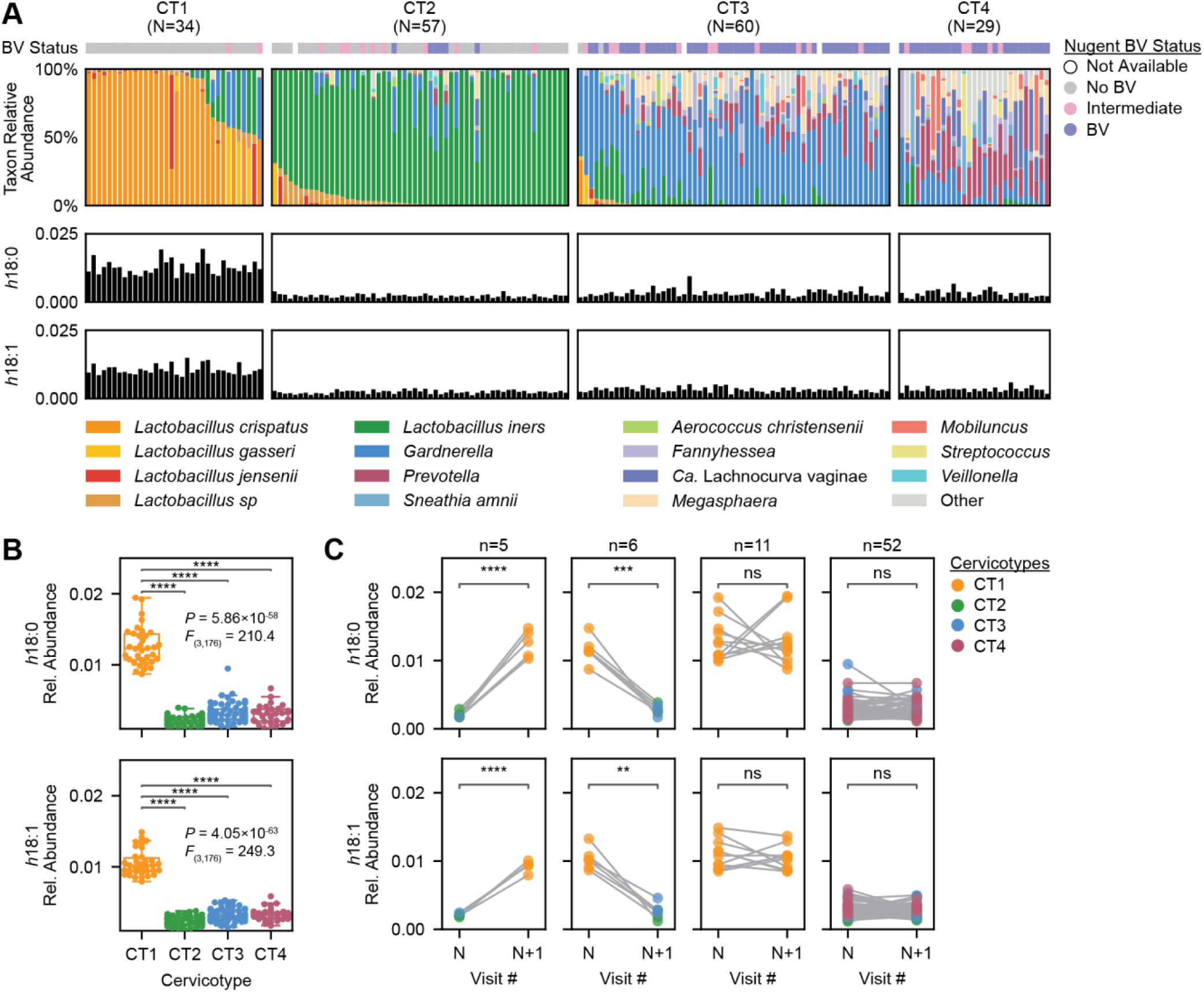
Women with non-*iners* lactobacilli have uniquely elevated vaginal concentrations of OhyA products. (A) FGT bacterial microbiota composition of 180 distinct FGT swab samples from 106 total South African women, determined by bacterial 16S rRNA gene sequencing (top, stacked barplot). Samples were classified into cervicotype (CT) based on microbiota composition as previously described^10,43^. Middle and bottom bar plots show relative concentrations of OhyA products 10-HSA (*h*18:0) and hydroxy 18:1 (*h*18:1) respectively, measured in paired cervicovaginal lavage (CVL) samples by targeted lipidomics. The top colorbar shows Nugent score-based BV status^89^. (B) Relative *h*18:0 (top) and *h*18:1 (bottom) concentrations within each CT for the 180 CVL samples shown in (A). Significance was determined by one-way ANOVA with post-hoc Tukey’s test; selected pairwise differences are shown (****p < 0.0001; full statistical results in Table S4). (C) The change in relative *h*18:0 (top) and *h*18:1 (bottom) concentrations within 74 paired consecutive samples from participants whose microbiota transitioned to CT1 (n=5), away from CT1 (n=6), remained CT1 (n=11), or remained non-CT1 (n=52). Significance was determined by paired t-test on log-transformed values (**p < 0.01; ***p < 0.001; ****p < 0.0001; ns: p ≥ 0.05).

To further establish the relationship between microbiota composition and OhyA activity, we investigated how individual women’s cervicovaginal *h*18:0 and *h*18:1 levels changed over time in serially collected samples. Since some participants provided samples at multiple study visits, the dataset included 74 pairs of consecutively collected samples, including 5 sample pairs in which a participant transitioned from a non-CT1 community to a CT1 community; 6 pairs in which a participant transitioned from a CT1 to a non-CT1 community; 11 pairs in which a participant remained CT1; and 52 pairs in which a participant remained non-CT1. Relative concentrations of *h*18:0 and *h*18:1 significantly increased in CVL samples from participants who transitioned from non-CT1 to CT1 (p=0.00006 with median fold-increase 7.11 for *h*18:0; p=0.00003 with median fold-increase 4.50 for *h*18:1), and significantly decreased in samples from participants who transitioned away from CT1 (p=0.000256 with median fold-decrease 4.12 for *h*18:0; p=0.00118 with median fold-decrease 3.88 for *h*18:1; Figure 4C). By contrast, *h*18:0 and *h*18:1 levels did not significantly change in participants who remained in CT1 or remained in non-CT1 states (median fold changes for these transitions ranged between 0.96 and 1.02 with p>0.05 for all types of sample pairs and metabolites). Thus, changes in levels of OhyA enzymatic products within cervicovaginal fluid of individual women closely tracked with bacterial community changes over time, further confirming the close relationship between OhyA activity and microbiota composition.

### *farE* is required for OA resistance and growth enhancement

We used a genetic approach to interrogate the relationship between OA growth phenotypes and the conserved, OA-induced genes, *ohyA9* and *farE*. Tools to genetically modify *L. crispatus* and *L. iners* were not available, so we conducted these studies in *L. gasseri* because of the close phenotypic, genomic, transcriptional, and phylogenetic resemblance of its *cis-*9-uLCFA responses to those of *L. crispatus*. Using adaptations of previously reported methods, we made genetic knockouts of *ohyA9* (*ΔohyA9*) and *farE* (*ΔfarE*) in the ATCC 33323 strain of *L. gasseri* via in-frame gene deletions generated by homologous recombination (Figure S8A; WGS verification, Supp. Data 3; see methods). We found that *farE*, but not *ohyA9*, was required for the OA resistance and growth enhancement phenotypes (Figures 5A-B). *ΔfarE* was fully inhibited by OA in MRS+CQ broth (Figures 5A) and failed to exhibit growth with OA supplementation in delipidated MRS+CQ broth (Figure 5B). Complementing *ΔfarE* with plasmid-overexpressed *farE* (*ΔfarE*/p*farE*) fully rescued both the inhibition and growth phenotypes (Figures 5A-B). However, OA-mediated growth inhibition in *ΔfarE* was not as potently bactericidal as in *L. iners*, indicating that additional factors may also contribute to *L. iners* susceptibility (Figure 5C). Together, these results show FarE is required for OA resistance and growth enhancement in non-*iners* FGT lactobacilli.

**Figure 5.**
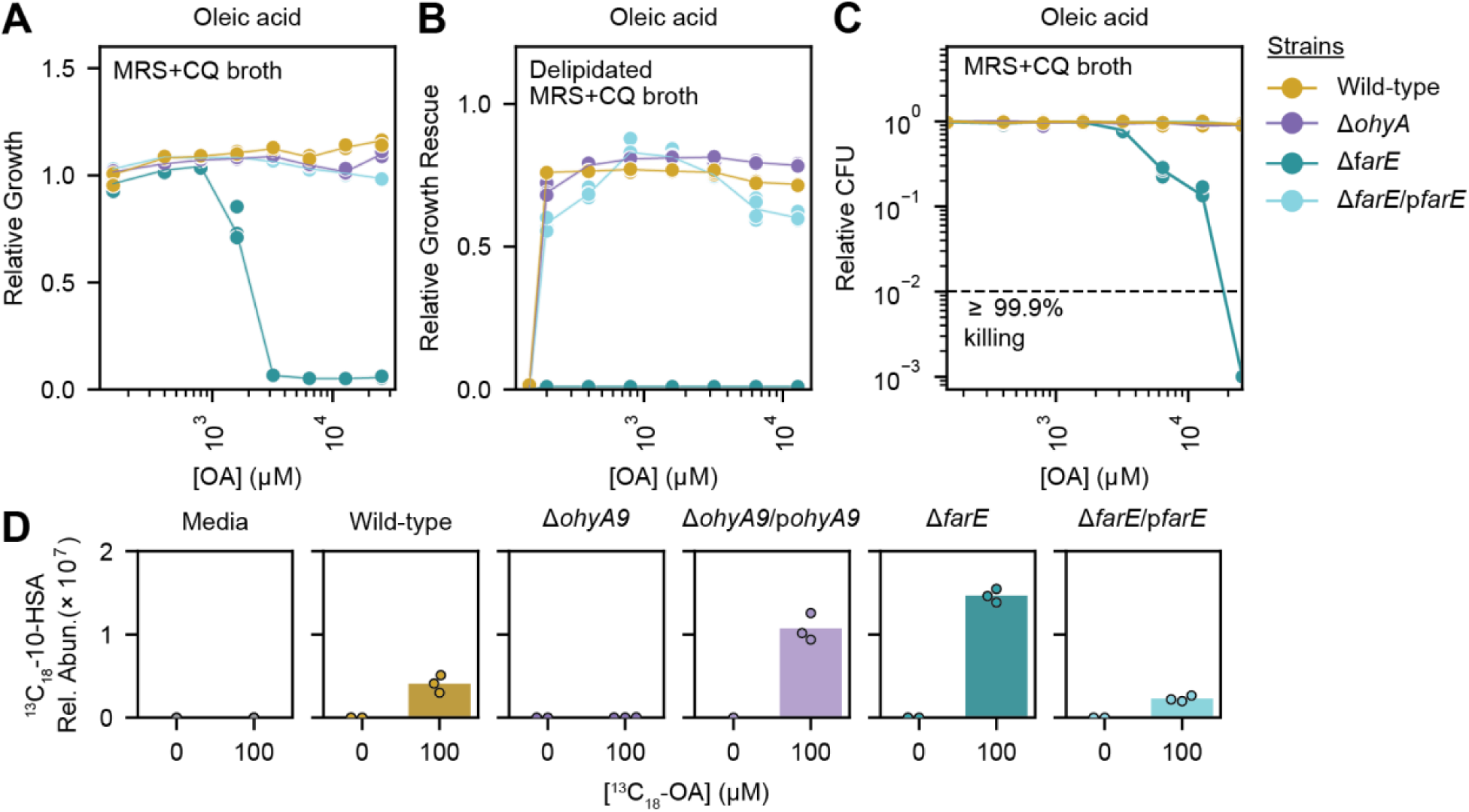
*farE* is required for OA resistance and growth enhancement. (A) Relative growth of *L. gasseri* ATCC 33323 wild-type (WT) strain, and derivative genetic mutant strains created by double crossover homologous recombination, including knockout strains of *ohyA9* (*ΔohyA9*) and *farE* (*ΔfarE*), and *ΔfarE* complemented with plasmid-overexpressed autologous *farE* (*ΔfarE*/p*farE*) in MRS+CQ broth supplemented with varying concentrations of OA. (B) Relative growth rescue of *L. gasseri* ATCC 33323 WT and derivative genetic mutant strains in delipidated MRS+CQ broth supplemented with varying concentrations of OA. (C) MBC assay results for *L. gasseri* ATCC 33323 WT and derivative genetic mutant strains in MRS+CQ broth. MBC assay performed following the same methods described in Figure 1C. (D) Detection of ^13^C_18_-10-HSA in blank media and culture supernatants from *L. gasseri* WT, *ΔohyA9*, *ΔohyA9* complemented with plasmid-overexpressed autologous *ohyA9* (*ΔohyA9*/p*ohyA9*), *ΔfarE*, and *ΔfarE*/p*farE* cultured for 24 hours in NYCIII broth with and without ^13^C_18_-OA (100 µM). Plotted points represent 2 or 3 technical replicates per condition. (A and B) Growth was measured by OD600 after 24 hours of culture. Relative growth or growth rescue was calculated relative to the median OD600 measurement in the non-delipidated media, no OA supplementation control. (A, B, and C) Plotted points represent 3 technical replicates per condition and are representative of ≥2 independent experiments per condition.

To genetically confirm the role of *ohyA9* in 10-HSA production, we grew *L. gasseri* wild-type (WT), *ΔfarE*, *ΔfarE*/p*farE*, *ΔohyA9*, and *ΔohyA9* complemented with plasmid-overexpressed *ohyA9* (*ΔohyA9*/p*ohyA9*) in media supplemented with a sublethal concentration of ^13^C_18_-OA and measured ^13^C_18_-10-HSA production in media supernatants and cell pellets. As hypothesized, knocking out *ohyA9* fully ablated ^13^C_18_-10-HSA production, which was rescued by genetic complementation (Figures 5D and S8B). Interestingly, ^13^C_18_-10-HSA production and secretion into supernatant was increased in *ΔfarE* but decreased in *ΔfarE*/p*farE* relative to wild-type, indicating that presence of FarE modifies 10-HSA production and export, but is not required. Based on these results, we propose that OA-induced genes in non-*iners* FGT *Lactobacillus* species are required for two major mechanisms of response to exogenous *cis*-9-uLCFAs: FarE-mediated transport to prevent toxic intracellular accumulation and OhyA9-mediated bioconversion to produce their *h*FA counterparts (model schematic, Figure S8C).

### FGT *Lactobacillus* species are fatty acid auxotrophs

The observation that OA supplementation enhanced non-*iners Lactobacillus* growth in S-broth, a media not optimized for lactobacilli, prompted us to hypothesize that FGT *Lactobacillus* species are fatty acid auxotrophs. To confirm this hypothesis, we investigated FGT *Lactobacillus* genomes for presence of fatty acid synthesis II (FASII) pathway gene functions^59^. FGT *Lactobacillus* species were largely predicted to lack an intact FASII pathway, including genes predicted to perform the functions of AccABCD, FabH, FabB/F, and FabA/Z (Figures 6A and S9A). Additionally, *L. iners* lacked genes predicted to perform the functions of FabG and FabD. Among all FGT *Lactobacillus* genomes examined, only a minority of *L. crispatus* genomes were predicted to contain an intact FASII pathway. We therefore investigated growth of diverse FGT *Lactobacillus* species and strains in delipidated MRS+CQ broth supplemented with OA or with acetate, a precursor to the FASII initiating molecule, acetyl-Coenzyme A. All strains, including predicted FASII-containing *L. crispatus* strains, failed to grow in delipidated media and in acetate-supplemented delipidated media, suggesting all were fatty acid auxotrophs (Figure 6B). In contrast, OA supplementation in delipidated media largely restored growth of non-*iners* FGT *Lactobacillus* species (n=45 strains) at levels equivalent to growth in untreated media, while *L. iners* failed to grow with OA supplementation (n=13 strains; Figure 6B). Acetate supplementation was not inhibitory towards non-*iners* FGT *Lactobacillus* species in non-delipidated media, demonstrating that acetate’s failure to rescue growth was due to these species’s inability to use it as a FASII precursor rather than direct toxicity (n=45 strains; Figure S9B). Taken together, these results confirm that all FGT *Lactobacillus* species are fatty acid auxotrophs but only *farE*-harboring species can withstand and utilize high concentrations of OA to enhance growth.

**Figure 6.**
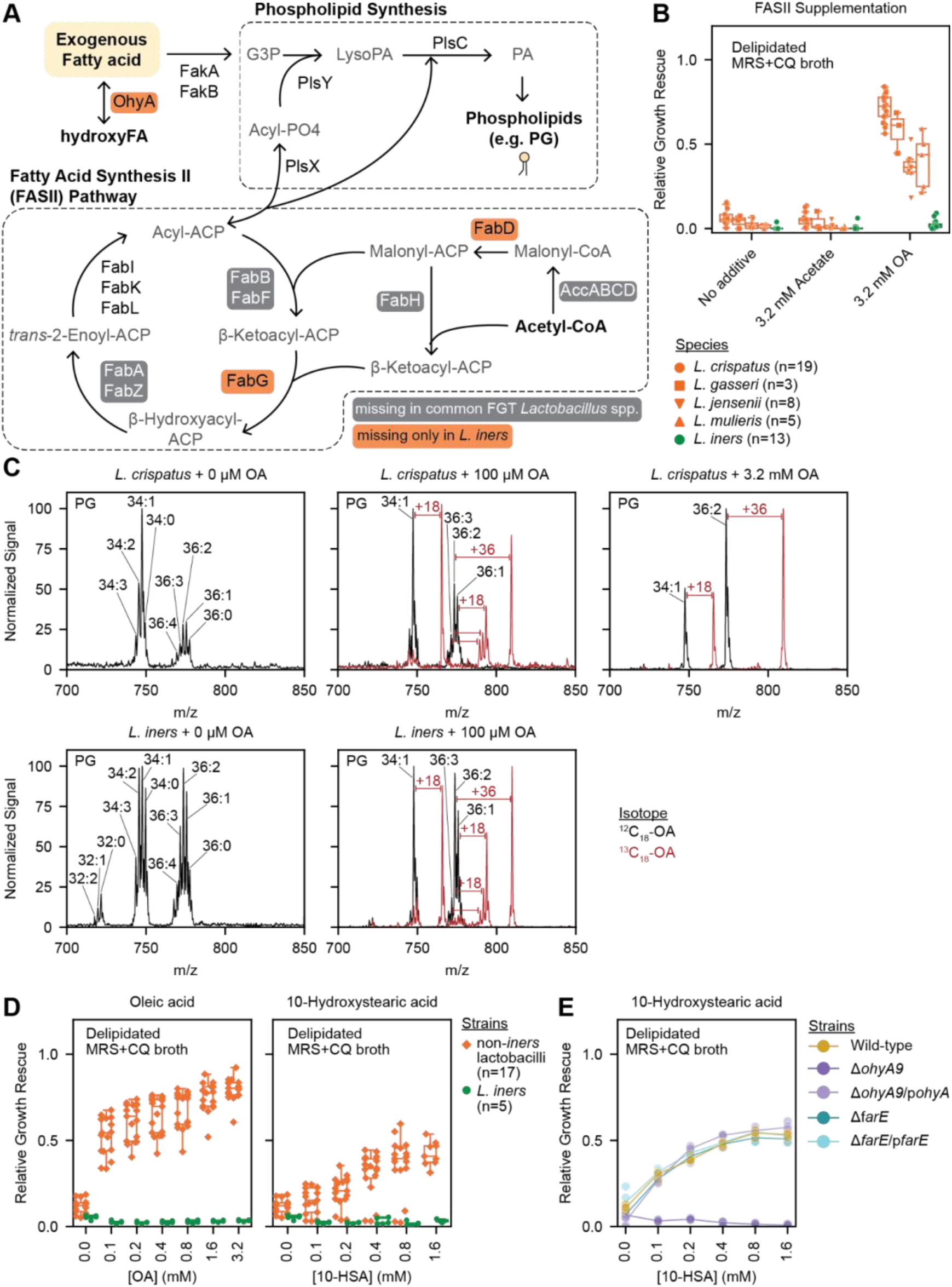
FGT lactobacilli are fatty acid auxotrophs and OhyA9 activity permits non-*iners* lactobacilli to utilize 10-HSA. (A) Fatty acid synthesis II (FASII) and phospholipid synthesis pathways annotated with the predicted presence of gene functions in *L. iners* and in non-*iners* FGT *Lactobacillus* genomes. Genes marked as missing for a species if the gene function that was predicted to be absent in >50% of genomes for each species. (B) Relative growth rescue of diverse *L. crispatus* (n=19), *L. gasseri* (n=3), *L. iners* (n=13), *L. jensenii* (n=8), and *L. mulieris* (n=5) strains in delipidated MRS+CQ broth supplemented with 3.2 mM acetate or 3.2 mM OA after 72 hours of culture. (C) Detection of phosphatidylglycerol lipids in cell pellets from *L. crispatus* (top) and *L. iners* (bottom) cultured for 72 hours in NYCIII broth with no OA supplementation (left), 100 µM OA (middle), or 3.2 mM OA (right). Unlabeled OA (^12^C_18_-OA) or universally ^13^C-labeled OA (^13^C_18_-OA) was used for supplementation. Plots represent MS1 spectra of one representative sample per condition with major unlabeled (black) and ^13^C-labeled (red) lipid species annotated. (D) Relative growth rescue of diverse non-*iners* FGT *Lactobacillus* (n=17) and *L. iners* (n=5) strains in delipidated MRS+CQ broth supplemented with varying concentrations of OA (left) or 10-HSA (right) after 72 hours of culture. (E) Relative growth rescue of *L. gasseri* ATCC 33323 WT and derivative genetic mutant strains in delipidated MRS+CQ broth supplemented with varying concentrations of 10-HSA after 24 hours of culture. (B, D, and E) Relative growth rescue was calculated as growth relative to the median OD600 measurement in non-delipidated MRS+CQ broth with no OA supplementation. (B and D) Plotted points represent the median relative growth for 3 technical replicates per condition. (E) Plotted points represent 3 technical replicates per condition and are representative of ≥2 independent experiments per condition.

Based on these findings, we hypothesized that FGT lactobacilli may utilize exogenous OA as precursors for phospholipid synthesis. To characterize exogenous OA utilization, we used ^13^C_18_-OA to isotopically trace the ^13^C label in potential downstream metabolites and biosynthetic pathways. As hypothesized, we observed the ^13^C label to be incorporated into structural lipid metabolites, such as phosphatidylglycerol (PG) and diglyceride lipids, in all FGT *Lactobacillus* species (Figures 6C and S9C). These were primarily observed in PG34:1, PG36:1, and PG36:2. Notably, membranes of *L. crispatus*, *L. gasseri*, and *L. jensenii* grown in high OA concentrations consisted largely of the OA-containing PGs, PG34:1 and PG36:2, indicating immense versatility of these organisms to incorporate exogenous OA into their membranes (Figures 6C and S9D). In contrast to their ability to use exogenous fatty acids for phospholipid synthesis, genomic analysis of FGT *Lactobacillus* species revealed no predicted beta-oxidation pathway gene functions. We experimentally confirmed this prediction by showing that no ^13^C label was incorporated into TCA cycle metabolites, indicating that FGT lactobacilli do not exploit OA for central carbon metabolism (Figure S9C). Thus, all FGT *Lactobacillus* species rely on environmental fatty acids to build their membranes while *farE*-harboring non-*iners* FGT lactobacilli possess a greater ability to resist and utilize high levels of exogenous OA.

### Non-*iners* FGT lactobacilli use OhyA to sequester OA in a derivative form only they can exploit

We hypothesized that since OhyA activity can be bidirectional, OhyA-mediated bioconversion of uLCFAs to their *h*FA counterparts might allow non-*iners* FGT *Lactobacillus* species to sequester OA in a derivative form that they could exploit but that competing *ohyA*-deficient species could not. To test this hypothesis, we assessed whether supplementation with 10-HSA (*h*18:0), the OhyA9-produced metabolite that is highly elevated in bacterial communities dominated by non-*iners* FGT lactobacilli, could enhance growth of these species in a similar manner to unmodified OA. As hypothesized, non-*iners* FGT *Lactobacillus* species (n=17 strains) exhibited growth enhancement when cultured in delipidated MRS+CQ broth supplemented with either OA or 10-HSA, while *L. iners* (n=5 strains) failed to grow in either condition (Figure 6D). 10-HSA was also inhibitory towards *L. iners*, albeit with somewhat lower potency than unmodified OA (∼4-fold higher MIC; Figure S9F). These results confirmed that only non-*iners* FGT *Lactobacillus* species derived a growth benefit from 10-HSA.

To determine whether *ohyA9* was required for this 10-HSA-driven growth benefit, we performed 10-HSA growth experiments in delipidated media using the *L. gasseri* genetic mutant strains (WT, *ΔfarE*, *ΔfarE*/p*farE*, *ΔohyA9*, and *ΔohyA9*/p*ohyA9*). The *ΔohyA9* strain failed to grow when supplemented with 10-HSA, but this growth defect was rescued in the *ΔohyA9*/p*ohyA9* complemented strain (Figure 6E), confirming that *ohyA9* was necessary for 10-HSA-dependent growth enhancement. Importantly, 10-HSA did not inhibit *ΔohyA9* in non-delipidated MRS+CQ broth, indicating that the lack of 10-HSA growth enhancement was due to inability to bioconvert 10-HSA to OA rather than direct toxicity (Figure S9E). These results show that non-*iners* FGT lactobacilli can employ OhyA9 to bioconvert and sequester OA in a derivative form that they are able to exploit in an OhyA9-dependent manner for growth.

### OA inhibits growth of key BV-associated bacteria, including MTZ-resistant strains

To evaluate the potential of OA to improve BV treatment, we characterized effects of OA alone and in combination with MTZ on growth of various non-*Lactobacillus* FGT bacteria, including canonically BV-associated species (Figures 7A and S10A-B). Remarkably, mono-cultures of the BV-associated species *Gardnerella vaginalis*, *Gardnerella piotii*, *Fannyhessea* (formerly *Atopobium*) *vaginae*, *Sneathia amnii*, and *Prevotella timonensis* were all robustly inhibited by OA, while other *Prevotella* species including *P. bivia, P. amnii,* and *P. disiens* were largely unaffected in NYCIII broth (Figure 7A). These assays were repeated in S-broth with similar results except for additional OA-driven inhibition of *P. amnii* (Figure S10B). Interestingly, even MTZ-resistant *G. piotii* and *F. vaginae* strains were susceptible to OA inhibition. To further characterize these OA-sensitive non-*Lactobacillus* species, we performed dose-response assays in NYCIII broth and observed *Gardnerella* MICs to be approximately 2-fold greater than the median MIC of diverse *L. iners* strains (Figure S10C). Taken together, these results suggest that OA may additionally improve existing BV treatment via inhibition of key BV-associated bacteria, including MTZ-resistant strains, without any expected detriment to *L. crispatus*.

**Figure 7.**
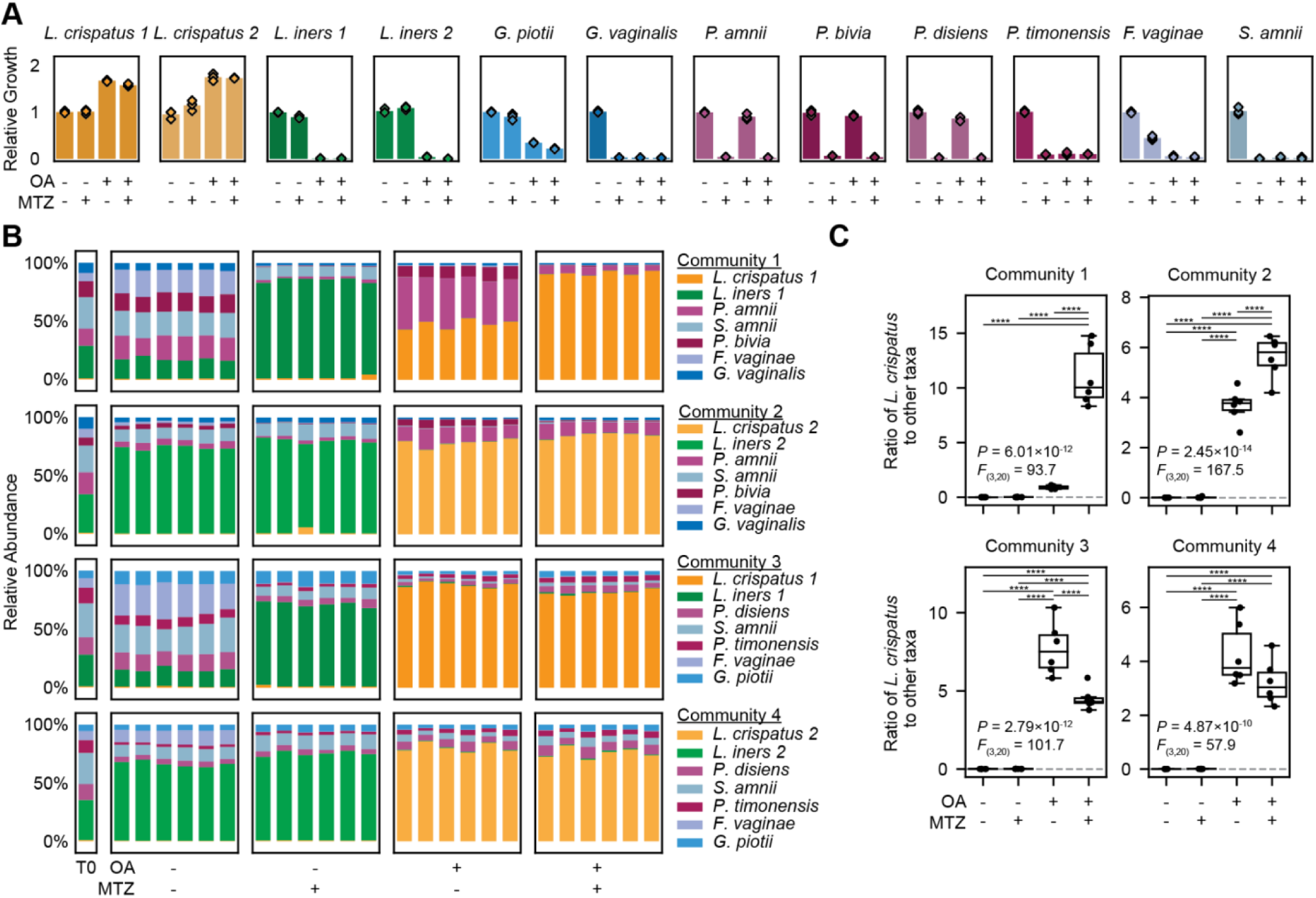
OA treatment shifts *in vitro* BV-like communities towards *L. crispatus*-dominance alone or in combination with MTZ. (A) Relative growth of the indicated bacterial species in NYCIII broth supplemented with or without metronidazole (MTZ; 50 µg/mL) and/or OA (3.2 mM) after 72 hours of culture. Growth was measured by OD600. Relative growth was calculated relative to the median OD600 measurement in the no OA supplementation control. Plotted points represent 3 technical replicates per condition. (B) Relative bacterial abundance in 4 representative, defined BV-like communities grown for 72 hours in NYCIII broth with or without MTZ (50 µg/mL) and/or OA (3.2 mM). Composition of the cultured communities and of the input mixture (T0) was determined by bacterial 16S rRNA gene sequencing. Plots depict 6 technical replicates per condition. (C) Ratios of *L. crispatus* to the sum of all other taxa in the mock communities shown in (B). The gray dotted line represents the input ratio measured in the input sample (T0). Between-group differences were determined by one-way ANOVA with post-hoc Tukey’s test; selected significant pairwise differences are shown (****p < 0.0001; full statistical results in Table S7).

### OA treatment shifts *in vitro* BV-like bacterial communities towards *L. crispatus* dominance alone or in combination with MTZ

The effects of OA on both FGT lactobacilli and BV-associated bacteria suggested it could have therapeutic potential to shift vaginal bacterial communities towards *L. crispatus* dominance, alone or in combination with MTZ. We tested this hypothesis using an established approach employing defined, BV-like bacterial communities *in vitro*^43^. The input communities, which included a predominance of BV-associated bacteria, *L. iners* at lower abundance, and *L. crispatus* at very low abundance (<2%), were cultured in rich, non-selective media with or without addition of OA and/or MTZ for 72 hours (Figure 7B). Composition of the input mixtures and the resulting *in vitro* communities was assessed by 16S rRNA gene sequencing, and total community growth was confirmed by optical density (Figures S11A-C). Untreated communities maintained diverse, BV-like compositions, while communities treated with MTZ alone became dominated by *L. iners*, recapitulating patterns observed in human studies of BV treatment^21,25–28^ (Figure 7B). By contrast, treatment with OA alone significantly promoted *L. crispatus* dominance and suppressed *L. iners* and most BV-associated species, although some *Prevotella* species were not inhibited. Combining OA with MTZ generally preserved or enhanced *L. crispatus* dominance relative to OA alone, while decreasing *Prevotella* abundance. Similar results were observed using defined, BV-like communities containing *L. gasseri* or *L. jensenii* instead of *L. crispatus*, and in experiments culturing these same communities in S-broth (Figures S12A-B). Thus, OA showed robust ability to enhance microbiota dominance by *L. crispatus* and other non-*iners* FGT *Lactobacillus* species alone or combined with MTZ in this model, highlighting its potential to improve existing BV therapies.

## Discussion

In this study, we described important differences in fatty acid metabolism among FGT *Lactobacillus* species that reveal fundamental principles of nutrient utilization in the FGT microbiota and point to new strategies for improving BV treatment. Specifically, we found that *cis-* 9-uLCFAs, exemplified by OA, selectively inhibit *L. iners* and key BV-associated species while robustly promoting growth of *L. crispatus* and other health-associated non-*iners* FGT lactobacilli. Transcriptional profiling of non-*iners* FGT lactobacilli revealed a core set of genes upregulated in response to OA and other *cis-*9-uLCFAs, including a predicted oleate hydratase enzyme and putative fatty acid efflux pump. These OA-induced genes were genomically conserved in non-*iners* FGT *Lactobacillus* species but universally absent in *L. iners*, mirroring the observed growth phenotypes. We characterized the predicted *Lactobacillus* OhyA orthologs, confirming they exhibited OhyA activity by hydrating uLCFA *cis-*9 double bonds *in vitro*. We further showed that their biochemical products were uniquely elevated in cervicovaginal fluid of women with *L. crispatus*-dominated microbiota. Notably, we overcame the historical technical challenge of conducting genetic experiments in FGT lactobacilli to assess the roles of *farE* and *ohyA* in mediating these observed growth phenotypes. Knocking out *farE* abolished the *cis-*9-uLCFA resistance phenotype, which could be restored by genetic complementation. We further showed that FGT lactobacilli are fatty acid auxotrophs, but only *farE*-harboring species could withstand and utilize high levels of OA for phospholipid synthesis. Additionally, we found that non-*iners* FGT *Lactobacillus* species can employ OhyA to sequester exogenous OA in a derivative form that only they can exploit, demonstrating an important nutrient competition strategy for these fatty acid auxotrophic species. Finally, we demonstrated that OA treatment alone or in combination with MTZ could robustly shift defined BV-like bacterial communities towards *L. crispatus*-dominated states. Together, these results advance our mechanistic understanding of FGT *Lactobacillus* metabolism and identify novel therapeutic strategies to modify the vaginal microbiota.

Our description of a role for uLCFAs in selectively inhibiting *L. iners* adds to existing knowledge about antimicrobial properties of uLCFAs and their modes of action in mammalian-adapted bacteria^38^. Antimicrobial properties of host-produced free fatty acids contribute to innate defense mechanisms on the human skin, in oral mucosa, and likely other mucosal surfaces^40,60,61^. The potency and mechanisms of antibacterial activity of these fatty acids vary across bacterial strains and species. For example, POA and sapienic acid potently inhibit *S. aureus* via physical disruption of its cell membrane, leading to leakage of low molecular weight proteins and solutes and subsequent interference with energy metabolism^39^. Similarly, sapienic acid and lauric acid physically disrupt the membranes of *Porphyromonas gingivalis* and lead to cell lysis^62^. In contrast, high concentrations of OA and LOA are reported to induce bacteriostatic inhibition in certain *S. aureus* strains by inhibiting the FASII enzyme, FabI^63^. Based on the rapid lysis induced by OA in *L. iners*, we propose that its bactericidal action results from a membrane-disrupting mechanism. We speculate that *L. iners*’ inability to prevent toxic intracellular accumulation of these fatty acids and regulate their membrane insertion contributes to its *cis*-9-uLCFA susceptibility.

Our identification of a conserved *cis*-9-uLCFA-induced response mechanism in non-*iners* FGT lactobacilli is consistent with uLCFA resistance mechanisms previously described in other organisms. For example, *S. aureus* FarE was reported to confer resistance to LOA and arachidonic acid via uLCFA efflux^64^ and Tet38, a separate *S. aureus* fatty acid efflux pump, promotes resistance to POA^65^. A functionally similar MtrCDE efflux system in *Neisseria gonorrhoeae* exports hydrophobic molecules, such as LCFAs, and enhances survival in fatty acid enriched environments, including the genital tract of a female mouse model^66^. Thus, our finding that FarE mediates *cis*-9-uLCFA resistance in non-*iners* FGT lactobacilli, agrees with this larger body of literature showing that fatty acid efflux pumps serve as an important resistance mechanism against exogenous fatty acids and can confer a competitive advantage in LCFA-rich environments.

Interestingly, knocking out *ohyA9* in *L. gasseri* did not increase *cis*-9-uLCFA susceptibility in our *in vitro* experimental conditions, contrasting with findings in *S. aureus* and *S. pyogenes*, where OhyA was protective against POA toxicity^49,52^. Thus, the role of OhyA in protecting bacteria against uLCFAs may be specific to certain species or environmental contexts. In addition to protecting against uLCFA toxicity, OhyA activity may play a role in tolerogenic host signaling through *h*FA products as shown for *S. aureus* infections^56^ or in producing products inhibitory to other bacteria or fungi^67^. Future studies will be required to determine how OhyA enzymes contribute to host-microbe crosstalk and competitive microbial chemical warfare.

Our finding that FGT *Lactobacillus* species are fatty acid auxotrophs offers insight into a previously underappreciated nutrient requirement and their metabolic niche in the FGT microbiome. These results agree with previous reports of closely related non-FGT *Lactobacillus* species (*Lactobacillus johnsonii, Lactobacillus helveticus, and L. acidophilus*) being fatty acid auxotrophs as well^68,69^. Such findings imply that many *Lactobacillus* species rely on fatty acids derived from their environment, host, or other microbial species. A critical next step will be understanding which specific sources within the FGT support *Lactobacillus* species growth *in vivo*. Based on our finding that *ohyA9* confers a growth advantage to non-*iners Lactobacillus* species in the presence of 10-HSA, we propose that these species employ OhyA9 to sequester environmental *cis*-9-uLCFAs in a derivative form inaccessible to non-*ohyA9*-harboring organisms in the FGT microbiota. This proposed strategy aligns with previously described competitive microbial mechanisms for molecularly encrypting limited nutrients to prevent access by competing species^70^. For example, bacteria produce species-specific siderophores to bind and sequester environmental metals^71,72^, rendering them inaccessible to organisms lacking the cognate siderophore receptor. Siderophores also serve as only a temporary molecular modification of the acquired nutrient, which is shed following entry into the bacterial host^71,72^. Corrinoid cofactors are another example of molecular nutrient encryption, with their diverse structures dictating which bacteria can take up and utilize the cofactor with corrinoid-compatible transporters and enzymes^73,74^. We propose that OhyA9 activity in FGT lactobacilli is another example of the diverse mechanisms bacteria employ to acquire, sequester, and utilize limited nutrients in competition with other species.

Finally, we demonstrated the therapeutic potential of OA to improve upon standard treatment for BV. Promotion of *L. crispatus* over *L. iners* in the FGT microbiota is a core objective of BV treatment for which novel approaches are needed. Methods to target MTZ-resistant BV-associated bacteria may also be important, although the extent to which MTZ-resistant bacteria contribute to BV recurrence remains unclear^29–31^. We find that – in addition to inhibiting *L. iners* growth – OA also inhibits several BV-associated species, including MTZ-resistant *G. piotii* and *F. vaginae* strains. Of particular interest is OA-driven inhibition of *F. vaginae*, a frequently MTZ-resistant species that has been linked to BV treatment failure^31^. We found that treatment of defined, *in vitro* BV-like communities with OA (alone or in combination with MTZ) strongly promoted *L. crispatus* dominance while suppressing both *L. iners* and BV-associated anaerobes, such as *Gardnerella* species, *P. timonensis*, *F. vaginae*, and *S. amnii*. Current investigational therapies for BV include vaginal microbiome transplants and single-or multi-strain *Lactobacillus* live biotherapeutic products that aim to shift microbiota composition by introducing health-associated species^24,32,33^. However, preliminary clinical efficacy has been modest. We propose that supplementing delivery of a *L. crispatus*-containing live biotherapeutic and MTZ treatment with OA or other uLCFAs could improve efficacy by promoting *L. crispatus* colonization and inhibiting competition from *L. iners* and BV-associated bacteria.

In summary, we identified and characterized important species-level differences in fatty acid resistance, metabolism, and nutrient sequestration mechanisms among FGT *Lactobacillus* species, demonstrated *in vivo* relevance of these mechanisms, and provide preclinical evidence to support translating the findings into a novel therapeutic strategy for BV. This study illustrates how functional, metabolic, and genomic approaches can inform development of microbiota-targeted therapies to improve human health.

## Limitations of the study

While this study provides mechanistic and functional support for its primary findings, the work has several limitations. Genetic manipulation experiments were performed only in *L. gasseri* due to lack of accessible genetic tools for other FGT *Lactobacillus* species. Genetic manipulation of these organisms (including *L. gasseri*) has been a major challenge for the field^75–79^, but we succeeded in generating and complementing the necessary mutants in *L. gasseri*, which closely resembles *L. crispatus* in genomic, phylogenetic, transcriptional, enzymatic, and phenotypic features relevant to uLCFA resistance. Similarly, lack of genetic tools for manipulating *L. iners* prevented us from performing gain-of-function genetic experiments in *L. iners* to heterologously express uLCFA-induced genes from non-*iners* lactobacilli^80^. Finally, we employed a previously described *in vitro* model of BV^43^ because animal models (including gnotobiotic mice and non-human primates) have been largely intractable to vaginal colonization by human FGT lactobacilli or fully diverse BV-like communities^81–84^, precluding their use for preclinical testing of uLCFAs *in vivo*. Human trials will likely be needed to fully assess the effects of uLCFA treatments on relevant FGT bacterial species and communities.

## Acknowledgements

We dedicate this manuscript to the memory of Dr. Charles O. Rock, who made significant contributions to the field of bacterial lipid biology and the work described in this paper. Additionally, we would like to thank the FRESH study participants for donating clinical samples used in this study; study staff at the FRESH study and laboratory staff at the HIV Pathogenesis Programme at UKZN for sample processing; B. Bowman for assistance with 16s rRNA gene sequencing on human vaginal swab samples; J. A. Elsherbini (Ragon Institute) for bioinformatics guidance and scientific discussion; R. T. Walton (MIT) and J. Chen (MIT) for genetics guidance; K. Miller (St. Jude’s Children’s Research Hospital) for technical assistance on the enzyme characterization work; C. N. Tzouanas (MIT) and S. Goldman (MIT) for scientific discussion and manuscript review; R. Majovski (Broad Institute) for manuscript review; and the Vaginal Microbiome Research Consortium for valuable feedback and discussion on the work. M.Z. and M.W.T. were supported by the National Science Foundation Graduate Research Fellowship under Grant No. 1745302; S.M.B. by the NIH Grant No. 1K08AI171166; C.D.R. by the NIH Grant No. 4R00AI166116; F.A.H. by the Schmidt Science Fellowship; P.C.B., M.Z., and M.W.T. by the NIH NIAID Grant No. 5U19AI42780; C.O.R. in part by the American Lebanese Syrian Associated Charities (ALSAC) and St. Jude Children’s Research Hospital; and, D.S.K, S.M.B., and M.Z. by grants from the Bill & Melinda Gates Foundation.

EC1000 was a gift from Todd Klaenhammer & Jan Kok (Addgene plasmid #71852). pORI28 was a gift from Todd Klaenhammer & Jan Kok (Addgene plasmid #71595; http://n2t.net/addgene:71595; RRID:Addgene_71595). pTRK892 was a gift from Rodolphe Barrangou & Todd Klaenhammer (Addgene plasmid #71803; http://n2t.net/addgene:71803; RRID:Addgene_71803). pTRK669 was a gift from Rodolphe Barrangou & Todd Klaenhammer (Addgene plasmid #71313; http://n2t.net/addgene:71313; RRID:Addgene_71313). Bulk RNA-sequencing samples were processed and data were generated by the Infectious Disease and Microbiome Program’s Microbial Omics Core at the Broad Institute of MIT and Harvard. Electron Microscopy Imaging, consultation and services were performed in the HMS Electron Microscopy Facility. DNA sequencing of plasmids constructed for the complemented *S. aureus* strains was performed by St. Jude Children’s Research Hospital’s Hartwell Center for Biotechnology.

## Supplemental Figures

**Figure S1.**
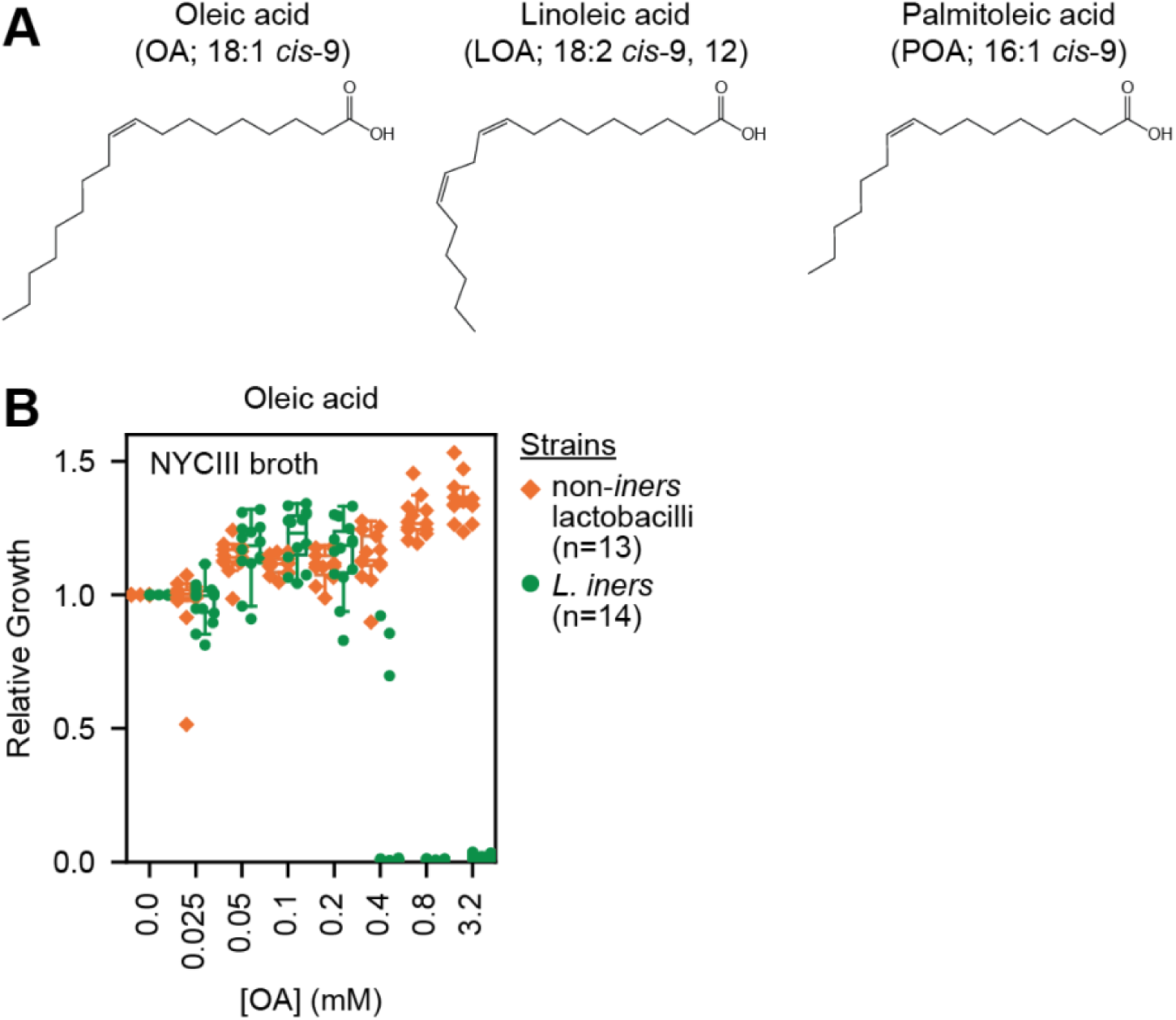
*cis*-9-uLCFA structures and *L. iners* selective inhibition in NYCIII broth. (A) Chemical structures of oleic acid, linoleic acid, and palmitoleic acid. (B) Relative growth of diverse non-*iners* FGT *Lactobacillus* (n=13) and *L. iners* (n=14) strains in NYCIII broth supplemented with varying concentrations of OA. Growth was measured by OD600 after 72 hours of culture. Relative growth was calculated relative to the median OD600 measurement in the no fatty acid supplementation control. Plotted points represent the median relative growth for 3 technical replicates per condition.

**Figure S2.**
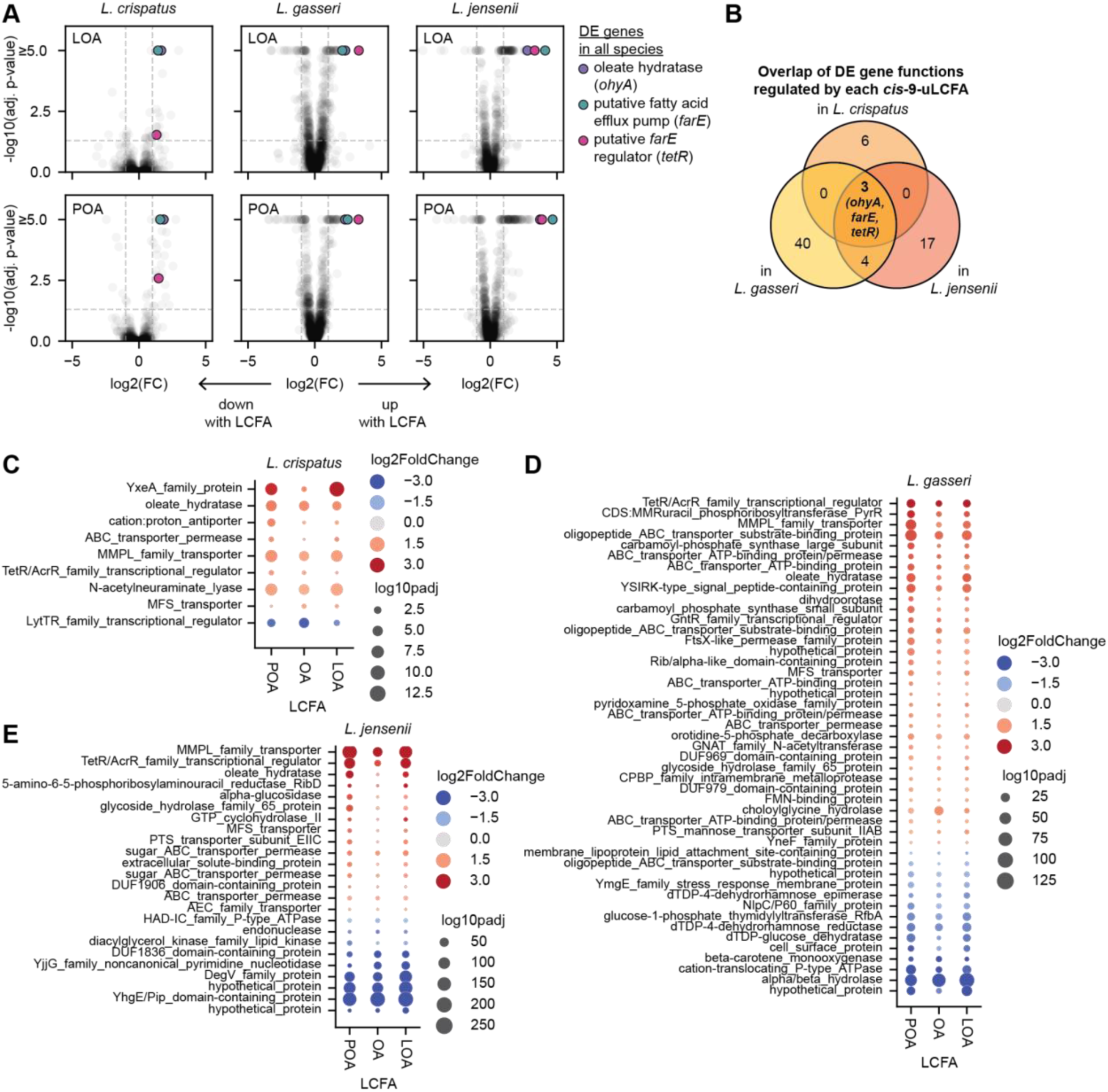
*cis-*9-uLCFAs upregulate a conserved set of genes in non-*iners* FGT lactobacilli. (A) Transcriptomic responses of cultured *L. crispatus* (left), *L. gasseri* (middle), and *L. jensenii* (right) that were grown to exponential phase in MRS+CQ broth, then exposed to LOA (top, 3.2 mM), or POA (bottom, 3.2 mM) for 1 hour. Data were analyzed as in Figure 2A. Consistently DE genes included a predicted *ohyA* (purple), putative *farE* (teal), and its putative *tetR* (pink). The untreated controls for each species shown in Figures 2A and S2A are the same. (B) Venn diagram showing the sets of genes differentially expressed in response to all three *cis*-9-uLCFA treatments (OA, LOA, and POA) in each species and the overlap between these sets of DE gene functions that were shared among all three species. Only three shared DE gene functions (*ohyA*, *farE*, and *tetR*; all up-regulated) were observed in all species under all treatment conditions. (C-E) Dot plot showing the shared sets of DE genes induced by OA, LOA and POA treatments in *L. crispatus* (C), *L. gasseri* (D), and *L. jensenii* (E). Each point depicts a gene with color representing log2(fold change, FC) and size representing -log10(adjusted p-value).

**Figure S3.**
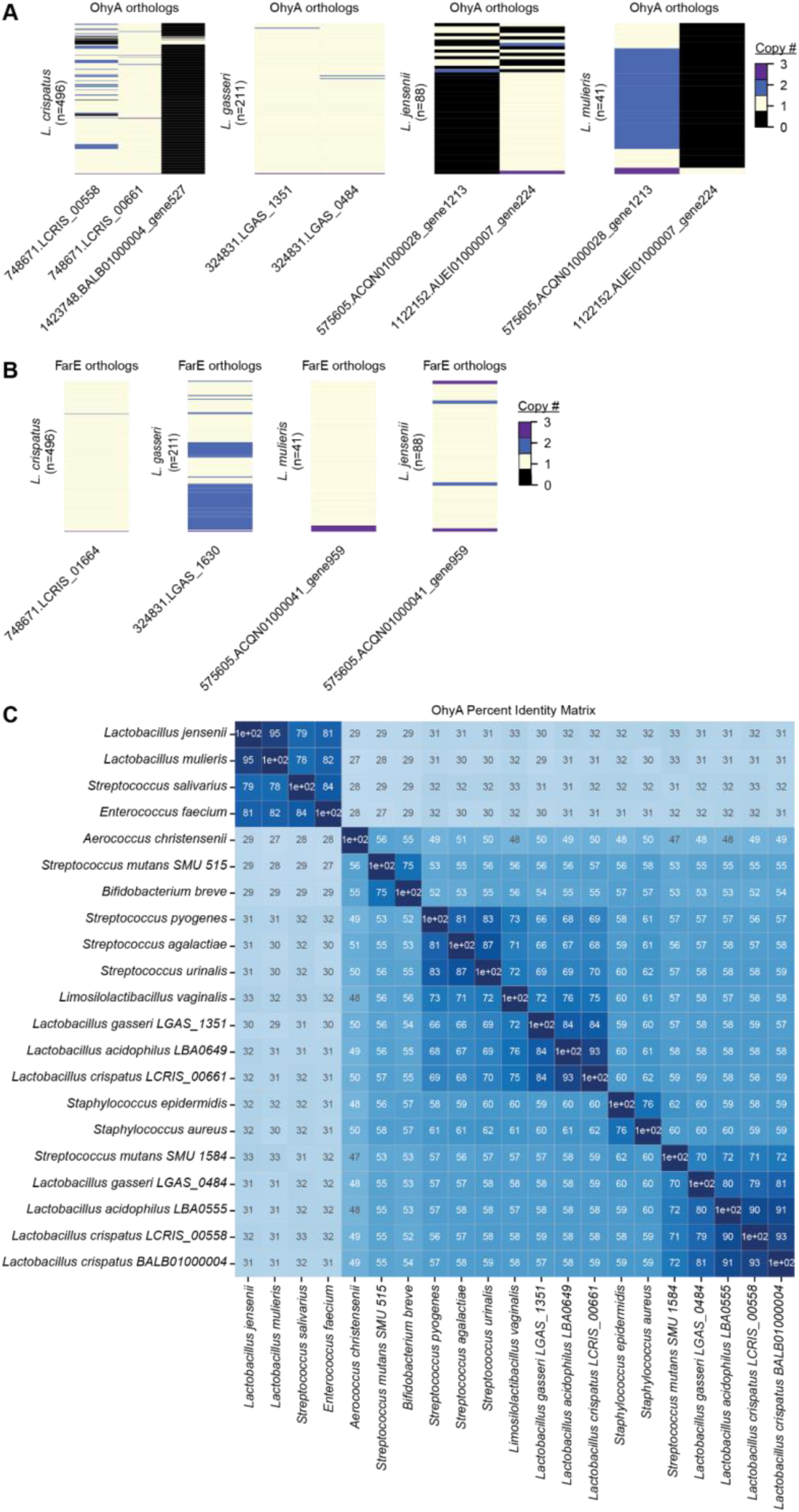
OhyA and FarE ortholog diversity in FGT *Lactobacillus* species and OhyA homology to other human-adapted species. (A-B) EggNOG-predicted orthologs for OhyA (A) and FarE (B), and their respective ortholog copy numbers per genome and MAG for each species. Number of genomes and MAGs for each species are shown. (C) Percent identity matrix (PIM) of representative OhyA orthologs in FGT *Lactobacillus* species and other human-adapted bacteria, determined by protein sequence alignment (MUSCLE v5.1^87^).

**Figure S4.**
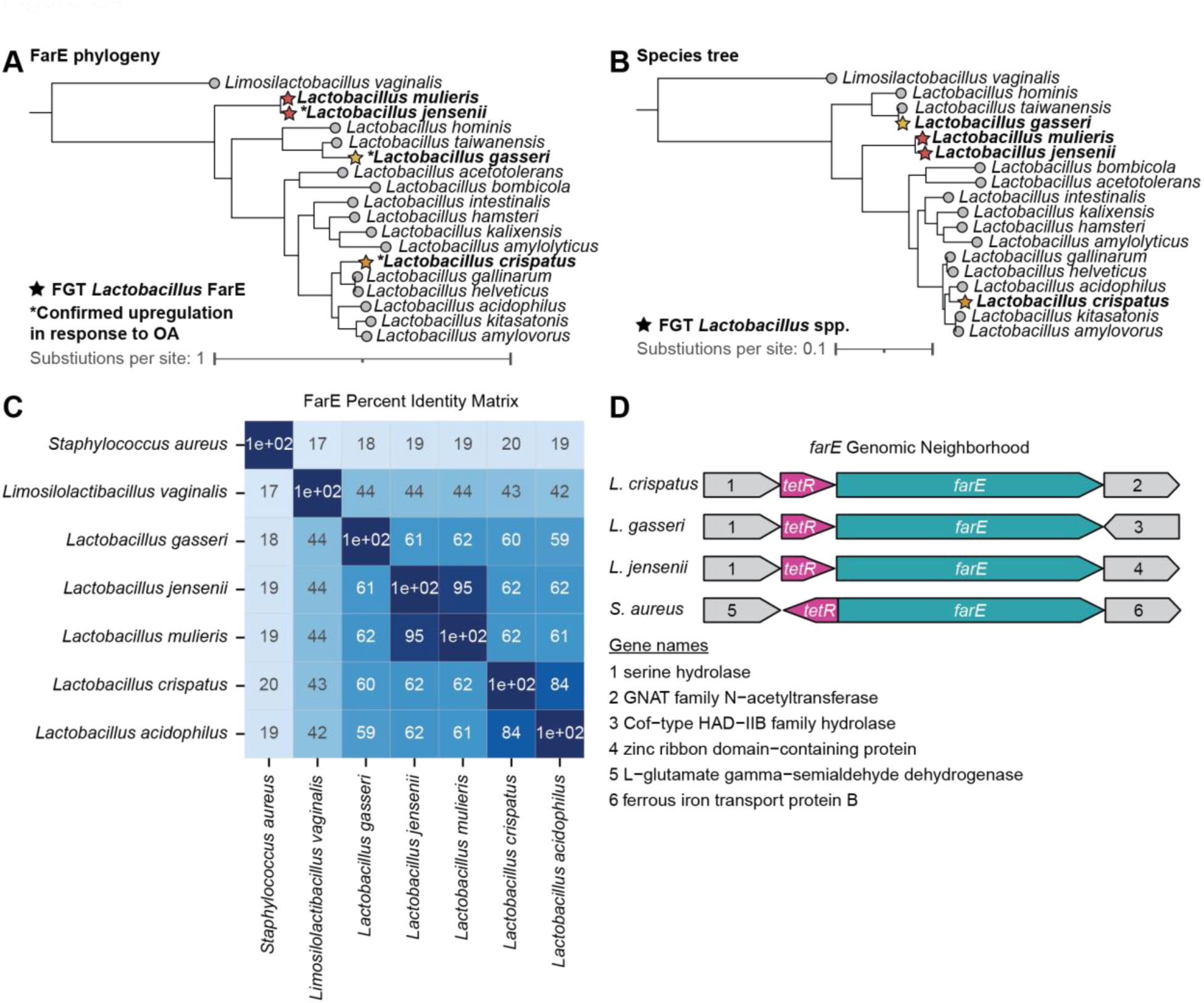
FarE ortholog phylogeny in *Lactobacillus* species and homology to previously characterized FarE in *S. aureus*. (A) FarE protein phylogenetic tree for representative *farE* orthologs from the indicated *Lactobacillus* species. Starred leaf tips correspond to FGT *Lactobacillus* orthologs shown in bold. FarE orthologs upregulated in response to OA (Figure 2A) are marked with asterisks. (B) Species phylogenetic tree of representative *Lactobacillus* genomes constructed based on core ribosomal protein sequences. Starred leaf tips correspond to FGT *Lactobacillus* species shown in bold. (C) PIM of representative EggNOG-predicted FarE orthologs in FGT *Lactobacillus* species and other human-adapted bacteria, determined by protein sequence alignment (MUSCLE v5.1^87^). (D) Genomic neighborhood of *farE* in representative *L. crispatus*, *L. gasseri*, *L. jensenii*, and *S. aureus* genomes. (A and B) Trees were constructed from MUSCLE v5.1^87^-aligned protein sequences using RAxML-NG^88^ (see methods).

**Figure S5.**
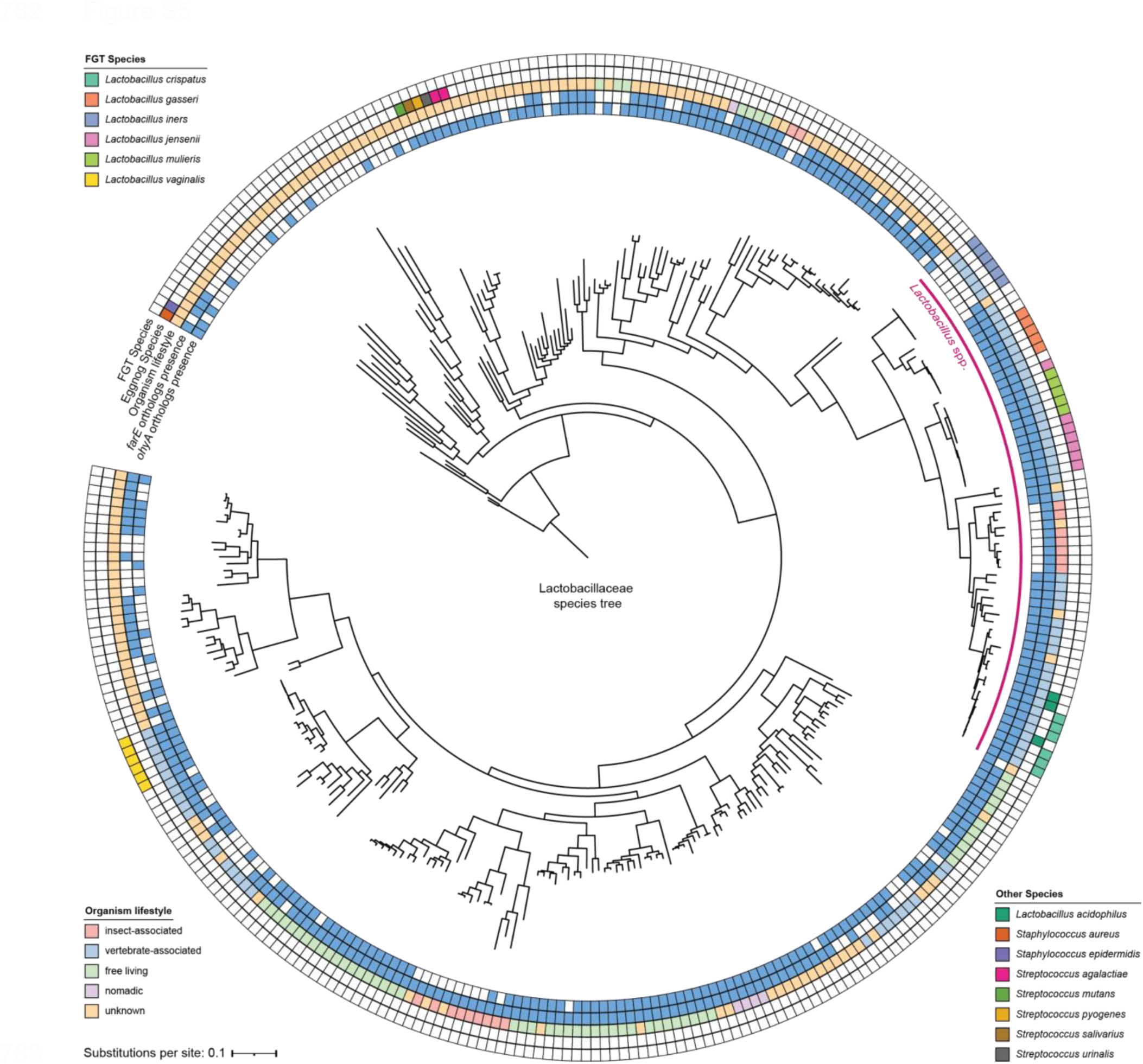
*ohyA* and *farE* orthologs presence is widespread among Lactobacillaceae species. Species phylogenetic tree of representative species genomes from the Lactobacillaceae family, constructed based on core ribosomal genes contained in all species. The species genomes derive from a recent comprehensive review and taxonomic revision of the 48 genera within the Lactobacillaceae family^55^. The tree also includes *Staphylococcus aureus* and *Staphylococcus epidermidis* genomes to root the phylogenetic reconstruction. Metadata rings mark the genomes of FGT lactobacilli species, genomes of other Gram positive host-adapted species, each organism’s lifestyle if known, and the presence of *farE* and *ohyA* orthologs in each represented genome. Trees were constructed from MUSCLE v5.1^87^-aligned protein sequences using FastTree v2.1 (see methods).

**Figure S6.**
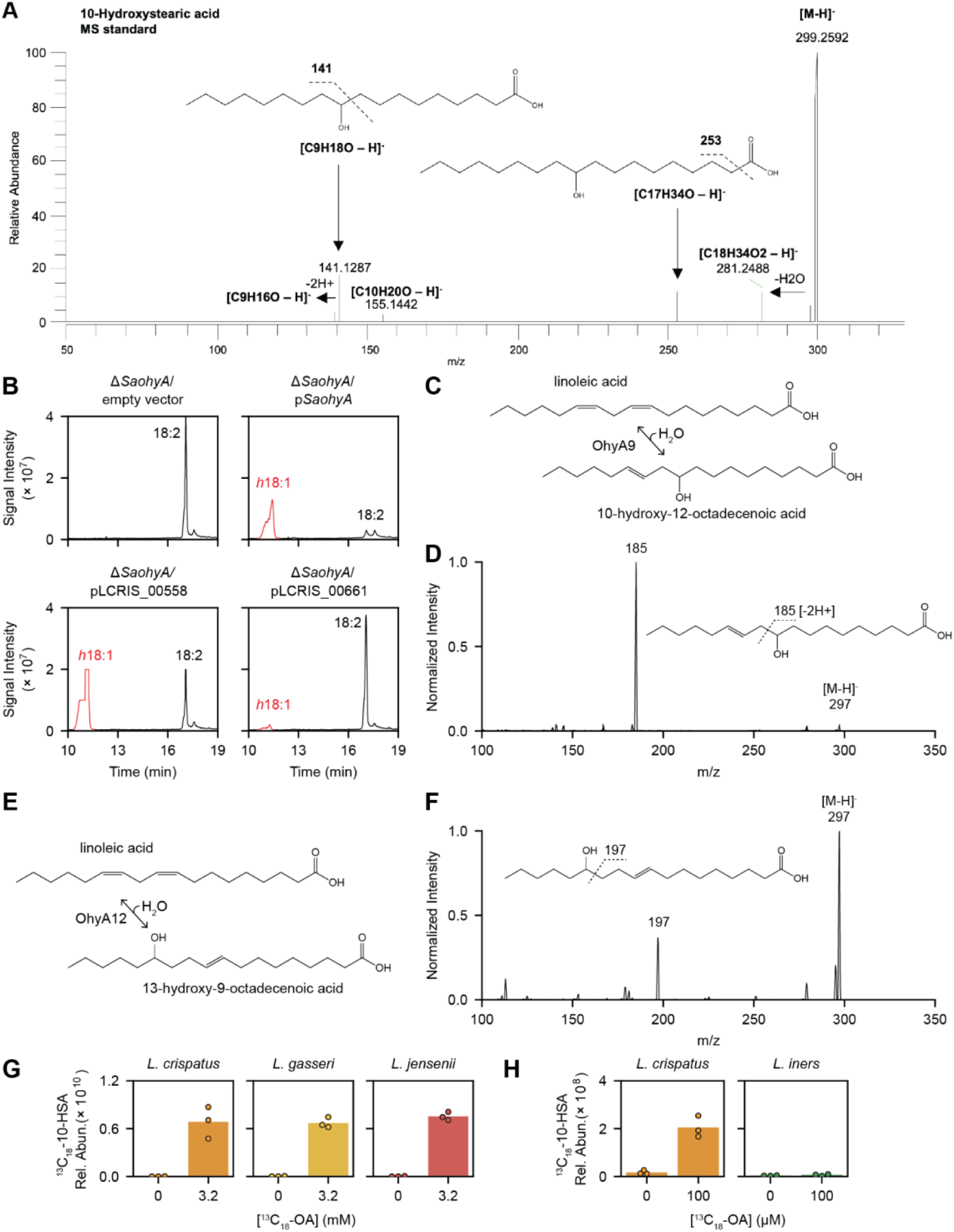
Detection and identification of enzymatic products from *ohyA* orthologs found in non-*iners* FGT *Lactobacillus* genomes. (A) MS2 spectra with major fragmentation labels for 10-HSA standard (Ambeed, A125712-50MG). (B) Extracted ion chromatograms from supernatants of *ΔSaohyA*/empty vector, *ΔSaohyA*/p*SaohyA*, *ΔSaohyA*/pLCRIS_00558, and *ΔSaohyA*/pLCRIS_00661 cultured with LOA for 1 hour. Annotated peaks include LOA (18:2) and the detected hydroxyFA (*h*18:1). (C) OhyA9 enzymatic activity reaction diagram with LOA substrate. (D) MS2 spectra with major fragmentation labels for the *h*18:1 peak from the supernatant of *ΔSaohyA*/pLCRIS_00661 cultured with LOA and shown in the lower right panel of Figure S6B, identified as 10-hydroxy-12-octadecenoic acid (*h*18:1). (E) OhyA12 enzymatic activity reaction diagram with LOA substrate. (F) MS2 spectra with major fragmentation labels for the *h*18:1 peak from the supernatant of *ΔSaohyA*/pLCRIS_00558 cultured with LOA and shown the lower left panel of Figure S6B, identified as 13-hydroxy-9-octadecenoic acid. (G) Detection of ^13^C_18_-10-HSA in cell pellets of *L. crispatus*, *L. gasseri*, and *L. jensenii* cultured for 72 hours in NYCIII broth with and without ^13^C_18_-OA (3.2 mM, which is a lethal concentration for *L. iners*). The cell pellets are from the same cultures as the supernatants shown in Figure 3E. (H) Detection of ^13^C_18_-10-HSA in cell pellets of *L. crispatus* and *L. iners* cultured for 72 hours in NYCIII broth with and without ^13^C_18_-OA (100 µM, which is a sublethal concentration for *L. iners*). The cell pellets are from the same cultures as the supernatants shown in Figure 3F. (G and H) Plotted points represent 3 technical replicates per condition.

**Figure S7.**
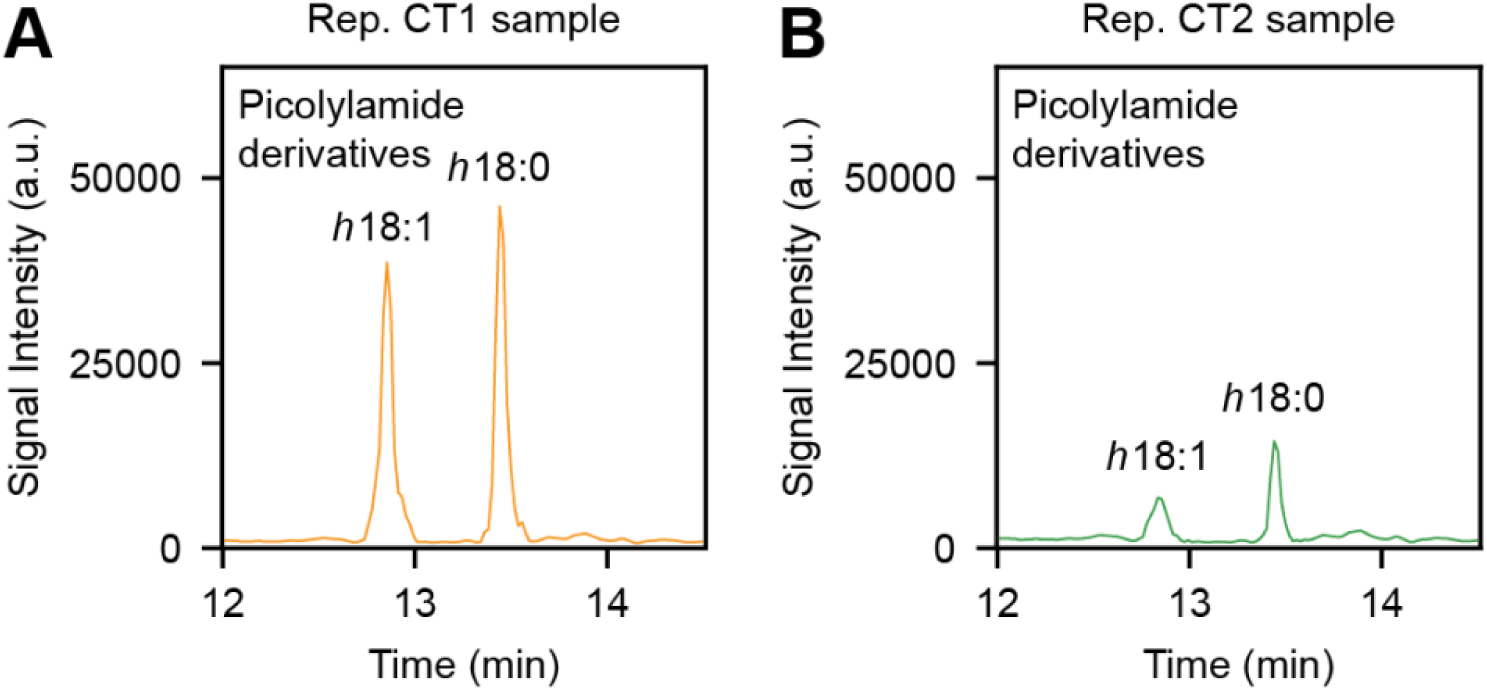
Representative chromatograms for CT1 and CT2 human CVL samples for the detection of *h*FAs. (A-B) Extracted ion chromatogram of picolylamide derivatized *h*FAs recovered from a representative CVL sample from a participant with a CT1 microbiome composition (A) and from a participant with a CT2 microbiome composition (B).

**Figure S8.**
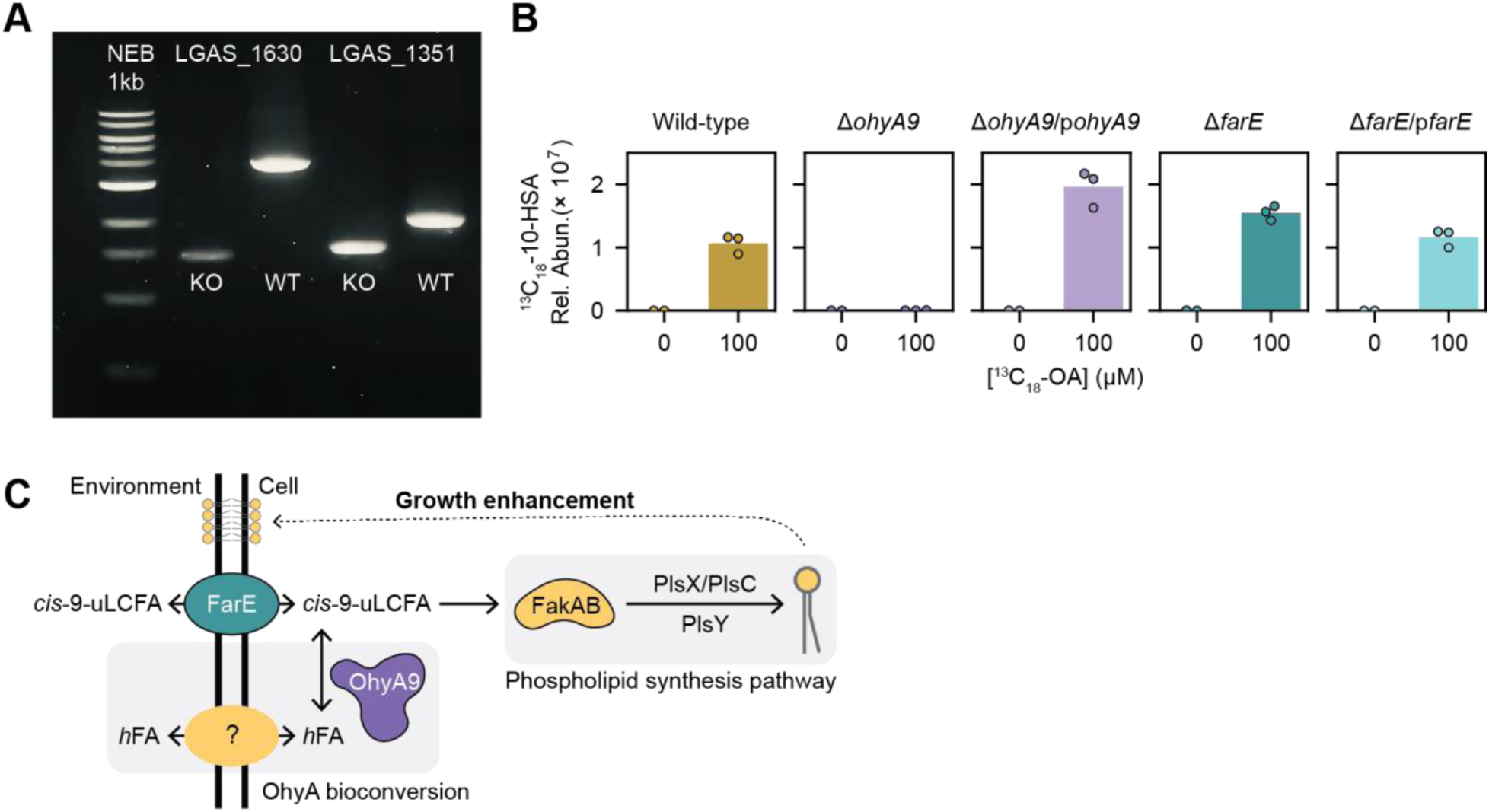
Production and characterization of *ohyA9* and *farE* genetic knockouts in *L. gasseri*. (A) DNA gel of PCR products amplified from the primers flanking either LGAS_1630 (*farE*) or LGAS_1351 (*ohyA9*) from the *L. gasseri* ATCC 33323 wild-type (WT) strain and *ΔohyA9* and *ΔfarE* genetic knockout (KO) strains. Expected amplicon lengths were 3.9kb for LGAS_1630 in WT, 2.0kb for LGAS_1351 in WT, and 1.2kb for each in-frame gene deletion of LGAS_1351 in *ΔohyA9* and of LGAS_1630 in *ΔfarE*. (B) Detection of ^13^C_18_-10-HSA in cell pellets from *L. gasseri* WT, *ΔohyA9*, *ΔohyA9*/p*ohyA9*, *ΔfarE*, and *ΔfarE*/p*farE* cultured for 24 hours in NYCIII broth with and without ^13^C_18_-OA (100 µM). The cell pellets are from the same cultures as the supernatants show in Figure 5D. Plotted points represent 2 or 3 technical replicates per condition. (C) Schematic depicting a proposed model for FarE and OhyA9 activity with exogenous *cis*-9-uLCFAs in non-*iners* FGT *Lactobacillus* species.

**Figure S9.**
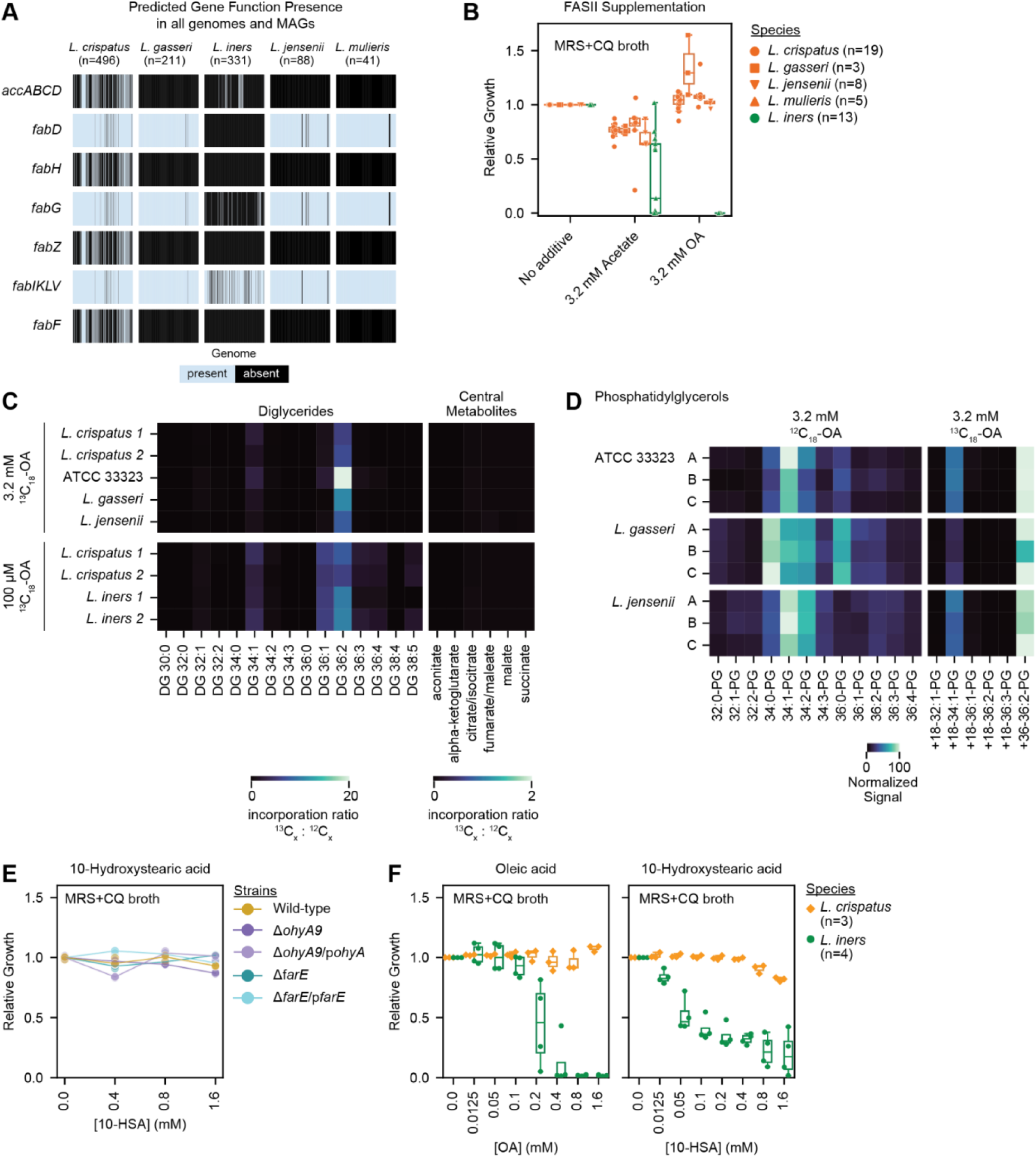
Genomic analysis of FASII pathway in FGT *Lactobacillus* genomes, OA isotope tracing in cultured FGT lactobacilli, and 10-HSA growth effects. (A) Distribution of genes predicted to encode FASII pathway genes in isolate genomes and MAGs of the indicated FGT *Lactobacillus* species (n=1,167). Gene function was predicted using eggNOG 5.0^85^ employing eggNOG-mapper v2.1.9^86^. (B) Relative growth of diverse *L. crispatus* (n=19), *L. gasseri* (n=3), *L. iners* (n=13), *L. jensenii* (n=8), and *L. mulieris* (n=5) strains in MRS+CQ broth supplemented with 3.2 mM acetate or 3.2 mM OA. Growth was measured by OD600 after 72 hours of culture. (C) Heatmap representing the median incorporation ratio of ^13^C_18_-OA in detected diglycerides and central metabolites involved in the TCA cycle in cell pellets from representative strains of FGT *Lactobacillus* species. Bacteria were cultured for 72 hours in NYCIII broth with 3.2 mM (top) or 100 µM (bottom). ^13^C_18_-OA incorporation ratio was calculated as the signal from the detected ^13^C-labeled metabolite relative to the signal of the detected unlabeled metabolite. (D) Heatmap representing the normalized signal of detected unlabeled and labeled phosphatidylglycerol in cell pellets from representative strains of FGT *Lactobacillus* species. Bacteria were cultured for 72 hours in NYCIII broth with 3.2 mM ^12^C_18_-OA (left) or ^13^C_18_-OA (right). Each row labeled A, B, and C represents a technical replicate of the given condition. (E) Relative growth of *L. gasseri* ATCC 33323 WT and derivative genetic mutant strains in MRS+CQ broth supplemented with varying concentrations of 10-HSA. Growth was measured by OD600 after 24 hours of culture. (F) Relative growth of *L. crispatus* (n=3) and *L. iners* (n=4) strains in MRS+CQ broth supplemented with varying concentrations of OA (left) or 10-HSA (right). Growth was measured by OD600 after 72 hours of culture. (B, E, and F) Relative growth was calculated relative to the median OD600 measurement in the no supplementation control. (B and F) Plotted points represent the median of 3 technical replicates per condition. (E) Plotted points represent 3 technical replicates per condition.

**Figure S10.**
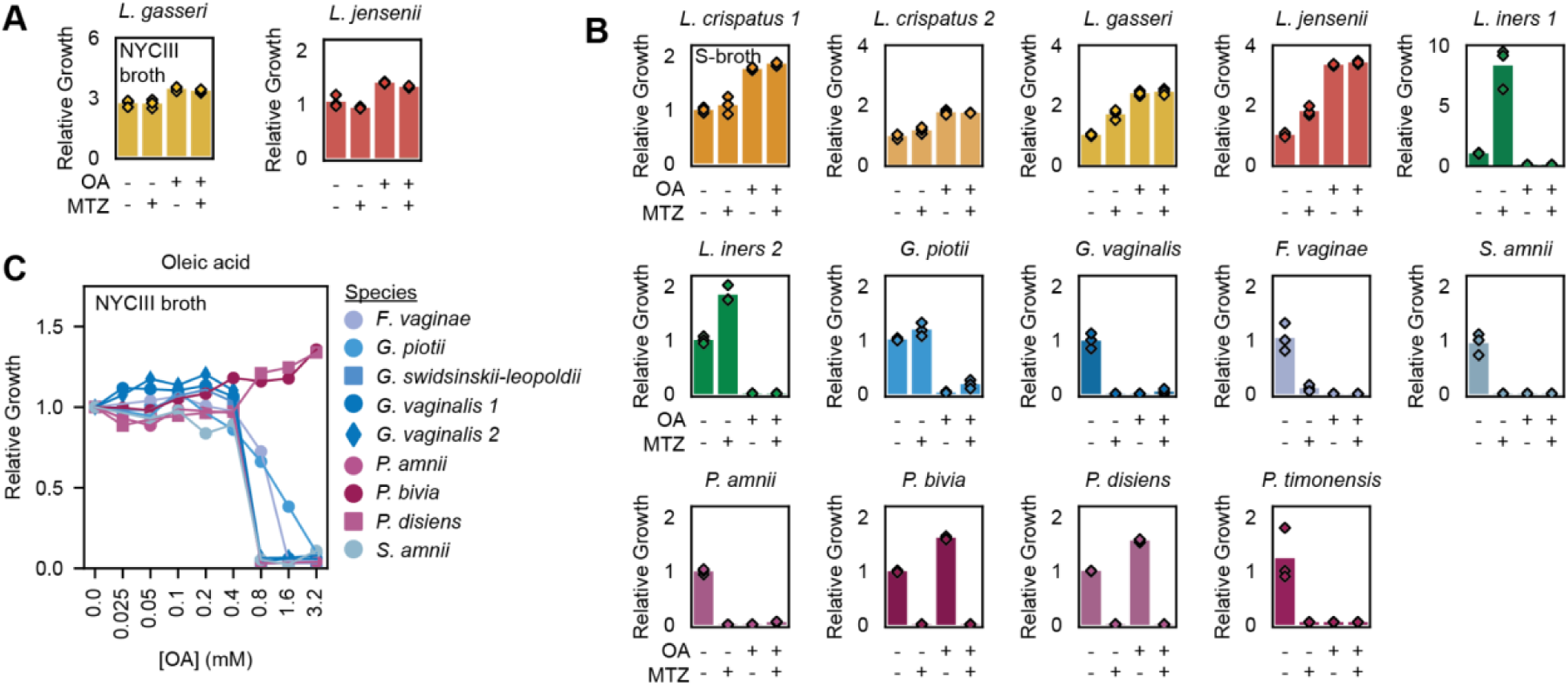
Effects of OA and MTZ on growth of FGT bacterial species. (A) Relative growth of representative *L. gasseri* and *L. jensenii* strains in NYCIII broth supplemented with or without MTZ (50 µg/mL) and/or OA (3.2 mM). (B) Relative growth of the indicated bacterial species in S-broth supplemented with or without MTZ (50 µg/mL) and/or OA (3.2 mM). (C) Relative growth of representative *F. vaginae, G. piotii, G. swindsinskii-leopoldii, G. vaginalis* (n=2), *P. amnii, P. bivia, P. disiens,* and *S. amnii* strains in NYCIII broth supplemented with varying concentrations of OA. Plotted points represent the median relative growth for 3 technical replicates per condition. (A-C) Growth was measured by OD600 after 72 hours of culture. Relative growth was calculated relative to the median OD600 measurement in the no OA supplementation control. (A-B) Plotted points represent 3 technical replicates per condition.

**Figure S11.**
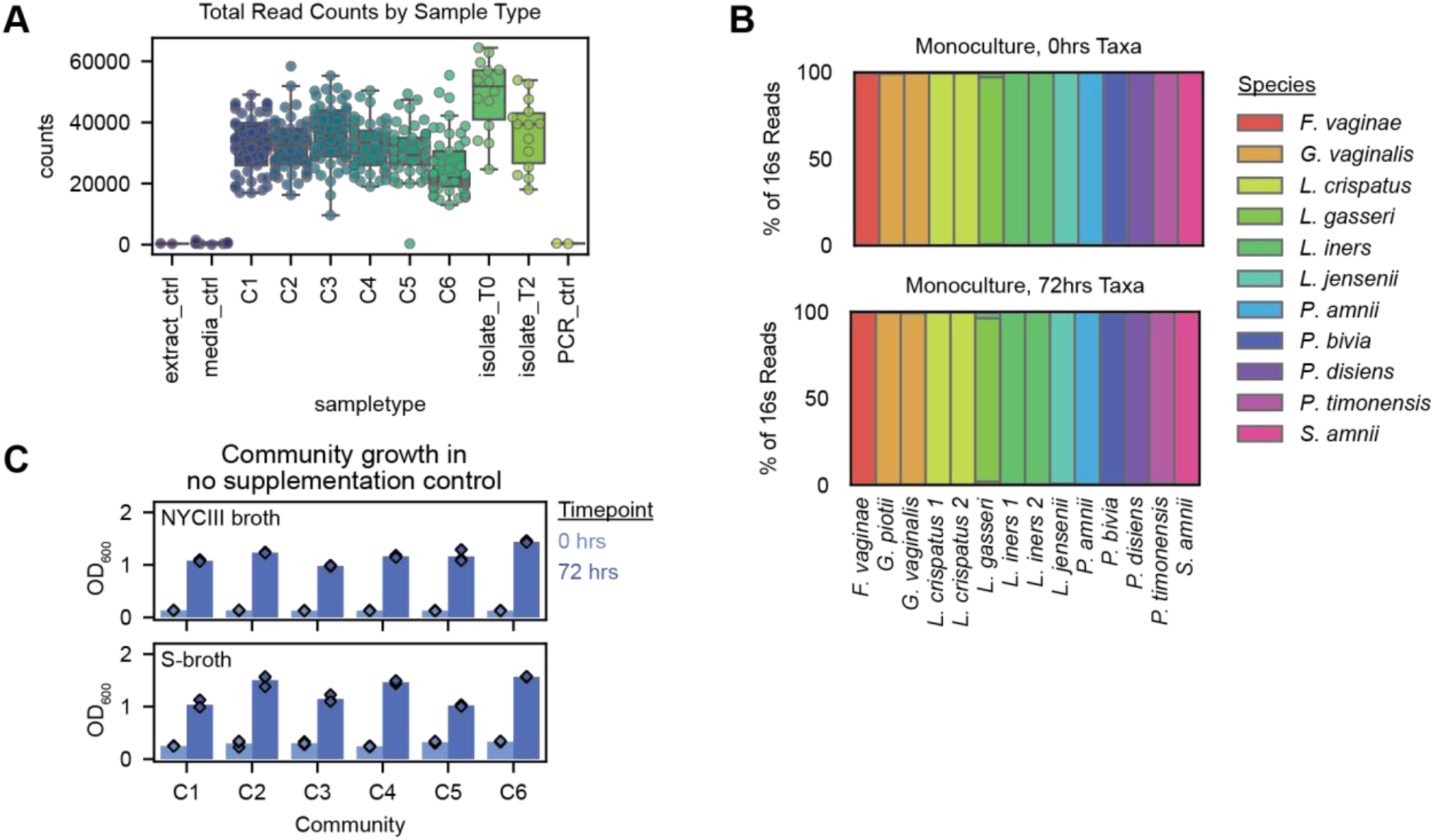
Controls for defined, *in vitro* BV-like bacterial community experiments. (A) Total sequencing read counts from mock BV-like community experiments per sample type, including extraction controls (extract_ctrl), blank media controls (media_ctrl), community samples (C1-C6), input bacterial isolate mono-cultures (isolate_T0), 72-hour cultured bacterial isolate mono-cultures (isolate_T2), and no template PCR controls (PCR_ctrl). (B) Percent of 16S rRNA gene sequencing reads for each mono-cultured bacterial isolate used to construct the defined communities. Plot shows data for input (top) isolates and for the corresponding 72-hour mono-culture controls (bottom). (C) OD600 for each total community (C1-C6) at 0 and 72 hours of culture in NYCIII broth (top) and S-broth (bottom).

**Figure S12.**
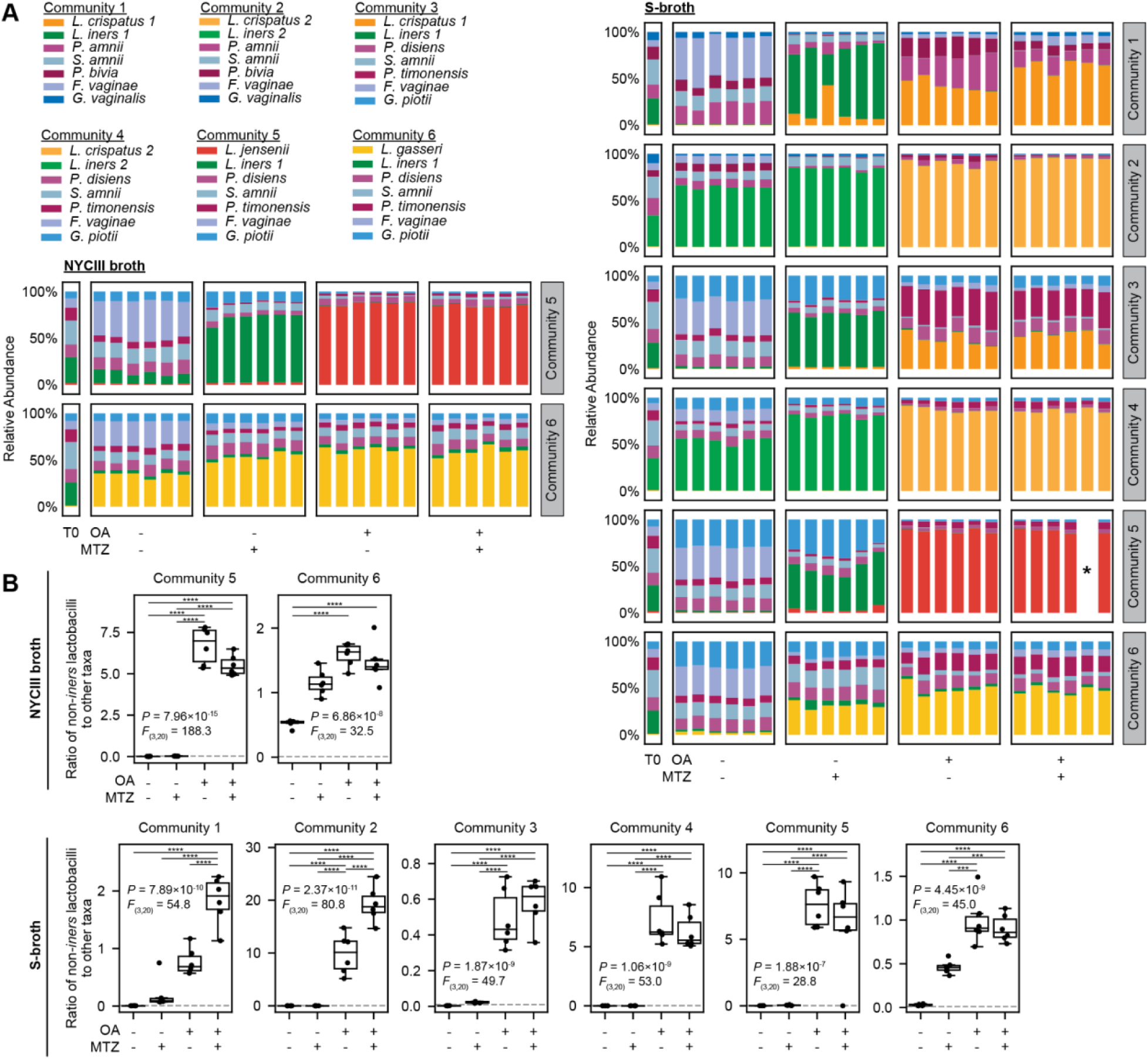
OA shifts BV-like communities towards non-*iners* FGT *Lactobacillus*-dominance in NYCIII and S-broth. (A) Relative bacterial abundance in 2 representative defined BV-like communities grown for 72 hours in NYCIII broth and S-broth with or without MTZ (50 µg/mL) and/or OA (3.2 mM). Composition of the cultured communities and of the input mixtures (T0) was determined by bacterial 16S rRNA gene sequencing. Plots depict 6 technical replicates per condition. Sequencing reads were not recovered from a single technical replicate of community 5 cultured in S-broth with MTZ and OA and is marked with an asterisk; its low read count is shown in Figure S11A. (A) Ratios of non-*iners* FGT *Lactobacillus* species taxa to the sum of all other taxa in the mock communities shown in A. The gray dotted line represents the input ratio measured in the input sample (T0). Between-group differences were determined by one-way ANOVA with post-hoc Tukey’s test; selected significant pairwise differences are shown (***p < 0.001, ****p < 0.0001; full statistical results in Table S4).

## Resource availability

### Lead contact

Further information and requests for resources and reagents should be directed to and will be fulfilled by the lead contact, Douglas S. Kwon.

### Materials availability

All unique/stable reagents directly used in this study are available from the lead contact with a completed Materials Transfer Agreement. ATCC 33323 *upp* gene deletion mutant and the empty vector, pMZ7, used for double homologous recombination will be deposited at public repositories.

## Data and code availability

### Supplemental Data

Original data will be deposited in a public repository at XX.

● Supp. Data 1. Full sized TEM images
● Supp. Data 2. RNA-sequencing data and reference genomes
● Supp. Data 3. WGS of genetic knockout strains
● Supp. Data 4. 16s rRNA gene sequencing data for competition assays
● Supp. Data 5. 16s rRNA gene sequencing data for human samples

### Supplemental Code

Original code will be deposited in a public repository at XX.

## Methods

### Bacteria strains and culture conditions

Bacterial strains used in this study for each experiment/figure are summarized in Table S1. All vaginal bacterial isolates were cultured under anaerobic conditions at 37–40°C using a palladium catalyst-based anaerobic chamber (COY) with an atmosphere of 5% carbon dioxide, 5% hydrogen, and 90% nitrogen (Airgas #X03NI90C3001054). All media, culture reagents, and plastic-ware were pre-reduced by placement overnight in the anaerobic chamber before use. Vaginal bacterial isolates were first revived on solid agar plates to obtain single colonies. After 4– 5 days of incubation, representative colonies were picked into liquid culture. All *Lactobacillus* species isolates were revived from frozen 25% glycerol stocks on MRS agar plates (Hardy Diagnostics #89407-144), except for *Lactobacillus iners*, which was revived on Columbia Blood Agar (CBA) agar plates (Hardy Diagnostics #A16). Non-*Lactobacillus* species were revived from frozen 25% glycerol stocks on CBA agar plates. For cultivation of *Lactobacillus* species in liquid media, MRS+CQ broth was used unless otherwise noted. For cultivation of non-*Lactobacillus* species, NYCIII broth was used unless otherwise noted.

ATCC 33323 *Lactobacillus gasseri* and derivative mutant strains were cultured under anaerobic conditions for solid media and aerobic conditions for liquid media at 37°C. Strains were revived from frozen 25% glycerol stocks on MRS agar plates for 48 hours under anaerobic conditions using an anaerobic box with oxygen-eliminating sachets padded with a methylene blue anaerobic indicator (BD GasPak™ EZ Anaerobe Container System Sachets with Indicator #260001). After incubation, a single colony was picked into liquid culture and incubated aerobically at 37°C with shaking. MRS+CQ broth was used for liquid culture unless otherwise noted.

*Escherichia coli* cloning strains, EC1000 (Addgene #71852) and MC1061 (Molecular Cloning Laboratories #MC1061), *Staphylococcus aureus*, and *S. aureus* derivative mutant strains were cultured under aerobic conditions at 37°C with shaking for liquid cultures. EC1000 was revived from frozen 25% glycerol stocks on LB agar plates supplemented with 40 µg/mL kanamycin (KAN). After overnight incubation, a single colony was picked into liquid culture and incubated aerobically at 37°C with shaking. LB media supplemented with 40 µg/mL KAN was used for liquid culture unless otherwise noted. MC1061 strains with a pTRK892-derived plasmid (Addgene #71803) were revived from frozen 25% glycerol stocks on LB agar plates supplemented with 200 µg/mL erythromycin (ERY). LB media supplemented with 200 µg/mL ERY was used for liquid culture.

Media additives were added to autoclave-sterilized broth or agar plates after cooling to room temperature. Broth was re-sterilized using a 0.22 µM PES filter after all additives were added. To prepare MRS+CQ broth, BD Difco-formulated *Lactobacillus* MRS (De Man, Rogosa and Sharpe) broth (BD Biosciences #288130) was prepared as per the manufacturer’s instructions and then supplemented with L-cysteine (4 mM) and *L.* glutamine (1.1 mM) as previously described^43^. To prepare delipidated MRS+CQ broth, four parts MRS+CQ broth was combined with one part Cleanascite™ Lipid Removal Reagent (Biotech Support Group #X2555-10) and shaken (220 rpm) at room temperature for 10 min, then centrifuged at 2,000 x g for 15 min. The media supernatant was collected and re-sterilized using a 0.22 uM PES filter. NYCIII broth (ATCC medium 1685) pre-media was prepared with 15 g/L Proteose Peptone No. 3 (BD Biosciences #211693), 5 g/L sodium chloride, and 4 g/L HEPES in 875 mL of distilled water, pH-adjusted to 7.3, and autoclaved. After autoclaving and cooling the pre-media, dextrose solution (3 g / 45 mL distilled water) was added at 7.5% v/v, Gibco yeast extract solution (Gibco #18180-059) was added at 2.5% v/v, and heat inactivated horse serum (Gibco #26050070) was added at 10% v/v. The complete NYCIII broth was then sterilized by passage through a 0.22 μm PES filter. S-broth^43^ pre-media was prepared with 37 g/L Brian Heart Infusion (BHI) broth (BD Biosciences #211059), 10 g/L yeast extract powder, and 1 g/L dextrose in 880 mL distilled water, brought to a boil until dissolved, and then autoclaved. After autoclaving and cooling the pre-media, fetal bovine serum (Millipore Sigma #F4135) was added at 5% v/v, vitamin K1-Hemin solution (BD Biosciences #B12354) was added at 5% v/v, and IsoVitaleX (BD Biosciences #B11875) enrichment was added at 2% v/v. The complete S-broth was then sterilized by passage through a 0.22 µm PES filter. To prepare LB media and agar, BD Difco-formulated Lysogeny broth (LB) media (BD Biosciences #244610) or agar (BD Biosciences #240110) plates were prepared as per the manufacturer’s instructions and then supplemented with pre-sterilized stocks of antibiotics or sodium acetate where indicated. For all fatty acid supplementations, oleic acid (≥99% purity Millipore Sigma #O1008), universally ^13^C_18_-labeled oleic acid (^13^C_18_-OA; ≥99% purity, Millipore Sigma #490431), palmitoleic acid (≥98.5% purity, Millipore Sigma #P9417), and linoleic acid (≥99% purity, Millipore Sigma #L1012) were pre-sterilized by passage through a 0.22 µm filter and directly added to broth media.

### FRESH cohort and human samples

The FRESH cohort is an ongoing prospective observational study based in Umlazi, South Africa, that enrolls 18–23 year old, HIV-uninfected, sexually active, healthy women. Exclusion criteria included pregnancy, anemia, any chronic medical condition or other conflict likely to prevent study protocol adherence, and/or enrollment in any other study that involves frequent blood sampling or that might otherwise interfere with the FRESH study protocol. Study characteristics and inclusion and exclusion criteria have been described in detail elsewhere^10,13,43,90^. The study protocol was approved by the Massachusetts General Hospital (MGH) Institutional Review Board (IRB, 2012P001812/ MGH) and the Biomedical Research Ethics Committee of the University of KwaZulu-Natal (UKZN; Ethics Reference Number BF131/11). All participants provided written informed consent.

Twice per week at the study site, participants received HIV RNA viral load testing by PCR and attended classes focused on personal empowerment, job skills training, and HIV prevention. Once every 12 weeks, participants provided peripheral blood and mucosal specimens described in more detail below. Participants were additionally provided a light meal during study site visits and cumulative monetary compensation over a 36-week period of ZAR3,700 (∼US$280), meant to help defray transportation expenses for the twice-weekly study site visits.

For FRESH study mucosal specimen collections, cervicovaginal swab samples (Puritan 6” sterile standard foam swab with polystyrene handle) were collected by swabbing the ectocervix in two full revolutions under direct visualization during speculum exam. The swabs were then used to make a slide preparation for Gram stain analysis, cryopreserved in thioglycolate broth with 20% glycerol for bacterial isolation, or frozen without cryopreservatives for bacterial isolation or nucleic acid extraction and microbiota profiling. Cervicovaginal lavage (CVL) samples were collected using a flexible plastic bulb pipette to dispense 5 ml of sterile normal saline into the vagina and wash the cervix four times. Fluid was then re-aspirated into a 15 ml conical tube. CVL and swab samples were stored on ice for 1–4 hours during transport to the processing laboratory at the HIV Pathogenesis Programme, Doris Duke Medical Research Institute at the Nelson R. Mandela School of Medicine at UKZN, where swabs were stored at –80°C. CVL samples were centrifuged (700xg, 10 min at 4°C). Supernatants from the centrifuged CVL samples were then transferred to cryovials and stored at –80°C.

### Bacterial culture conditions for growth inhibition, growth enhancement, and killing assays

To assess mono-culture growth effects, bacterial isolate stocks were first revived on solid media for 2–5 days until fully formed colonies were present. Starter liquid cultures were then inoculated with a few representative colonies for each strain and grown for 48 hours. After 48 hours, the optical density at 600 nm (OD600) of each liquid mono-culture was measured and each cultivated strain was back-diluted to 40X the starting OD600 (e.g. 40X OD600=0.8 for a starting OD600=0.02) for each experiment using the base media specific to that experiment. In cases where growth enhancement was being assessed, starter cultures were pelleted (5,000xg, 5 min at RT), washed with phosphate-buffered saline (PBS) three times, resuspended in PBS, and back-diluted to 40X the starting OD600 prior to assay inoculation to prevent nutrient carryover. Back-diluted starter cultures were used to inoculate the experimental media conditions. At the selected time point for each experiment, growth was assessed by optical density at 600 nm (OD600) with a SpectraMax M5 (Molecular Devices).

To assess bactericidal activity of OA, a minimum bactericidal concentration (MBC) assay was performed in MRS+CQ broth with varying concentrations of OA. Starter cultures and experimental conditions were prepared as described above. After 24 hours of incubation, colony forming units (CFU) were measured by serially diluted each culture condition 10-fold (e.g. 10 µL of culture into 100 µL of media) and 10 µL of each dilution was plated from neat (undiluted) to 10 ^-7^ dilution. Colonies were counted manually after 48–72 hours of incubation.

### Competition and mock community culture experiments

Starter cultures of representative strains of *L. crispatus*, *L. gasseri*, *L. iners*, *L. jensenii*, *G. piotii*, *G. vaginalis*, *P. amnii*, *P. bivia*, *P. disiens*, *P. timonensis*, *F. vaginae*, and *S. amnii* were prepared in NYCIII broth. Aliquots of mono-cultured strains were mixed in defined ratios (see CFU input for each strain relative to *L. crispatus* 1 in Table S9) and then divided and used to inoculate replicate cultures across different treatment conditions for incubation (6 replicates per condition) by adding 150 µL of mixture in V-bottom 96 well plates. Treatment conditions included NYCIII broth (untreated), NYCIII broth supplemented with 3.2 mM oleic acid, NYCIII supplemented with 50 µg/mL metronidazole, and NYCIII broth supplemented with both 3.2 mM oleic acid and 50 µg/mL metronidazole. CFU (colony forming unit) titres were determined for each axenic culture as described above and used to calculate starting ratios within the mixed cultures. At 72 hours, cultures were harvested by centrifuging at 5,000xg for 10 min at 4°C, supernatant was removed, and pellets were frozen at –20°C for subsequent DNA extraction and analysis. Relative growth within mixed cultures was assessed by bacterial 16S rRNA gene sequencing as described below. OD600 was determined for each input community mixture (t=0 hours) and the final time point (t=72 hours) in the no-treatment control condition to confirm growth of the defined, in vitro community cultures. Aliquots of the mono-cultured strains were assayed separately in all treatment conditions to confirm expected growth patterns, and assessed by OD600 at the corresponding time points to confirm purity.

### Sample preparation and conditions for TEM imaging

For TEM sample preparation, representative strains of *L. crispatus* and *L. iners* were revived on solid media, representative colonies were picked into liquid cultures, and incubated for 48 hours under standard culture conditions. Cultures were back diluted to a starting OD600 of 0.02 in untreated media and grown to exponential phase, OD600=0.4–0.6. Exponential cultures were then either left untreated or treated with 3.2 mM of oleic acid for 1 hour. After fatty acid treatment, cultures were removed from the anaerobic chamber, combined in a 1:1 ratio with fixative solution prepared at 2X the working concentration, and promptly pelleted (5,000xg, 5 min at RT). Samples were kept at room temperature for 2 hours to allow for pellet fixation. Fixative solution at 2X the working concentration contained 5% Glutaraldehyde (Electron Microscopy Services (EMS) #16000), 2.5% Paraformaldehyde (EMS #19200), 0.06% picric acid (EMS #19552), 20 mM Lysine (EMS #L5626), and 0.2% Ruthenium Red (EMS #20600) in 0.2 M cacodylate buffer pH 7.4.

Fixed pellets were washed three times in 0.1% Ruthenium Red in 0.1 M cacodylate buffer (EMS #11650) and postfixed with 1% Osmium tetroxide (OsO4, EMS #19100) + 0.1% Ruthenium Red for 2 hours, washed two times in water + 0.1% Ruthenium Red, and subsequently dehydrated in grades of ethanol (10 min each; 50%, 70%, and 90% ethanol once, and 100% ethanol twice). The samples were then put in propylene oxide (EMS #20401) for 1 hour and infiltrated overnight in a 1:1 mixture of propylene oxide and Spurr’s Low Viscosity Embedding media (EMS Catalog #14300). The following day the samples were embedded in Spurr’s Low Viscosity Embedding media (EMS #14300) and polymerized at 60°C for 48 hours. Ultrathin sections (about 80 nm) were cut on a Reichert Ultracut-S microtome, picked up on to copper grids (EMS #G-GVPB-Cu) stained with lead citrate (EMS 22410) and examined in a JEOL 1200EX Transmission electron microscope and images were recorded with an AMT 2k CCD camera.

### Sample preparation for bulk RNA-sequencing of bacterial isolates

For transcriptomic profiling of *cis*-9-uLCFA-treated non-*iners Lactobacillus* cultures, representative strains of *L. crispatus*, *L. iners*, *L. gasseri*, and *L. jensenii* were revived on solid media, then representative colonies were picked into liquid cultures and incubated for 48 hours under standard culture conditions. Cultures were back-diluted to a starting OD600 of 0.02 in untreated media and grown to exponential phase, OD600=0.4–0.6. Exponential cultures were then either left untreated or treated with OA, LOA, or POA (3.2 mM each). To collect samples for bulk RNA sequencing after 1 hour of treatment, cultures were pelleted (5,000xg, 5 min at 4°C), the supernatant removed, and the pellets were resuspended in 500 µL of Trizol (Life Technologies Corporation #15596026), immediately put on dry ice, and stored at –80°C until RNA extraction and library preparation. Three replicates, each from an independent starter culture, were included for each strain and condition.

### RNA extraction from bacterial isolates

Trizol-preserved bacterial samples were thawed, transferred to 2 mL FastPrep tubes (MP Biomedicals #115065002) containing 500 µL of 0.1 mm Zirconia/Silica beads (BioSpec Products #11079101z), and bead beaten for 90 seconds at 10 m/sec speed using the FastPrep-24 5G (MP Biomedicals #116005500) with a Metal QuickPrep adapter. After bead beating, samples were incubated on ice for 3 min, and 200 µL chloroform was added to each sample and mixed by tube inversion. After 3 min incubation at room temperature, samples were centrifuged (12,000xg, 15 min at 4°C) to form separated phase layers between the organic & aqueous sections. For each sample, 200 µL of the clear aqueous phase was transferred to separate clean tubes, mixed with an equal volume (200 µL) of 100% ethanol by tube inversion, and incubated for 5 min at RT. Using a Direct-zol RNA Purification kit (Zymo Research #R2070), each sample was then transferred to a Direct-zol spin column and centrifuged (16,500xg, 1 min at RT). Columns were washed twice with 400 µL Direct-zol RNA Pre-Wash Buffer, and once with 700 µL Direct-zol RNA Wash Buffer and then centrifuged at max speed for 2 min. To dry the pellet and remove any ethanol carryover, columns were centrifuged with lids open (12,000xg, 1 min at RT). Finally, columns were moved to RNase-free 1.5mL tubes, incubated with 100 µL of nuclease-free water for 5 min, and then eluted by centrifugation (12,000xg, 1 min at RT). Extracted RNA samples were kept on ice for use and QC or stored at –80°C.

### Library preparation for bulk RNA-sequencing of bacterial isolates

Illumina cDNA libraries were generated using a modified version of the RNAtag-seq protocol^91^. Briefly, 250 ng of total RNA was fragmented, depleted of genomic DNA, dephosphorylated, and ligated to DNA adapters carrying 5’-AN_8_-3’ barcodes of known sequence with a 5’ phosphate and a 3’ blocking group (IDT). Barcoded RNA molecules were pooled and depleted of rRNA using the Pan-Bacteria riboPOOL depletion kit (siTOOLs Biotech, Galen Laboratories #dp-K096). Pools of barcoded RNAs were converted to Illumina cDNA libraries in 2 main steps: (i) reverse transcription of the RNA using a primer designed to the constant region of the barcoded adaptor with addition of an adapter to the 3’ end of the cDNA by template switching using SMARTScribe Reverse Transcriptase (Takara ClonTech #639538) as described^92^; (ii) PCR amplification using primers whose 5’ ends target the constant regions of the 3’ or 5’ adaptors and whose 3’ ends contain the full Illumina P5 or P7 sequences. cDNA libraries were sequenced on the Illumina NovaSeq SP 100 platform to generate paired end reads.

### Analysis of RNA-sequencing data

Since samples were barcoded in library preparation and then pooled for sequencing, reads from each sample were demultiplexed based on their associated barcode sequence using custom scripts. Up to 1 mismatch in the barcode was allowed provided it did not assign the read a separate barcode included in the sequencing pool. Barcode sequences were removed from the first read as were terminal G’s from the second read that may have been added by SMARTScribe during template switching.

For each sample, reads were aligned to the sample species reference genome (GCF_022455535.1 / ASM2245553v1 for *L. crispatus*; GCF_022456925.1 / ASM2245692v1 for *L. gasseri*; GCF_022456915.1 / ASM2245691v1 for *L. jensenii*) using BWA^93^ and read counts were assigned to genes and other genomic features using custom scripts. Differential expression analysis was conducted using DESeq2^46^.

### Preparation of *Lactobacillus* spent media and pellets for untargeted lipidomics and isotopic tracing

Starter cultures were prepared as described above, pelleted, and washed 3 times with PBS. Washed pellets were resuspended in PBS using the same volume as the initial start culture. Washed starter cultures were then back-diluted into 0, 0.1, or 3.2 mM OA or ^13^C_18_-OA the tested media conditions and allowed to incubate under standard culture conditions for 72 hours. After incubation, cultures were pelleted (5,000xg, 10 min at 4°C). Supernatants were removed, collected into separate tubes, and placed onto dry ice until storage at –80°C. Cell pellets were washed once with ice cold PBS. Washed pellets were also placed onto dry ice and then stored at –80°C once frozen. Three replicates were included for each strain and condition.

### Plasmid construction methods for strain generation in *L. gasseri* ATCC 33323

All vectors and primers with their respective annealing temperatures are noted in Tables S5 and S6, respectively. Primers were purchased/synthesized by IDT. All cloning host strains and mutants generated are noted in Table S7. All plasmids were verified with nanopore-based whole plasmid sequencing using Primordium Labs services. All mutant strains were verified with Illumina-based WGS using SeqCenter, LLC.

Genomic DNA extractions were performed using the DNeasy Blood & Tissue Kit (Qiagen Beverly LLC #69504) following manufacturer protocols for Gram positive bacterial pellet samples. Minipreps were performed using the QIAprep Spin Miniprep Kit (Qiagen Beverly LLC #27104) following the manufacturer’s protocol. Gel extractions were performed using the QIAquick Gel Extraction Kit (Qiagen Beverly LLC #28704) following the manufacturer’s protocol. Extracted and amplified DNA products were quantified using a NanoDrop 2000. DNA bands were visualized using E-Gel™ EX Agarose Gels (2%, Invitrogen #G401002) under UV light.

Q5® High-Fidelity DNA Polymerase (New England Bio Labs (NEB) #M0491S) was used for PCR amplification of products to be used for plasmid construction following the manufacturer’s protocol. Briefly, Q5® High-Fidelity DNA Polymerase PCR reactions were performed in 25 µL reactions such that the final concentration of reagents were as follows, 1X Q5 Reaction Buffer, 200 µM dNTPs, 0.5 µM Forward Primer, 0.5 µM Reverse Primer, 0.02 units/µL Q5 High-Fidelity DNA Polymerase, and 1–10 ng of template DNA; and, thermocycling was performed at 98°C for 30 s, followed by 30 cycles of 98°C for 10 s, annealing temperature (noted in Table S6 for each primer pair) for 30 s, and 72°C for 20 s/kb of desired amplicon, with a final 2 min extension at 72°C. KAPA HiFi HotStart ReadyMix (Roche Holding AG #07958935001) was used for all colony PCR-screening reactions following the manufacturer’s protocol. KAPA HiFi HotStart ReadyMix PCR reactions were performed in 25 µL reactions such that the final concentration of reagents were as follows, 1X Ready Mix, 0.3 µM Forward Primer, 0.3 µM Reverse Primer, and a pipette-tip sample of a single colony; and, thermocycling was performed at 95°C for 3 min, followed by 35 cycles of 98°C for 20 s, annealing temperature (noted in Table S6 for each primer pair) for 15 s, and 72°C for 20 s/kb of desired amplicon, with a final 1 min extension at 72°C.

For restriction and ligation-based cloning, restriction enzyme-based digestion was performed at 37°C for 15 min following manufacturer’s protocols. Digestion reactions were performed in 50 µL reactions such that the final concentration of reagents were as follows, 1X rCutSmart Buffer and 0.4 units/µL per enzyme. Ligation was performed at room temperature for 10 min and then heat inactivated at 65°C for 10 min. Reactions were performed in 20 µL reactions such that the final concentration of reagents were as follows, 1X T4 DNA Ligase Buffer, 37.5 ng of insert fragment, 50 ng of linearized backbone, and 1 unit/µL of T4 DNA Ligase (NEB #M0202). A no-insert with backbone only control was prepared in parallel with each ligation reaction to serve as a backbone self-ligation control. For Gibson cloning, Gibson Assembly® Master Mix (NEB #E2611) was used for all Gibson assembly reactions following the manufacturer’s protocol. Gibson assemblies were performed in 20 µL reactions such that the final concentration of reagents were as follows, 1X Gibson Assembly Master Mix with 0.02–0.5 pmols of DNA fragments comprising 3–5 parts per insert fragment and 1 part linearized backbone. For plasmid construction, *E. coli* EC1000 strain^94^ (Addgene #71852), a kanamycin resistant strains carrying a single copy of the repA gene in the glgB gene, was used as the cloning host for all pORI28-based plasmids, and *E. coli* MC1061 strain (Molecular Cloning Laboratories #MC1061) was used as the cloning host for all pTRK892-based plasmids.

### Plasmid transformations methods for strain generation in *L. gasseri* ATCC 33323

Competent cells of *E. coli* EC1000 strain were prepared using the Mix & Go! *E. coli* Transformation Kit (Zymo Research #T3001) following the manufacturer’s protocol. 0.5 mL of an overnight culture of EC1000 grown in LB media at 37°C was transferred to 50 mL of ZymoBroth (Zymo Research #M3015) and cultured for 15–16 hours at 22°C with shaking to obtain a culture of OD600=0.4. Wash and Competent Buffers were diluted to 1X working concentration using the provided Dilution Buffer and kept on ice. The grown ZymoBroth culture was then pelleted (5,000xg, 10 min at 4°C), supernatant was removed, and pellets were washed with 5 mL of 1X Wash Buffer. The washed pellets were gently resuspended in 5 mL of ice cold 1X Competent Buffer and aliquoted into 100 µL volumes on ice. Aliquots were immediately used or stored at –80°C for transformation at a later time. For transformation of competent EC1000, 5 µL of plasmid DNA was added to 100 µL of cells, gently mixed, and incubated for 10 min on ice. After incubation, 400 µL of S.O.C. medium (Invitrogen #15544034) was added to the cells, which were then incubated for 1 hour at 37°C with shaking plated onto pre-warmed LB agar with antibiotic selection, and incubated overnight at 37°C. The no-insert cloning control and no DNA plasmid control samples were transformed and plated in parallel as negative controls for each transformation.

For transformation of *E. coli* MC1061, competent MC1061 were thawed on ice, aliquoted into 100 µL volumes into chilled, clean 1.5 mL microcentrifuge tubes, gently mixed with 5 µL of plasmid DNA, and incubated for 15 min on ice. Cells were heat-shocked for 45 seconds in a 42°C water bath and then immediately placed on ice for 2 min. 0.9 mL of S.O.C. medium was added, cells were incubated for 1 hour at 37°C with shaking, plated onto pre-warmed LB agar with antibiotic selection, and incubated overnight at 37°C. The no-insert cloning control and no DNA plasmid control samples were transformed and plated in parallel as negative controls for each transformation.

Competent cells of *L. gasseri* ATCC 33323 and derivative mutants were prepared fresh for each transformation. 3.5X sucrose:MgCl2 electroporation buffer (3.5X SMEB) buffer was used as the transformation buffer and prepared such that the final concentration was 952 mM sucrose and 3.5 mM MgCl2 at pH 7.2 in DI water and then sterilized using a 0.22 uM PES filter. A 500 mL stock of 3.5X SMEB was prepared fresh each month and stored at 4°C. A single colony of *L. gasseri* wild-type or mutant was picked and cultured aerobically in 100 mL of MRS broth for 15– 16 hours at 37°C with shaking. After 15–16 hours, cells were pelleted (5,000xg, 10 min at 4°C) and resuspended on ice with 100 mL of ice cold 3.5X SMEB buffer. The resuspension was carefully mixed until the pellet was completely resuspended into a homogenous solution, re-pelleted (5,000xg, 10 min at 4°C), and then resuspended on ice again in 5 mL of ice cold 3.5X SMEB to obtain the electrocompetent cell mixture. 100 µL of the electrocompetent cells were aliquoted into pre-chilled 0.2 cm gap Gene Pulser Electroporation Cuvettes (Bio-rad Laboratories #1652086) and kept on ice. 1 ug of DNA plasmid (volume kept < 10 µL to prevent sparking) was added to the cuvette, gently mixed without creating any air bubbles, and incubated on ice for 5 min. After incubation, electroporation was performed with a Gene Pulser Xcell Electroporation System using the following conditions: 1.25 kV, 25 µF, and 200 Ω. Immediately after electroporation, the cuvette was held on ice for 5 min, then the sample was added to 1 mL of pre-warmed MRS broth and incubated for 3 hours at 37°C with shaking. After incubation, cells were gently pelleted (300xg, 5 min), plated onto MRS agar plates with selection antibiotic, and incubated for 48 hours at 37°C in an anaerobic box.

### Gene knockout and complementation in *L. gasseri* ATCC 33323

*L. gasseri* ATCC 33323 was selected as a genetically tractable, representative non-*iners Lactobacillus* strain exhibiting *cis-*9-uLCFA resistance and OA growth enhancement phenotypes similar to other strains of *L. gasseri*, *L. crispatus*, *L. jensenii*, and *L. mulieris*. To generate gene knockout mutants in *L. gasseri* ATCC 33323, we adapted a previously reported uracil phosphoribosyltransferase (*upp*)-based two-plasmid homologous recombination system^95–97^. This system exploits the 5-fluorouracil (5-FU) resistance of a *upp*-deficient parent strain for knockout construction. To briefly summarize the underlying principles, uracil phosphoribosyltransferase, central to the pyrimidine salvage pathway, catalyzes the conversion of uracil to uridine monophosphate. When provided 5-FU, uracil phosphoribosyltransferase will produce 5-fluorouridine-5’-monophosphate, which is a suicide inhibitor to thymidylate synthase, a required enzyme in DNA synthesis. Vectors used to generate the *upp*-deficient strain and other gene deletion mutants included pTRK669^95^, pORI28^94^, and pORI28-derived plasmids. pORI28 is an empty backbone that requires RepA for stable plasmid replication and propagation and that encodes an erythromycin (ERY) resistance gene as a selectable marker; it is used for chromosomal integration in Gram positive bacteria. pMZ7 is a pORI28-derived backbone inserted with *lacZ* and the *L. acidophilus upp* gene (*Laupp*), which serves as the counterselection gene when integrated chromosomally. pTRK669 is a chloramphenicol (CHLOR) resistant, temperature sensitive helper plasmid that encodes RepA. To summarize the overall approach, we first generated an *upp* gene-deleted mutant in *L. gasseri* ATCC 33323 (Δ1245-WT; generated as described below), which was resistant to 5-FU. This Δ1245-WT mutant served as the parent strain for all additional mutants and is therefore referred to as the WT strain in the figures and the Results and Discussion text. The pTRK669 helper plasmid was electro-transformed into Δ1245-WT to create Δ1245-WT/pTRK669. Then pMZ7-derived vectors containing ∼600-bp homology arms within each gene-of-interest were constructed and respectively electro-transformed into Δ1245-WT/pTRK669, where RepA expression from pTRK669 enabled them to be replicated and propagated. Δ1245-WT/pTRK669 strains containing each gene-of-interest-specific pMZ7-derived vector were cultured under ERY and CHLOR selection (7.5 µg/mL CHLOR and 5 µg/mL ERY) at 37°C, then subcultured 3–5 times under ERY selection only (2.5 µg/mL) at 42°C to cure the pTRK669 helper plasmid and select for single-crossover chromosomal integrants of the pMZ7-derived vector. After these subculturing steps, cells were plated onto MRS agar plates with ERY (5 µg/mL) and colony PCR-screened for pMZ7-derived vector chromosomal integration. Confirmed clones of Δ1245-WT containing single-crossover integrants of the pMZ7-derived vector targeting the gene-of-interest were next cultured without antibiotic selection to allow for vector resolution from the chromosome, then subcultured in 5-FU to counter-select for the desired gene deletion mutant strain via a second crossover event to generate an in-frame gene deletion knockout by double homologous recombination. All subculturing steps were performed by transferring 5% of the grown culture volume into the newly prepared broth of equal volume to make a 5% inoculum.

We generated the Δ1245-WT parent strain used for the above knockout approach as follows. To construct the pMZ4 vector for generating Δ1245-WT, we generated a pORI28 derivative with homology arms flanking the endogenous *upp* gene (LGAS_1245), each being ∼600 bp for the upstream and downstream arms, cloned into pORI28 using restriction digest and ligation cloning methods. In brief, a 594-bp upstream region and a 616-bp downstream region flanking LGAS_1245 were each amplified from genomic DNA extracted from *L. gasseri* ATCC 33323 to produce PCR amplicons, LGAS_1245-up and LGAS_1245-dwn, (primers for LGAS_1245-up: 1245-SOE-1 and 1245-SOE-2; primers for LGAS_1245-dwn: 1245-SOE-3 and 1245-SOE-4). Splicing by overlap-extension PCR was used to fuse LGAS_1245-up and LGAS_1245-dwn, producing a single 1187-bp amplicon (primers: 1245-SOE-1 and 1245-SOE-4), which was gel-purified to yield only the fused amplicon product. The fused LGAS_1245 homology arm amplicon and pORI28 were each digested separately using BamHI-HF and SacI-HF. The digested pORI28 backbone was additionally treated with shrimp Alkaline Phosphatase (rSAP) during the digest reaction and gel-purified before use in cloning. The digested pORI28 backbone and fused LGAS_1245 homology arm amplicon were ligated using T4 ligase and subsequently deactivated by heating. 10 µL of the ligation reaction was used to transform competent EC1000 cells, which were then plated onto LB agar plates with 200 µg/mL ERY. The constructed plasmid, pMZ4, was miniprepped from a single colony and confirmed by sequencing.

To use the pMZ4 plasmid to generate the Δ1245-WT strain, the helper plasmid pTRK669 was first electro-transformed into freshly prepared competent cells of *L. gasseri* ATCC 33323, then plated onto MRS agar plates with 7.5 µg/mL CHLOR and incubated anaerobically for 48 hours at 37°C. Clones containing pTRK669 (*L. gasseri* ATCC 33323/pTRK669) were verified by colony PCR screening (primers: pTRK699_F1 and pTRK699_R1). Next, pMZ4 was electro-transformed into freshly prepared competent cells of verified *L. gasseri* ATCC 33323/pTRK669, which were then plated onto MRS agar plates with 7.5 µg/mL CHLOR and 5 µg/mL ERY and incubated anaerobically for 48 hours at 37°C. ATCC 33323/pTRK669+pMZ4 clones were verified by colony PCR screening (primers: pTRK699_F1 and pTRK699_R1; pORI28F1 and pORI28R1). ATCC 33323/pTRK669+pMZ4 was then broth cultured under 7.5 µg/mL CHLOR and 5 µg/mL ERY selection at 37°C, then subcultured 3–5 times under 2.5 µg/mL ERY selection only at 42°C to cure the pTRK669 helper plasmid and select for pMZ4 single-crossover chromosomal integrants, then plated onto MRS agar plates with 5 µg/mL ERY to identify pMZ4 single-crossover chromosomal integrant clones. pMZ4-integration clones verified by colony PCR screening (using primers 1245-up and 1245-dwn) were cultured without antibiotic selection, then subcultured into MRS broth with 100 µg/mL 5-FU to counter-select for a second crossover event to produce the *upp* gene deletion mutant strain, then plated onto MRS media agar with 100 µg/mL 5-FU and incubated anaerobically for 48 hours at 37°C. Multiple subcultures were grown in parallel to increase chances of obtaining an *upp* gene deletion strain. Obtained colonies were PCR-screened (primers: 1245-up and 1245-dwn) and single-colony purified 1–2 times. Purified clones were verified by WGS to confirm deletion of *upp* by double homologous recombination. This *upp*-deleted (Δ1245-WT) strain (referred to in figures and main text as WT) was thus 5-FU resistant.

To construct the pMZ7 backbone vector for generating gene-specific knockouts, *lacZ* and the *L. acidophilus upp* gene (*Laupp*) were cloned into pORI28 using two-piece Gibson cloning. In brief, *lacZ* and *Laupp* were each amplified from pUC19 and genomic DNA extracted from *L. acidophilus* ATCC 4356, respectively, to produce PCR amplicons, *lacZ* and *Laupp* (primers for *lacZ*: pUC19lacZ-F and pUC19lacZ-R; primers for *LAupp*: upp-F and upp-R). Splicing by overlap-extension PCR was used to fuse *lacZ* (780 bp) and *LAupp* (759 bp), producing a single 1519-bp amplicon (primers: pUC19lacZ-F and upp-R), which was gel-purified. The fused *lacZ*-*LAupp* amplicon and pORI28 were each PCR-amplified separately (primers for pORI28: lacZupp_intF1 and lacZupp_intR1; primers for *lacZ*-*LAupp*: lacZupp_intF2 and lacZupp_intR2) and assembled using NEB Gibson Master Mix (NEB #E2611). 10 µL of the Gibson reaction was used to transform competent EC1000 cells, then plated onto LB agar plates with 200 µg/mL ERY. The constructed plasmid, pMZ7, was miniprepped from a single colony and confirmed by sequencing.

To generate pMZ7-derived vectors for generating gene-specific deletion mutants in *L. gasseri* Δ1245-WT, pMZ9 (containing homology arms for *ohyA9*, LGAS_1351) and pMZ10 (containing homology arms for *farE*, LGAS_1630) were constructed using three-piece Gibson cloning in EC1000 cells. For pMZ9 construction, pMZ7 was linearized by PCR amplification (primers: LGAS_1351F9 and LGAS_1351R9). A 600-bp in the upstream region of LGAS_1351 (primers: LGAS_1351F10 and LGAS_1351R10) and a 600-bp in the downstream region of LGAS1351 (primers: LGAS_1351F11 and LGAS_1351R11) were PCR-amplified. All amplicons products were combined via Gibson reaction to make pMZ9. For pMZ10 construction, pMZ7 was linearized by PCR amplification (primers: LGAS_1630F9 and LGAS_1630R9). A 628-bp in the upstream region of LGAS_1630 (primers: LGAS_1630F10 and LGAS_1630R10) and a 627-bp in the downstream region of LGAS1630 (primers: LGAS_1630F11 and LGAS_1630R11) were PCR-amplified. All amplicons products were combined via Gibson reaction to make pMZ10.

To generate gene deletion mutants in Δ1245-WT, pTRK669 was electro-transformed into freshly prepared competent cells of Δ1245-WT, then plated onto MRS agar plates with 7.5 µg/mL CHLOR and incubated anaerobically for 48 hours at 37°C. Clones containing pTRK669 (Δ1245-WT/pTRK669) were verified by colony PCR screening (primers: pTRK699_F1 and pTRK699_R1). Next, the pMZ7-derived vector containing homology arms for the gene of interest (pMZ9 or pMZ10) was electro-transformed into freshly prepared competent cells of Δ1245-WT/pTRK669, then plated onto MRS agar plates with 7.5 µg/mL CHLOR and 5 µg/mL ERY and incubated anaerobically for 48 hours at 37°C. Δ1245-WT/pTRK669+pMZ7-derived_vector clones (pMZ9 or pMZ10) were verified by colony PCR screening (primers for PTRK669: pTRK699_F1 and pTRK699_R1; primers for pMZ7-derived vector: pORI28F1 and pORI28R1). To select for pMZ7-derived_vector chromosomal integration clones, Δ1245-WT/pTRK669+pMZ7-derived_vector was subjected to the same protocol described above for generating pMZ4 single-crossover chromosomal integrant clones from ATCC 33323/pTRK669+pMZ4. Colonies were PCR-screened for pMZ7-derived_vector chromosomal integration (primers for LGAS_1351: S1351_F1 and S1351_R2; primers for LGAS_1630: S1630_F1 and S1630_R2). To isolate the desired gene deletion mutant, verified pMZ7-derived_vector-integration clones were subjected to the same protocol described above for generating the Δ1245-WT strain from the pMZ4-integration clones. Obtained colonies were PCR-screened with primers flanking the gene of interest (primers for LGAS_1351: S1351_F1 and S1351_R2; primers for LGAS_1630: S1630_F1 and S1630_R2) and single-colony purified 1–2 times. Purified clones were verified by WGS to confirm deletion of the gene.

To construct expression vectors for genetic complementation, genes were cloned into pTRK892^98^, an erythromycin resistant vector backbone containing a strong Ppgm promoter, using Gibson cloning. In brief, pTRK892 was linearized by PCR amplification such that the strong Ppgm promoter was retained and original gene insert, a mutated form of GusA, was excluded (primers: pTRK892_F1G and pTRK892_R1G). To construct pMZ12 (the *ohyA9* expression vector, also referred to a p*ohyA9*), *ohyA9* (LGAS_1351) was amplified from purified genomic DNA from *L. gasseri* ATCC 33323 (primers: 1351.FOR and 1351.REV) and combined with the linearized pTRK892 via a Gibson reaction. To construct pMZ13 (the *farE* expression vector, also referred to as p*farE*), *farE* (LGAS_1630) was amplified from purified genomic DNA from *L. gasseri* ATCC 33323 (primers: 1630G.FOR and 1630G.REV) and combined with the linearized pTRK892 via a Gibson reaction. Constructs were transformed into competent MC1061 and plated onto LB agar with 200 µg/mL ERY. Plasmids were miniprepped from a single colony and confirmed by sequencing. pMZ12 was electro-transformed into freshly prepared competent cells of *ΔohyA9* to make *ΔohyA9*/p*ohyA9*, and pMZ13 was electro-transformed into freshly prepared competent cells of *ΔfarE* to make *ΔfarE*/p*farE*. Transformed cells were plated onto MRS agar plates with 5 µg/mL ERY and incubated anaerobically for 48 hours at 37°C. Complementation was verified by colony PCR-screening (primers: Erm_F1 and Erm_R1).

### Plasmid construction and complementation in *Staphylococcus aureus* USA300 *ΔSaohyA*

All vectors and genetic mutants generated in *S. aureus* are noted in Tables 5 and 7, respectively. To construct vectors for genetic complementation in the *ohyA*-knockout *S. aureus* USA300 strain (*ΔSaohyA*), His-tagged LCRIS_00661 and LCRIS_00558 genes with the appropriate restriction sites were ordered from Invitrogen and subcloned into a previously constructed *S. aureus* expression vector^49^ (pPJ480) using restriction enzymes NcoI and XhoI in TOP10 chemically competent *E. coli* (Invitrogen #C4040) to make pLCRIS_00661 and pLCRIS_00558. Constructed plasmids were verified by nanopore-based whole plasmid sequencing using Primordium Labs services. Plasmids were next laundered through *S. aureus* strain RN4220, which can accept plasmids propagated through *E. coli* due its inactivated restriction system^99,100^, and then *S. aureus* RN4220-derived plasmids were purified and used to transform *ΔSaohyA*. Electroporation was used to transform all *S. aureus* strains. Briefly, 1.5–2 µg of DNA was incubated with the *S. aureus* strain on ice for 5 min, the mixture was then electroporated in pre-chilled 0.1 cm gap Gene Pulser Electroporation Cuvettes (Bio-rad Laboratories #1652089) with a Gene Pulser Xcell Electroporation System using the following conditions: 1.6 kV, 25 µF, and 200 Ω. Immediately after electroporation, cells were incubated for 2 hours in Brain Heart Infusion broth (BHI; BD Biosciences #DF0037) at 37°C with shaking. After incubation, cells were gently pelleted (300xg, 5 min at RT), plated onto BHI agar plates with 10 µg/mL CHLOR, and incubated overnight at 32°C.

### Enzyme product characterization

In order to measure production of OhyA metabolites, strains *ΔSaohyA*/empty vector, *ΔSaohyA*/p*SaohyA*, *ΔSaohyA*/pLCRIS_00558, and *ΔSaohyA*/pLCRIS_00661 were grown to an OD600 of 0.5 in Tryptone broth containing 1% DMSO. OA or LOA was added to a final concentration of 20 µM, and the cultures were grown for 1 hour at 37°C with shaking. The cells were separated from media by centrifugation, and the medium was extracted by adding methanol to a final concentration of 80%. Extracts were centrifuged to pellet debris, and the supernatant was analyzed by LC-MS as described below to detect the presence of the *h*FAs.

Culture supernatants containing the fatty acid substrate and *h*FA product were analyzed with a Shimadzu Prominence UFLC attached to a QTrap 4500 equipped with a Turbo V ion source (Sciex). Samples were injected onto an XSelect® HSS C18, 2.5 μm, 3.0 × 150-mm column (Waters) at 45°C with a flow rate of 0.4 ml/min. Solvent A was water, and solvent B was acetonitrile. The HPLC program was as follows: starting solvent mixture of 60% B, 0–1 min isocratic with 60% B; 1–16 min linear gradient to 100% B; 16–21 min isocratic with 100% B; 21– 23 min linear gradient to 0% B; and 23–28 min isocratic with 0% B. The Sciex QTrap 4500 was operated in the negative mode, and the ion source parameters were: ion spray voltage, −4500 V; curtain gas, 30 psi; temperature, 320°C; collision gas, medium; ion source gas 1, 20 psi.; ion source gas 2, 35 psi; and declustering potential, −35 V. The system was controlled by Analyst® software (Sciex).

The Sciex QTrap 4500 mass spectrometer was operated in the negative mode using the product scan to determine the position of the hydroxyl group by direct injection. The source parameters were: ion spray voltage, −4500 V; curtain gas, 15 psi.; temperature, 250°C; collision gas, high; ion source gas 1, 15 psi; ion source gas 2, 20 psi; declustering potential, −25 V; and collision energy, −35 V. The system was controlled by Analyst® software (Sciex).

### Targeted lipidomics for the detection of *h*FAs in cervicovaginal lavage samples

Picolylamide derivatization was used to sensitively and accurately detect *h*FA abundance in CVL supernatant samples using the unique ions generated from breakage at the hydroxyl group position^56,58^. 250 µL of each human CVL sample was added to 750 µL of methanol containing 200 nM ^13^C_18_-OA (Cambridge Isotope Laboratories, Inc. #CLM-460-PK), incubated on ice for 15 min, and centrifuged at 4,000 rpm for 10 min. Supernatant was removed to a new tube and dried in a speed-vac overnight. 500 µL of oxalyl-chloride was added to the dried samples and incubated at 65°C for 15 min. After drying the samples under N_2_ gas, 500 µL of 1% 3-picolylamine in acetonitrile was added and incubated at room temperature for 15 min. After drying the samples under N_2_ gas, the samples were resuspended in 100 µL of ethanol for analysis.

Picolylamide-*h*FA were analyzed using a Shimadzu Prominence UFLC attached to a QTrap 4500 equipped with a Turbo V ion source (Sciex). Samples (5 µL) were injected onto an XSelect HSS C18, 2.5 µm, 3.0 x 150 mm column (Waters) at 45°C with a flow rate of 0.4 mL/min. Solvent A was 0.1% formic acid in water, and solvent B was acetonitrile with 0.1% formic acid. The HPLC program was the following: starting solvent mixture of 70% A/30% B; 0 to 5 min, isocratic with 30% B; 5 to 15 min, linear gradient to 100% B; 15 to 23 min, isocratic with 100% B; 23 to 25 min, linear gradient to 30% B; 25 to 30 min, isocratic with 30% B. The QTrap 4500 was operated in the positive mode, and the ion source parameters for the picolylamide-*h*FA multiple reaction monitoring (MRM) parameters were: ion spray voltage, 5,500 V; curtain gas, 15 psi; temperature, 300°C; ion source gas 1, 15 psi; ion source gas 2, 20 psi; declustering potential, 25 V, and a collision energy, 40 V. MRM masses (Q1/Q3) were: picolylamide-*h*18:0, 391.1/109.0; picolylamide-*h*18:1, 389.1/109.0; and picolylamide-^13^C_18_-OA, 391.1/109.0. The system was controlled by the Analyst software (Sciex) and analyzed with MultiQuant™ 3.0.2 software (Sciex). The relative concentration of each *h*FA was calculated based on the known amount of ^13^C_18_-OA spiked into the sample at the beginning of the sample preparation.

### Targeted lipidomics for the detection of PG metabolites in cell pellets

Lipids were extracted from bacterial cell pellets using the Bligh and Dyer method^101^. In brief, cell pellets were homogenized with a 1:2 mixture of chloroform:methanol with bead beating. The mixture was then filtered through Whatman No. 1 filter paper (Millipore Sigma #WHA1001090), and the filtrate was allowed to separate into two layers, an alcohol and chloroform layer. The alcohol layer was removed and the remaining chloroform layer contained the lipid extract. Lipid extracts were resuspended in chloroform/methanol (1:1). PG was analyzed using a Shimadzu Prominence UFLC attached to a QTrap 4500 equipped with a Turbo V ion source (Sciex). Samples were injected onto an Acquity UPLC BEH HILIC, 1.7 um, 2.1 x 150 mm column (Waters) at 45°C with a flow rate of 0.2 ml/min. Solvent A was acetonitrile, and solvent B was 15 mM ammonium formate, pH 3. The HPLC program was the following: starting solvent mixture of 96% A/4% B; 0 to 2 min, isocratic with 4% B; 2 to 20 min, linear gradient to 80% B; 20 to 23 min, isocratic with 80% B; 23 to 25 min, linear gradient to 4% B; 25 to 30 min, isocratic with 4% B. The QTrap 4500 was operated in the Q1 negative mode. The ion source parameters for Q1 were as follows: ion spray voltage, –4,500 V; curtain gas, 25 psi; temperature, 350°C; ion source gas 1, 40 psi; ion source gas 2, 60 psi; and declustering potential, –40 V. The system was controlled by the Analyst software (Sciex). The sum of the areas under each peak in the mass spectra were calculated, using LipidView software (Sciex).

### Untargeted lipidomics and isotopic tracing methods and analysis

C18-neg: Reversed-phase C18 chromatography/negative ion mode MS detection was used to measure metabolites of intermediate polarity. Analyses of metabolites of intermediate polarity, including free fatty acids and bile acids, were conducted using an LC-MS system comprised of a Shimadzu Nexera X2 U-HPLC (Shimadzu Corp.) coupled to a Q-Exactive orbitrap mass spectrometer (Thermo Fisher Scientific). Media samples (30 µL) were extracted for analyses using 90 µL of methanol containing 50ng/mL 15-methyl PGE1, 15-methyl PGA2, 15-methyl PGE2 (Cayman Chemical Co.) as internal standards. Cell pellet samples (30 µL) were extracted for analyses using 90 µL of methanol containing 50ng/mL 15-methyl PGE1, 15-methyl PGA2, 15-methyl PGE2 (Cayman Chemical Co.) as internal standards. The cell pellets were homogenized using the QIAGEN TissueLyser II with 3mm Tungsten beads for 4 min at a frequency of 20 Hz. Samples were centrifuged (10 min, 15000xg at 4°C). After centrifugation, supernatants (2 µL) were injected directly onto a 150 x 2.1 mm, 1.8 µm ACQUITY HSS T3 C18 column (Waters). The column was eluted isocratically with 80% mobile phase A (0.01% formic acid in water) for 3 min followed by a linear gradient to 100% mobile phase B (0.01% acetic acid in acetonitrile) over 12 min. MS analyses were carried out using electrospray ionization in the positive ion mode using full scan analysis over 70–850 m/z at 70,000 resolution and 3 Hz data acquisition rate. Other MS settings were: sheath gas 45, in source CID 5 eV, sweep gas 10, spray voltage –3.5 kV, capillary temperature 320°C, S-lens RF 60, probe heater temperature 300°C, microscans 1, automatic gain control target 1e6, and maximum ion time 250 ms. Raw data were processed using TraceFinder software (Thermo Fisher Scientific) for targeted peak integration and manual review of a subset of identified metabolites and using Progenesis QI (Nonlinear Dynamics) for peak detection and integration of both metabolites of known identity and unknowns.

C8-pos: Reversed-phase C8 chromatography/positive ion mode MS detection was used to measure lipids. Analyses of polar and non-polar plasma lipids were conducted using an LC-MS system comprising a Shimadzu Nexera X2 U-HPLC (Shimadzu Corp.) coupled to an Exactive Plus orbitrap mass spectrometer (Thermo Fisher Scientific). Media samples (10 µL) were extracted for lipid analyses using 190 µL of isopropanol containing 1,2-didodecanoyl-sn-glycero-3-phosphocholine (Avanti Polar Lipids) as an internal standard. Cell pellets (∼30ul) were extracted for lipid analysis using 570ul isopropanol containing 1,2-didodecanoyl-sn-glycero-3-phosphocholine (Avanti Polar Lipids) as an internal standard. The cell pellets were homogenized using the QIAGEN TissueLyser II with 3mm Tungsten beads for 4 min at a frequency of 20Hz. The media and cell pellets were centrifuged (10 min, 9,000xg at 4°C), and were injected directly onto a 100 x 2.1 mm, 1.7 µm ACQUITY BEH C8 column (Waters). The column was eluted isocratically with 80% mobile phase A (95:5:0.1 vol/vol/vol 10mM ammonium acetate/methanol/formic acid) for 1 minute followed by a linear gradient to 80% mobile-phase B (99.9:0.1 vol/vol methanol/formic acid) over 2 min, a linear gradient to 100% mobile phase B over 7 min, then 3 min at 100% mobile-phase B. MS analyses were carried out using electrospray ionization in the positive ion mode using full scan analysis over 200–1100 m/z at 70,000 resolution and 3 Hz data acquisition rate. Other MS settings were: sheath gas 50, in source CID 5 eV, sweep gas 5, spray voltage 3 kV, capillary temperature 300°C, S-lens RF 60, heater temperature 300°C, microscans 1, automatic gain control target 1e6, and maximum ion time 100 ms. Raw data were processed using TraceFinder software (Thermo Fisher Scientific) for targeted peak integration and manual review of a subset of identified lipids and using Progenesis QI (Nonlinear Dynamics) for peak detection and integration of both lipids of known identity and unknowns. Lipid identities were determined based on comparison to reference plasma extracts and are denoted by total number of carbons in the lipid acyl chain(s) and total number of double bonds in the lipid acyl chain(s).

HILIC-neg: Targeted negative ion mode MS detection was used to measure central metabolites. HILIC (hydrophilic interaction chromatography) analyses of water soluble metabolites in the negative ionization mode (HILIC-neg) were conducted using an LC-MS system comprised of an AQUITY UPLC system (Waters) and a 5500 QTRAP mass spectrometer (SCIEX). Media samples (30 µL) were extracted with the addition of four volumes of 80% methanol containing inosine - 15N4, thymine-d4 and glycocholate-d4 internal standards (Cambridge Isotope Laboratories). Cell pellets (∼30ul) were extracted with the addition of four volumes of 80% methanol containing inosine-15N4, thymine-d4 and glycocholate-d4 internal standards (Cambridge Isotope Laboratories. The cell pellets were homogenized using the QIAGEN TissueLyser II with 3mm Tungsten beads for 4 min at a frequency of 20 Hz. The samples were centrifuged (10 min, 9,000xg at 4°C), and the supernatants were injected directly onto a 150 x 2.0 mm Luna NH2 column (Phenomenex). The column was eluted at a flow rate of 400 µL/min with initial conditions of 10% mobile phase A (20 mM ammonium acetate and 20 mM ammonium hydroxide in water) and 90% mobile phase B (10 mM ammonium hydroxide in 75:25 v/v acetonitrile/methanol) followed by a 10 min linear gradient to 100% mobile phase A. MS analyses were carried out using electrospray ionization and selective multiple reaction monitoring scans in the negative ion mode. To create the method, declustering potentials and collision energies were optimized for each metabolite by infusion of reference standards. The ion spray voltage was –4.5 kV and the source temperature was 500°C. Raw data from the 5500 QTRAP MS system were processed using MultiQuant 2.1 software (SCIEX).

### Nucleic acid extraction for 16s rRNA gene sequencing

For cervicovaginal swab samples, total nucleic acids were extracted from the swab samples using the phenol-chloroform method, which includes a bead beating process to disrupt bacteria, as previously described^13,102^. Briefly, swabs were thawed on ice, transferred into a solution consisting of phenol:chloroform:isoamyl alcohol (PCI, 25:24:1, pH 7.9, Ambion) and 20% sodium dodecyl sulfate in Tris-EDTA buffer with sterile 0.1 mm glass beads (BioSpec Products #11079101), vigorously rubbed against the walls of the tube to dislodge microbial material, and then incubated on ice for 5–10 min. Swabs were then removed by pressing the swab against the side of the tube using a sterile pipette tip as being lifted out to squeeze out excess fluid. Samples were homogenized using a bead beater for 2 min at 4°C, and then centrifuged at 6,800×g for 3 min at 4°C. The aqueous phase was transferred to a clean tube with equal volume of PCI solution, vortexed, and centrifuged again at 16,000×g for 5 min at 4°C. The aqueous phase was transferred to a second clean tube, precipitated using 0.8 volume of −20°C isopropanol with 0.08 volume (relative to initial sample) 3 M sodium acetate at pH 5.5, inverted to mix, and incubated overnight at −20°C. Samples were then centrifuged for 30 min at 21,100×g at 4°C, washed in 0.5 ml 100% ethanol and centrifuged for 15 min at 21,100×g at 4°C. The ethanol supernatant was discarded while keeping the pellet, which was allowed to air-dry and then resuspended in 20μl molecular-grade Tris-EDTA buffer. Genomic DNA from mock communities cultured in vitro was extracted using a plate-based adaptation of the above protocol including a bead beating process combined with phenol–chloroform isolation with QIAamp 96 DNA QIAcube HT kit (Qiagen Beverly LLC #51331) protocols^43^.

### 16s rRNA gene sequencing for vaginal swab and mock community samples

Bacterial microbiota composition from cervicovaginal swabs collected from FRESH study participants and compositions of defined bacterial mock community experiments were determined using Illumina-based amplicon sequencing of the V4 region of the bacterial 16S rRNA gene. Standard PCR-amplification protocols were used to amplify the V4 region of the bacterial 16S rRNA gene^13,102,103^. Briefly, samples were amplified using 0.5 units of Q5 high-fidelity DNA polymerase (NEB #M0491S) in 25 μl reaction with 1X Q5 reaction buffer, 0.2 mM deoxyribonucleotide triphosphate mix, 200 pM 515F primer (5’-AATGATACGGCGACCACCGAGACGTACGTACGGT<u>GTGCCAGCMGCCGCGGTAA</u>-3’, the underlined sequence representing the complementary region to the bacterial 16S rRNA gene; IDT) and 200 pM barcoded 806R primer (5’-CAAGCAGAAGACGGCATACGAGATXXXXXXXXXXXXAGTCAGTCAGC<u>CGGACTACHVGGG TWTCTAAT</u>-3’, the underlined sequence representing the complementary region to the bacterial 16S rRNA gene and the X characters representing the barcode position; IDT) in PCR-clean water. For each prepared barcode master mix, a water-template negative control reaction was performed in parallel. Blank extraction and amplification controls were additionally performed using unique barcoded primers in sequencing libraries. For FGT microbiota profiling from cervicovaginal swab samples, DNA was amplified in triplicate reactions and then triplicates were combined before library pooling to minimize stochastic amplification biases. For defined bacterial mock community experiments, DNA was amplified in a single reaction per replicate culture. Amplification was performed at 98°C for 30 s, followed by 30 cycles of 98°C for 10 s, 60°C for 30 s, and 72°C for 20 s, with a final 2 min extension at 72°C.

PCR products from all samples and the matched water-template control were visualized via agarose gel electrophoresis to confirm successful target amplification and absence of background amplification. Gel band strength was used to semi-quantitatively estimate relative amplicon concentrations for library pooling. To prepare the sequencing libraries, 3–20 μl of individual PCR products (adjusted on the basis of estimated relative amplicon concentration) were combined into 100 μl subpools and purified using an UltraClean 96 PCR cleanup kit (Qiagen Beverly LLC #12596-4). Despite not producing visible PCR bands, blank extractions, water-template and (for in vitro experiments) blank media controls were included in the sequencing libraries for additional quality control verification. Concentrations of the subpools were quantified using a Nanodrop 2000 and then pooled at equal molar concentrations to assemble the final library. Following standard Illumina protocols, the pooled library was diluted and supplemented with 10% PhiX, and then single-end sequenced on an Illumina MiSeq using a v2 300-cycle sequencing kit with addition of custom Earth Microbiome Project sequencing primers (read 1 sequencing primer: 5’-ACGTACGTACGGTGTGCCAGCMGCCGCGGTAA-3’; read 2 sequencing primer: 5’-ACGTACGTACCCGGACTACHVGGGTWTCTAAT-3’; index sequencing primer: 5’-ATTAGAWACCCBDGTAGTCCGGCTGACTGACT-3’; IDT)^103^.

### Analysis of 16S rRNA gene sequencing results

QIIME I v1.9.188^104^ was used to demultiplex Illumina MiSeq bacterial 16S rRNA gene sequence data. QIIME 1-formatted mapping files were used and validated using validate_mapping_file.py, sequences were demultiplexed using split_libraries_fastq.py with parameter store_demultiplexed_fastq with no quality filtering or trimming, and demultiplexed sequences were organized into individual fastq files using split_sequence_file_on_sample_ids.py^102^. Dada2 v1.6.089^105^ in R was used to filter and trim reads at positions 10 (left) and 230 (right) using the filterAndTrim function with parameters truncQ=11, MaxEE=2 and MaxN=0. Then, sequences were inferred and initial taxonomy assigned using the dada2 assignTaxonomy function, employing the Ribosomal Database Project training database rdp_train_set_16.fa.gz (https://www.mothur.org/wiki/RDP_reference_files). Taxonomic assignments were refined and extended via manual review (see Table S8 for amplicon sequence variant (ASV) taxonomy). Phlyoseq v1.30.090^106^ in R was used to analyze and process the denoised dada2 results with final taxonomic assignment and custom R scripts. Final analysis and visualization of results were performed in python using jupyter notebooks. Initial sequencing-based analysis of FGT microbiota composition from FRESH cohort swab samples, taxonomy assignment and cervicotype assignments was performed blinded to participants’ corresponding Nugent scores.

For 16S rRNA gene-based microbiome profiling of clinical samples, microbial communities were classified into 4 cervicotypes (CTs) as previously defined^13^ in a non-overlapping subset of participants from the FRESH cohort: CT1 includes communities with >50% relative abundance of non-*iners Lactobacillus* species (which consists almost entirely of *L. crispatus* in this population); CT2 consists of communities in which *L. iners* is the most dominant taxon; CT3 consists of communities in which the genus *Gardnerella* is the most dominant taxon; and CT4 consists of communities dominated by other species, typically featuring high abundance of one or more *Prevotella* species. ASVs that could not be defined to the level of taxonomic class were pruned from the dataset. Taxonomically defined ASVs were collapsed at the species or genus level as indicated for further visualization and statistical analyses.

For 16S rRNA gene sequence analysis for bacterial mock community experiments, sequences were generated, processed, and annotated as described above. For each community replicate, relative abundances of each experimental strain were determined and the ratios of non-*iners* FGT *Lactobacillus* species read counts to the sum of the read counts of all of the other experimental strains were determined to quantify non-*iners* FGT *Lactobacillus* species enrichment for each condition. Significance of between-group differences for each mixture was determined by one-way ANOVA, and significance of pairwise comparisons was calculated using Tukey’s test, with statistical values for all pairwise conditions reported in Table S4.

### Genomic analyses of FGT *Lactobacillus* and other species

All genomes of experimentally tested strains were obtained from RefSeq or Genbank (see Table S5 for accession numbers). We utilized the extensive genome collection of FGT *Lactobacillus* species reported by Bloom et al. (including isolate genomes and metagenome-assembled genomes)^43^ and the type strain genome sequences of species in the family Lactobacillaceae reported in Zheng et al.^55^ to identify presence/absence profiles of genes of interest across FGT *Lactobacillus* and Lactobacillaceae species. All genomes were downloaded from NCBI RefSeq or GenBank. Gene prediction for all genomes was performed using Prodigal, and gene functions were predicted using eggNOG 5.0^85^ employing eggNOG-mapper v2.1.9^86^. Custom Python scripts were used to parse the eggNOG outputs to identify the presence of genes of interest in each genome. We constructed a gene presence and absence map for all genes of interest across all genomes. For each gene of interest, MUSCLE v5.1^87^ was used for multiple sequence alignment of representative orthologs, ModelTest-NG^107^ was used to select the optimal substitution model, and RAxML-NG^88^ used for tree construction employed via raxmlGUI 2.0^108^ to map their phylogenetic relationships. For the construction of species phylogeny, core ribosomal genes present in all Lactobacillaceae genomes were aligned using MUSCLE v5.1^87^ and FastTree v2.1^109^ was used to construct the Lactobacillaceae species tree. Trees were constructed using ortholog sequences from each species that were representative of the majority of sequences found in their respective orthologous groups to ensure robustness of the phylogenetic reconstruction. Tree and corresponding metadata visualization was done using Interactive Tree Of Life (iTOL) v5^110^.

### Quantification and statistical analyses

For bacterial growth assays, figures depict representative results from 1 of ≥2 independent experiments prepared with distinct batches of media and bacterial input inocula. Growth data collection and analysis were not performed blind to the conditions of the experiments. For bulk RNA-sequencing samples, all conditions were performed in triplicate repeats with each inocula coming from a unique bacterial colony. For mass spectrometry experiments, all conditions were performed in duplicate or triplicate technical repeats. For mock community assays, all conditions were performed with six technical repeats.

Data analysis, statistics and visualization were performed using custom scripts written in python v3.9 using Jupyter Notebook v6.5.2 or R v.3.6.3. R packages used for analyses and plotting include seqinr v.4.2.5, tidyverse, v.1.3.1, knitr v.1.33, ggpubr v.0.4.0, DescTools v.0.99.41, gtools v.3.8.2, gridExtra v.2.3, cowplot v.1.1.1, scales v.1.1.1, grid v.3.6.3, broom v.0.7.6, e1071 v.1.7.6, and table1 v.1.4. Python packages used for analyses and plotting include biopython v1.79, matplotlib v3.7.1, numpy v1.22.3, pandas v1.5.1, scikit-bio v0.5.8, scipy v1.9.3, seaborn v0.11.2, statannot v0.2.3, and statsmodels v0.13.2. All P values are two-sided with statistical significance defined at α=0.05, unless otherwise indicated.

## References

1. Fettweis, J.M., Serrano, M.G., Brooks, J.P., Edwards, D.J., Girerd, P.H., Parikh, H.I., Huang, B., Arodz, T.J., Edupuganti, L., Glascock, A.L., et al. (2019). The vaginal microbiome and preterm birth. Nat. Med. 25, 1012–1021.

2. Moreno, I., Codoñer, F.M., Vilella, F., Valbuena, D., Martinez-Blanch, J.F., Jimenez-Almazán, J., Alonso, R., Alamá, P., Remohí, J., Pellicer, A., et al. (2016). Evidence that the endometrial microbiota has an effect on implantation success or failure. Am. J. Obstet. Gynecol. 215, 684–703.

3. Ravel, J., Moreno, I., and Simón, C. (2021). Bacterial vaginosis and its association with infertility, endometritis, and pelvic inflammatory disease. Am. J. Obstet. Gynecol. 224, 251–257.

4. Moore, D.E., Soules, M.R., Klein, N.A., Fujimoto, V.Y., Agnew, K.J., and Eschenbach, D.A. (2000). Bacteria in the transfer catheter tip influence the live-birth rate after in vitro fertilization. Fertil. Steril. 74, 1118–1124.

5. Gaudoin, M., Rekha, P., Morris, A., Lynch, J., and Acharya, U. (1999). Bacterial vaginosis and past chlamydial infection are strongly and independently associated with tubal infertility but do not affect in vitro fertilization success rates. Fertil. Steril. 72, 730–732.

6. Norenhag, J., Du, J., Olovsson, M., Verstraelen, H., Engstrand, L., and Brusselaers, N. (2020). The vaginal microbiota, human papillomavirus and cervical dysplasia: a systematic review and network meta-analysis. BJOG 127, 171–180.

7. Mitra, A., MacIntyre, D.A., Ntritsos, G., Smith, A., Tsilidis, K.K., Marchesi, J.R., Bennett, P.R., Moscicki, A.-B., and Kyrgiou, M. (2020). The vaginal microbiota associates with the regression of untreated cervical intraepithelial neoplasia 2 lesions. Nat. Commun. 11, 1999.

8. Usyk, M., Zolnik, C.P., Castle, P.E., Porras, C., Herrero, R., Gradissimo, A., Gonzalez, P., Safaeian, M., Schiffman, M., Burk, R.D., et al. (2020). Cervicovaginal microbiome and natural history of HPV in a longitudinal study. PLoS Pathog. 16, e1008376.

9. Allsworth, J.E., and Peipert, J.F. (2011). Severity of bacterial vaginosis and the risk of sexually transmitted infection. Am. J. Obstet. Gynecol. 205, 113.e1–e6.

10. Gosmann, C., Anahtar, M.N., Handley, S.A., Farcasanu, M., Abu-Ali, G., Bowman, B.A., Padavattan, N., Desai, C., Droit, L., Moodley, A., et al. (2017). Lactobacillus-Deficient Cervicovaginal Bacterial Communities Are Associated with Increased HIV Acquisition in Young South African Women. Immunity 46, 29–37.

11. McClelland, R.S., Lingappa, J.R., Srinivasan, S., Kinuthia, J., John-Stewart, G.C., Jaoko, W., Richardson, B.A., Yuhas, K., Fiedler, T.L., Mandaliya, K.N., et al. (2018). Evaluation of the association between the concentrations of key vaginal bacteria and the increased risk of HIV acquisition in African women from five cohorts: a nested case-control study. Lancet Infect. Dis. 18, 554–564.

12. Kenyon, C., Colebunders, R., and Crucitti, T. (2013). The global epidemiology of bacterial vaginosis: a systematic review. Am. J. Obstet. Gynecol. 209, 505–523.

13. Anahtar, M.N., Byrne, E.H., Doherty, K.E., Bowman, B.A., Yamamoto, H.S., Soumillon, M., Padavattan, N., Ismail, N., Moodley, A., Sabatini, M.E., et al. (2015). Cervicovaginal bacteria are a major modulator of host inflammatory responses in the female genital tract. Immunity 42, 965–976.

14. Lennard, K., Dabee, S., Barnabas, S.L., Havyarimana, E., Blakney, A., Jaumdally, S.Z., Botha, G., Mkhize, N.N., Bekker, L.-G., Lewis, D.A., et al. (2018). Microbial Composition Predicts Genital Tract Inflammation and Persistent Bacterial Vaginosis in South African Adolescent Females. Infect. Immun. 86. 10.1128/IAI.00410-17.

15. Jespers, V., Kyongo, J., Joseph, S., Hardy, L., Cools, P., Crucitti, T., Mwaura, M., Ndayisaba, G., Delany-Moretlwe, S., Buyze, J., et al. (2017). A longitudinal analysis of the vaginal microbiota and vaginal immune mediators in women from sub-Saharan Africa. Sci. Rep. 7, 11974.

16. McKinnon, L.R., Achilles, S.L., Bradshaw, C.S., Burgener, A., Crucitti, T., Fredricks, D.N., Jaspan, H.B., Kaul, R., Kaushic, C., Klatt, N., et al. (2019). The Evolving Facets of Bacterial Vaginosis: Implications for HIV Transmission. AIDS Res. Hum. Retroviruses 35, 219–228.

17. Anahtar, M.N., Gootenberg, D.B., Mitchell, C.M., and Kwon, D.S. (2018). Cervicovaginal Microbiota and Reproductive Health: The Virtue of Simplicity. Cell Host Microbe 23, 159– 168.

18. Kindinger, L.M., Bennett, P.R., Lee, Y.S., Marchesi, J.R., Smith, A., Cacciatore, S., Holmes, E., Nicholson, J.K., Teoh, T.G., and MacIntyre, D.A. (2017). The interaction between vaginal microbiota, cervical length, and vaginal progesterone treatment for preterm birth risk. Microbiome 5, 6.

19. Munoz, A., Hayward, M.R., Bloom, S.M., Rocafort, M., Ngcapu, S., Mafunda, N.A., Xu, J., Xulu, N., Dong, M., Dong, K.L., et al. (2021). Modeling the temporal dynamics of cervicovaginal microbiota identifies targets that may promote reproductive health. Microbiome 9, 163.

20. Lambert, J.A., John, S., Sobel, J.D., and Akins, R.A. (2013). Longitudinal analysis of vaginal microbiome dynamics in women with recurrent bacterial vaginosis: recognition of the conversion process. PLoS One 8, e82599.

21. Srinivasan, S., Liu, C., Mitchell, C.M., Fiedler, T.L., Thomas, K.K., Agnew, K.J., Marrazzo, J.M., and Fredricks, D.N. (2010). Temporal variability of human vaginal bacteria and relationship with bacterial vaginosis. PLoS One 5, e10197.

22. Bradshaw, C.S., Morton, A.N., Hocking, J., Garland, S.M., Morris, M.B., Moss, L.M., Horvath, L.B., Kuzevska, I., and Fairley, C.K. (2006). High recurrence rates of bacterial vaginosis over the course of 12 months after oral metronidazole therapy and factors associated with recurrence. J. Infect. Dis. 193, 1478–1486.

23. Schwebke, J.R., Lensing, S.Y., Lee, J., Muzny, C.A., Pontius, A., Woznicki, N., Aguin, T., and Sobel, J.D. (2021). Treatment of Male Sexual Partners of Women With Bacterial Vaginosis: A Randomized, Double-Blind, Placebo-Controlled Trial. Clin. Infect. Dis. 73, e672–e679.

24. Cohen, C.R., Wierzbicki, M.R., French, A.L., Morris, S., Newmann, S., Reno, H., Green, L., Miller, S., Powell, J., Parks, T., et al. (2020). Randomized Trial of Lactin-V to Prevent Recurrence of Bacterial Vaginosis. N. Engl. J. Med. 382, 1906–1915.

25. Joag, V., Obila, O., Gajer, P., Scott, M.C., Dizzell, S., Humphrys, M., Shahabi, K., Huibner, S., Shannon, B., Tharao, W., et al. (2019). Impact of Standard Bacterial Vaginosis Treatment on the Genital Microbiota, Immune Milieu, and Ex Vivo Human Immunodeficiency Virus Susceptibility. Clin. Infect. Dis. 68, 1675–1683.

26. Mitchell, C., Manhart, L.E., Thomas, K., Fiedler, T., Fredricks, D.N., and Marrazzo, J. (2012). Behavioral predictors of colonization with Lactobacillus crispatus or Lactobacillus jensenii after treatment for bacterial vaginosis: a cohort study. Infect. Dis. Obstet. Gynecol. 2012, 706540.

27. Ravel, J., Brotman, R.M., Gajer, P., Ma, B., Nandy, M., Fadrosh, D.W., Sakamoto, J., Koenig, S.S., Fu, L., Zhou, X., et al. (2013). Daily temporal dynamics of vaginal microbiota before, during and after episodes of bacterial vaginosis. Microbiome 1, 29.

28. Verwijs, M.C., Agaba, S.K., Darby, A.C., and van de Wijgert, J.H.H.M. (2020). Impact of oral metronidazole treatment on the vaginal microbiota and correlates of treatment failure. Am. J. Obstet. Gynecol. 222, 157.e1–e157.e13.

29. Beigi, R.H., Austin, M.N., Meyn, L.A., Krohn, M.A., and Hillier, S.L. (2004). Antimicrobial resistance associated with the treatment of bacterial vaginosis. Am. J. Obstet. Gynecol. 191, 1124–1129.

30. Marrazzo, J.M., Thomas, K.K., Fiedler, T.L., Ringwood, K., and Fredricks, D.N. (2008). Relationship of specific vaginal bacteria and bacterial vaginosis treatment failure in women who have sex with women. Ann. Intern. Med. 149, 20–28.

31. Ferris, M.J., Masztal, A., Aldridge, K.E., Fortenberry, J.D., Fidel, P.L., Jr, and Martin, D.H. (2004). Association of Atopobium vaginae, a recently described metronidazole resistant anaerobe, with bacterial vaginosis. BMC Infect. Dis. 4, 5.

32. Lev-Sagie, A., Goldman-Wohl, D., Cohen, Y., Dori-Bachash, M., Leshem, A., Mor, U., Strahilevitz, J., Moses, A.E., Shapiro, H., Yagel, S., et al. (2019). Vaginal microbiome transplantation in women with intractable bacterial vaginosis. Nat. Med. 25, 1500–1504.

33. Yockey, L.J., Hussain, F.A., Bergerat, A., Reissis, A., Worrall, D., Xu, J., Gomez, I., Bloom, S.M., Mafunda, N.A., Kelly, J., et al. (2022). Screening and characterization of vaginal fluid donations for vaginal microbiota transplantation. Sci. Rep. 12, 17948.

34. Jean, S., Huang, B., Parikh, H.I., Edwards, D.J., Brooks, J.P., Kumar, N.G., Sheth, N.U., Koparde, V., Smirnova, E., Huzurbazar, S., et al. (2019). Multi-omic Microbiome Profiles in the Female Reproductive Tract in Early Pregnancy. Infectious Microbes & Diseases 1, 49.

35. Khoury, S., Gudziol, V., Grégoire, S., Cabaret, S., Menzel, S., Martine, L., Mézière, E., Soubeyre, V., Thomas-Danguin, T., Grosmaitre, X., et al. (2021). Lipidomic profile of human nasal mucosa and associations with circulating fatty acids and olfactory deficiency. Sci. Rep. 11, 16771.

36. Ma, C., Vasu, R., and Zhang, H. (2019). The Role of Long-Chain Fatty Acids in Inflammatory Bowel Disease. Mediators Inflamm. 2019, 8495913.

37. Slomiany, B.L., Murty, V.L., Mandel, I.D., Zalesna, G., and Slomiany, A. (1989). Physico-chemical characteristics of mucus glycoproteins and lipids of the human oral mucosal mucus coat in relation to caries susceptibility. Arch. Oral Biol. 34, 229–237.

38. Kengmo Tchoupa, A., Eijkelkamp, B.A., and Peschel, A. (2022). Bacterial adaptation strategies to host-derived fatty acids. Trends Microbiol. 30, 241–253.

39. Parsons, J.B., Yao, J., Frank, M.W., Jackson, P., and Rock, C.O. (2012). Membrane disruption by antimicrobial fatty acids releases low-molecular-weight proteins from Staphylococcus aureus. J. Bacteriol. 194, 5294–5304.

40. Drake, D.R., Brogden, K.A., Dawson, D.V., and Wertz, P.W. (2008). Thematic review series: skin lipids. Antimicrobial lipids at the skin surface. J. Lipid Res. 49, 4–11.

41. France, M.T., Mendes-Soares, H., and Forney, L.J. (2016). Genomic Comparisons of Lactobacillus crispatus and Lactobacillus iners Reveal Potential Ecological Drivers of Community Composition in the Vagina. Appl. Environ. Microbiol. 82, 7063–7073.

42. Macklaim, J.M., Fernandes, A.D., Di Bella, J.M., Hammond, J.-A., Reid, G., and Gloor, G.B. (2013). Comparative meta-RNA-seq of the vaginal microbiota and differential expression by Lactobacillus iners in health and dysbiosis. Microbiome 1, 12.

43. Bloom, S.M., Mafunda, N.A., Woolston, B.M., Hayward, M.R., Frempong, J.F., Abai, A.B., Xu, J., Mitchell, A.J., Westergaard, X., Hussain, F.A., et al. (2022). Cysteine dependence of Lactobacillus iners is a potential therapeutic target for vaginal microbiota modulation. Nat Microbiol 7, 434–450.

44. Vedder, E.B. (1915). Starch Agar, a Useful Culture Medium. J. Infect. Dis. 16, 385–388.

45. Arthur L. Barry, William A. Craig, Harriette Nadler, L. Barth Reller, Christine C. Sanders, Jana M. Swenson (1999). M26-A: Methods for Determining Bactericidal Activity of Antimicrobial Agents; Approved Guideline (Clinical and Laboratory Standards Institute).

46. Love, M.I., Huber, W., and Anders, S. (2014). Moderated estimation of fold change and dispersion for RNA-seq data with DESeq2. Genome Biol. 15, 550.

47. France, M.T., Rutt, L., Narina, S., Arbaugh, S., McComb, E., Humphrys, M.S., Ma, B., Hayward, M.R., Costello, E.K., Relman, D.A., et al. (2020). Complete Genome Sequences of Six Lactobacillus iners Strains Isolated from the Human Vagina. Microbiol Resour Announc 9. 10.1128/MRA.00234-20.

48. Yao, J., and Rock, C.O. (2015). How Bacterial Pathogens Eat Host Lipids: Implications for the Development of Fatty Acid Synthesis Therapeutics*. J. Biol. Chem. 290, 5940–5946.

49. Subramanian, C., Frank, M.W., Batte, J.L., Whaley, S.G., and Rock, C.O. (2019). Oleate hydratase from Staphylococcus aureus protects against palmitoleic acid, the major antimicrobial fatty acid produced by mammalian skin. J. Biol. Chem. 294, 9285–9294.

50. Radka, C.D., Batte, J.L., Frank, M.W., Young, B.M., and Rock, C.O. (2021). Structure and mechanism of Staphylococcus aureus oleate hydratase (OhyA). J. Biol. Chem. 296, 100252.

51. O’Connell, K.J., Motherway, M.O., Hennessey, A.A., Brodhun, F., Ross, R.P., Feussner, I., Stanton, C., Fitzgerald, G.F., and van Sinderen, D. (2013). Identification and characterization of an oleate hydratase-encoding gene from Bifidobacterium breve. Bioengineered 4, 313–321.

52. Volkov, A., Liavonchanka, A., Kamneva, O., Fiedler, T., Goebel, C., Kreikemeyer, B., and Feussner, I. (2010). Myosin cross-reactive antigen of Streptococcus pyogenes M49 encodes a fatty acid double bond hydratase that plays a role in oleic acid detoxification and bacterial virulence. J. Biol. Chem. 285, 10353–10361.

53. Kim, K.-R., Oh, H.-J., Park, C.-S., Hong, S.-H., Park, J.-Y., and Oh, D.-K. (2015). Unveiling of novel regio-selective fatty acid double bond hydratases from Lactobacillus acidophilus involved in the selective oxyfunctionalization of mono- and di-hydroxy fatty acids. Biotechnol. Bioeng. 112, 2206–2213.

54. Hirata, A., Kishino, S., Park, S.-B., Takeuchi, M., Kitamura, N., and Ogawa, J. (2015). A novel unsaturated fatty acid hydratase toward C16 to C22 fatty acids from Lactobacillus acidophilus. J. Lipid Res. 56, 1340–1350.

55. Zheng, J., Wittouck, S., Salvetti, E., Franz, C.M.A.P., Harris, H.M.B., Mattarelli, P., O’Toole, P.W., Pot, B., Vandamme, P., Walter, J., et al. (2020). A taxonomic note on the genus Lactobacillus: Description of 23 novel genera, emended description of the genus Lactobacillus Beijerinck 1901, and union of Lactobacillaceae and Leuconostocaceae. Int. J. Syst. Evol. Microbiol. 70, 2782–2858.

56. Radka, C.D., Batte, J.L., Frank, M.W., Rosch, J.W., and Rock, C.O. (2021). Oleate Hydratase (OhyA) Is a Virulence Determinant in Staphylococcus aureus. Microbiol Spectr 9, e0154621.

57. Ndung’u, T., Dong, K.L., Kwon, D.S., and Walker, B.D. (2018). A FRESH approach: Combining basic science and social good. Sci Immunol 3. 10.1126/sciimmunol.aau2798.

58. Li, X., and Franke, A.A. (2011). Improved LC-MS method for the determination of fatty acids in red blood cells by LC-orbitrap MS. Anal. Chem. 83, 3192–3198.

59. Parsons, J.B., and Rock, C.O. (2013). Bacterial lipids: metabolism and membrane homeostasis. Prog. Lipid Res. 52, 249–276.

60. Miller, S.J., Aly, R., Shinefeld, H.R., and Elias, P.M. (1988). In vitro and in vivo antistaphylococcal activity of human stratum corneum lipids. Arch. Dermatol. 124, 209–215.

61. Dayan, N., and Wertz, P.W. (2011). Innate Immune System of Skin and Oral Mucosa: Properties and Impact in Pharmaceutics, Cosmetics, and Personal Care Products (John Wiley & Sons).

62. Fischer, C.L., Walters, K.S., Drake, D.R., Dawson, D.V., Blanchette, D.R., Brogden, K.A., and Wertz, P.W. (2013). Oral mucosal lipids are antibacterial against Porphyromonas gingivalis, induce ultrastructural damage, and alter bacterial lipid and protein compositions. Int. J. Oral Sci. 5, 130–140.

63. Zheng, C.J., Yoo, J.-S., Lee, T.-G., Cho, H.-Y., Kim, Y.-H., and Kim, W.-G. (2005). Fatty acid synthesis is a target for antibacterial activity of unsaturated fatty acids. FEBS Lett. 579, 5157–5162.

64. Alnaseri, H., Arsic, B., Schneider, J.E.T., Kaiser, J.C., Scinocca, Z.C., Heinrichs, D.E., and McGavin, M.J. (2015). Inducible Expression of a Resistance-Nodulation-Division-Type Efflux Pump in Staphylococcus aureus Provides Resistance to Linoleic and Arachidonic Acids. J. Bacteriol. 197, 1893–1905.

65. Truong-Bolduc, Q.C., Villet, R.A., Estabrooks, Z.A., and Hooper, D.C. (2014). Native efflux pumps contribute resistance to antimicrobials of skin and the ability of Staphylococcus aureus to colonize skin. J. Infect. Dis. 209, 1485–1493.

66. Jerse, A.E., Sharma, N.D., Simms, A.N., Crow, E.T., Snyder, L.A., and Shafer, W.M. (2003). A gonococcal efflux pump system enhances bacterial survival in a female mouse model of genital tract infection. Infect. Immun. 71, 5576–5582.

67. Chen, Y.Y., Liang, N.Y., Curtis, J.M., and Gänzle, M.G. (2016). Characterization of Linoleate 10-Hydratase of Lactobacillus plantarum and Novel Antifungal Metabolites. Front. Microbiol. 7, 1561.

68. Williams, W.L., Broquist, H.P., and Snell, E.E. (1947). OLEIC ACID AND RELATED COMPOUNDS AS GROWTH FACTORS FOR LACTIC ACID BACTERIA. J. Biol. Chem. 170, 619–630.

69. Denou, E., Pridmore, R.D., Berger, B., Panoff, J.-M., Arigoni, F., and Brüssow, H. (2008). Identification of genes associated with the long-gut-persistence phenotype of the probiotic Lactobacillus johnsonii strain NCC533 using a combination of genomics and transcriptome analysis. J. Bacteriol. 190, 3161–3168.

70. Taga, M.E., and Ludington, W.B. (2023). Nutrient encryption and the diversity of cobamides, siderophores, and glycans. Trends Microbiol. 31, 115–119.

71. Kramer, J., Özkaya, Ö., and Kümmerli, R. (2020). Bacterial siderophores in community and host interactions. Nat. Rev. Microbiol. 18, 152–163.

72. Leinweber, A., Fredrik Inglis, R., and Kümmerli, R. (2017). Cheating fosters species co-existence in well-mixed bacterial communities. ISME J. 11, 1179–1188.

73. Degnan, P.H., Barry, N.A., Mok, K.C., Taga, M.E., and Goodman, A.L. (2014). Human gut microbes use multiple transporters to distinguish vitamin B₁₂ analogs and compete in the gut. Cell Host Microbe 15, 47–57.

74. Shelton, A.N., Seth, E.C., Mok, K.C., Han, A.W., Jackson, S.N., Haft, D.R., and Taga, M.E. (2019). Uneven distribution of cobamide biosynthesis and dependence in bacteria predicted by comparative genomics. ISME J. 13, 789–804.

75. Aukrust, T.W., Brurberg, M.B., and Nes, I.F. (1995). Transformation of Lactobacillus by electroporation. Methods Mol. Biol. 47, 201–208.

76. Aune, T.E.V., and Aachmann, F.L. (2010). Methodologies to increase the transformation efficiencies and the range of bacteria that can be transformed. Appl. Microbiol. Biotechnol. 85, 1301–1313.

77. Fristot, E., Bessede, T., Camacho Rufino, M., Mayonove, P., Chang, H.-J., Zuniga, A., Michon, A.-L., Godreuil, S., Bonnet, J., and Cambray, G. (2023). An optimized electrotransformation protocol for Lactobacillus jensenii. PLoS One 18, e0280935.

78. Zuo, F., Chen, S., and Marcotte, H. (2020). Engineer probiotic bifidobacteria for food and biomedical applications - Current status and future prospective. Biotechnol. Adv. 45, 107654.

79. Luchansky, J.B., Muriana, P.M., and Klaenhammer, T.R. (1988). Application of electroporation for transfer of plasmid DNA to Lactobacillus, Lactococcus, Leuconostoc, Listeria, Pediococcus, Bacillus, Staphylococcus, Enterococcus and Propionibacterium. Mol. Microbiol. 2, 637–646.

80. Rampersaud, R. (2014). Identifcation and Characterization of Inerolysin, the Cholesterol Dependent Cytolysin Produced by Lactobacillus Iners. 10.7916/D8P26W2V.

81. Herbst-Kralovetz, M.M., Pyles, R.B., Ratner, A.J., Sycuro, L.K., and Mitchell, C. (2016). New Systems for Studying Intercellular Interactions in Bacterial Vaginosis. J. Infect. Dis. 214 *Suppl 1*, S6–S13.

82. Miller, E.A., Beasley, D.E., Dunn, R.R., and Archie, E.A. (2016). Lactobacilli Dominance and Vaginal pH: Why Is the Human Vaginal Microbiome Unique? Front. Microbiol. 7, 1936.

83. Yildirim, S., Yeoman, C.J., Janga, S.C., Thomas, S.M., Ho, M., Leigh, S.R., Primate Microbiome Consortium, White, B.A., Wilson, B.A., and Stumpf, R.M. (2014). Primate vaginal microbiomes exhibit species specificity without universal Lactobacillus dominance. ISME J. 8, 2431–2444.

84. Wolfarth, A.A., Smith, T.M., VanInsberghe, D., Dunlop, A.L., Neish, A.S., Corwin, E.J., and Jones, R.M. (2020). A Human Microbiota-Associated Murine Model for Assessing the Impact of the Vaginal Microbiota on Pregnancy Outcomes. Front. Cell. Infect. Microbiol. 10, 570025.

85. Huerta-Cepas, J., Szklarczyk, D., Heller, D., Hernández-Plaza, A., Forslund, S.K., Cook, H., Mende, D.R., Letunic, I., Rattei, T., Jensen, L.J., et al. (2019). eggNOG 5.0: a hierarchical, functionally and phylogenetically annotated orthology resource based on 5090 organisms and 2502 viruses. Nucleic Acids Res. 47, D309–D314.

86. Cantalapiedra, C.P., Hernández-Plaza, A., Letunic, I., Bork, P., and Huerta-Cepas, J. (2021). eggNOG-mapper v2: Functional Annotation, Orthology Assignments, and Domain Prediction at the Metagenomic Scale. Mol. Biol. Evol. 38, 5825–5829.

87. Edgar, R.C. (2021). MUSCLE v5 enables improved estimates of phylogenetic tree confidence by ensemble bootstrapping. bioRxiv, 2021.06.20.449169. 10.1101/2021.06.20.449169.

88. Kozlov, A.M., Darriba, D., Flouri, T., Morel, B., and Stamatakis, A. (2019). RAxML-NG: a fast, scalable and user-friendly tool for maximum likelihood phylogenetic inference. Bioinformatics 35, 4453–4455.

89. Nugent, R.P., Krohn, M.A., and Hillier, S.L. (1991). Reliability of diagnosing bacterial vaginosis is improved by a standardized method of gram stain interpretation. J. Clin. Microbiol. 29, 297–301.

90. Dong, K.L., Moodley, A., Kwon, D.S., Ghebremichael, M.S., Dong, M., Ismail, N., Ndhlovu, Z.M., Mabuka, J.M., Muema, D.M., Pretorius, K., et al. (2018). Detection and treatment of Fiebig stage I HIV-1 infection in young at-risk women in South Africa: a prospective cohort study. Lancet HIV 5, e35–e44.

91. Shishkin, A.A., Giannoukos, G., Kucukural, A., Ciulla, D., Busby, M., Surka, C., Chen, J., Bhattacharyya, R.P., Rudy, R.F., Patel, M.M., et al. (2015). Simultaneous generation of many RNA-seq libraries in a single reaction. Nat. Methods 12, 323–325.

92. Zhu, Y.Y., Machleder, E.M., Chenchik, A., Li, R., and Siebert, P.D. (2001). Reverse transcriptase template switching: a SMART approach for full-length cDNA library construction. Biotechniques 30, 892–897.

93. Li, H., and Durbin, R. (2009). Fast and accurate short read alignment with Burrows-Wheeler transform. Bioinformatics 25, 1754–1760.

94. Leenhouts, K., Buist, G., Bolhuis, A., ten Berge, A., Kiel, J., Mierau, I., Dabrowska, M., Venema, G., and Kok, J. (1996). A general system for generating unlabelled gene replacements in bacterial chromosomes. Mol. Gen. Genet. 253, 217–224.

95. Russell, W.M., and Klaenhammer, T.R. (2001). Efficient system for directed integration into the Lactobacillus acidophilus and Lactobacillus gasseri chromosomes via homologous recombination. Appl. Environ. Microbiol. 67, 4361–4364.

96. Goh, Y.J., Azcárate-Peril, M.A., O’Flaherty, S., Durmaz, E., Valence, F., Jardin, J., Lortal, S., and Klaenhammer, T.R. (2009). Development and application of a upp-based counterselective gene replacement system for the study of the S-layer protein SlpX of Lactobacillus acidophilus NCFM. Appl. Environ. Microbiol. 75, 3093–3105.

97. Selle, K., Goh, Y.J., O’Flaherty, S., and Klaenhammer, T.R. (2014). Development of an integration mutagenesis system in Lactobacillus gasseri. Gut Microbes 5, 326–332.

98. Duong, T., Miller, M.J., Barrangou, R., Azcarate-Peril, M.A., and Klaenhammer, T.R. (2011). Construction of vectors for inducible and constitutive gene expression in Lactobacillus. Microb. Biotechnol. 4, 357–367.

99. Schneewind, O., and Missiakas, D. (2014). Genetic manipulation of Staphylococcus aureus. Curr. Protoc. Microbiol. 32, Unit 9C.3.

100. Novick, R.P. (1991). Genetic systems in staphylococci. Methods Enzymol. 204, 587–636.

101. Bligh, E. Graham, and W. Justin Dyer. (1959). A rapid method of total lipid extraction and purification. Canadian journal of biochemistry and physiology 37, 911–917.

102. Hoang, T., Toler, E., DeLong, K., Mafunda, N.A., Bloom, S.M., Zierden, H.C., Moench, T.R., Coleman, J.S., Hanes, J., Kwon, D.S., et al. (2020). The cervicovaginal mucus barrier to HIV-1 is diminished in bacterial vaginosis. PLoS Pathog. 16, e1008236.

103. Caporaso, J.G., Lauber, C.L., Walters, W.A., Berg-Lyons, D., Lozupone, C.A., Turnbaugh, P.J., Fierer, N., and Knight, R. (2011). Global patterns of 16S rRNA diversity at a depth of millions of sequences per sample. Proc. Natl. Acad. Sci. U. S. A. 108 *Suppl 1*, 4516–4522.

104. Caporaso, J.G., Kuczynski, J., Stombaugh, J., Bittinger, K., Bushman, F.D., Costello, E.K., Fierer, N., Peña, A.G., Goodrich, J.K., Gordon, J.I., et al. (2010). QIIME allows analysis of high-throughput community sequencing data. Nat. Methods 7, 335–336.

105. Callahan, B.J., McMurdie, P.J., Rosen, M.J., Han, A.W., Johnson, A.J.A., and Holmes, S.P. (2016). DADA2: High-resolution sample inference from Illumina amplicon data. Nat. Methods 13, 581–583.

106. McMurdie, P.J., and Holmes, S. (2013). phyloseq: an R package for reproducible interactive analysis and graphics of microbiome census data. PLoS One 8, e61217.

107. Darriba, D., Posada, D., Kozlov, A.M., Stamatakis, A., Morel, B., and Flouri, T. (2020). ModelTest-NG: A New and Scalable Tool for the Selection of DNA and Protein Evolutionary Models. Mol. Biol. Evol. 37, 291–294.

108. Edler, D., Klein, J., Antonelli, A., and Silvestro, D. (2021). raxmlGUI 2.0: A graphical interface and toolkit for phylogenetic analyses using RAxML. Methods Ecol. Evol. 12, 373– 377.

109. Price, M.N., Dehal, P.S., and Arkin, A.P. (2010). FastTree 2--approximately maximum-likelihood trees for large alignments. PLoS One 5, e9490.

110. Letunic, I., and Bork, P. (2021). Interactive Tree Of Life (iTOL) v5: an online tool for phylogenetic tree display and annotation. Nucleic Acids Res. 49, W293–W296.

